# A Substantia Innominata-midbrain Circuit Controls a General Aggressive Response

**DOI:** 10.1101/2020.04.22.047670

**Authors:** Zhenggang Zhu, Qingqing Ma, Lu Miao, Hongbin Yang, Lina Pan, Kaiyuan Li, Ling-Hui Zeng, Xiaoxing Zhang, Jintao Wu, Sijia Hao, Shen Lin, Xiulin Ma, Weihao Mai, Xiang Feng, Yizhe Hao, Li Sun, Shumin Duan, Yan-qin Yu

**Author notes:** These authors contributed equally (Zhenggang Zhu, Qingqing Ma, Lu Miao). Correspondence (D.S.) and (Y.Y.Q.).

## Abstract

While aggressive behaviors are universal and essential for survival, ‘uncontrollable’ and abnormal aggressive behaviors in animals or humans may have severe adverse consequences or social costs. Neural circuits regulating specific forms of aggression under defined conditions have been described, but how brain circuits govern a general aggressive response remains unknown. Here, we found that posterior substantia innominata (pSI) neurons responded to several aggression-provoking cues with the graded activity of differential dynamics, predicting the aggressive state and the topography of aggression in mice. Activation of pSI neurons projecting to the periaqueductal gray (PAG) increased aggressive arousal and robustly initiated/promoted all the types of aggressive behavior examined in an activity level-dependent manner. Inactivation of the pSI circuit largely blocked diverse aggressive behaviors but not mating. By encoding a general aggressive response, the pSI-PAG circuit universally drives multiple aggressive behaviors and may provide a potential target for alleviating human pathological aggression.

**In Brief:** Zhu et al. find that the posterior substantia innominata neurons universally drive aggressive behaviors by representing aggressive state and topography of aggression in mice.

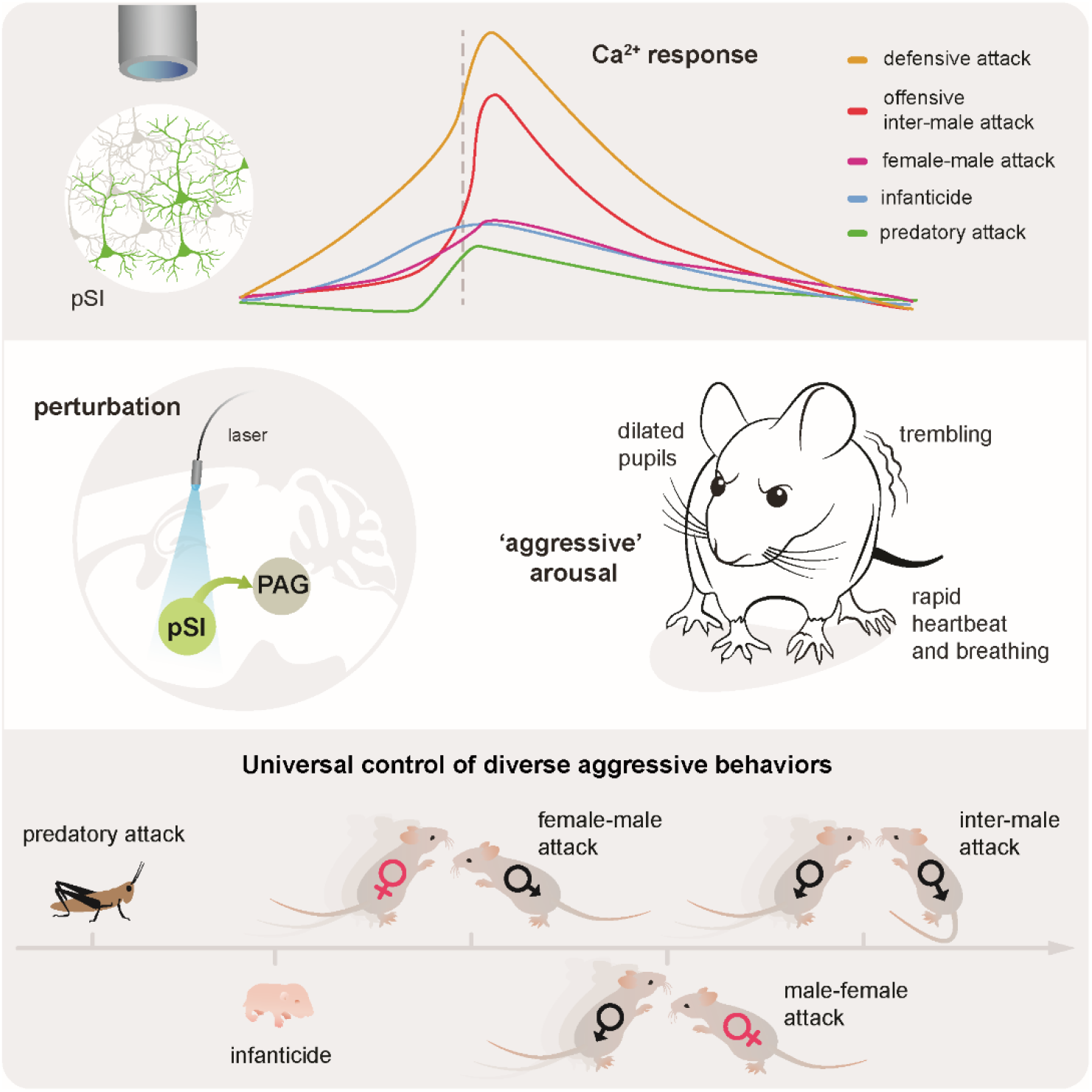

**Highlights:**

- pSI neuronal dynamics reflect aggressive state and the topography of aggression
- The pSI-PAG circuit promotes arousal and elicits thirteen aggressive behaviors
- The pSI controls various aggressive behaviors in an activity level-dependent manner
- Inactivation of the pSI circuit blocks diverse aggressive behaviors but not mating

## INTRODUCTION

Animals exhibit a range of aggressive behaviors (Moyer, 1968, Blanchard et al., 2003) essential for survival and reproduction (Nelson and Trainor, 2007), while pathological aggression causes serious social problems (Coccaro, 2012, Davidson et al., 2000). Distinct regions in the mouse brain are essential for male, female, predatory, and infant-directed aggressive behaviors (Lischinsky and Lin, 2020, Siegel and Victoroff, 2009). Furthermore, brain regions that have been examined for more than one type of aggressive behaviors suggest that distinct neural circuits might regulate aggression under specific internal and external conditions (Han et al., 2017, Yang et al., 2017, Hashikawa et al., 2017, Chen et al., 2019). For example, aggression initiated in the ventromedial hypothalamus (VMH) requires a specific genetic background, sexual-reproductive state, and social-contextual conditions (Yang et al., 2017, Lee et al., 2014, Hashikawa et al., 2017). On the other hand, social behavior-related conditions activated similar brain structures and overlapping circuits (Hashikawa et al., 2017, Kim et al., 2015, Lin et al., 2011, Dulac et al., 2014). Furthermore, stereotyped attack displays (e.g., biting, chasing, and aggressive arousal) are expressed in most aggressive behaviors (Hashikawa et al., 2018, Siegel and Victoroff, 2009), implying the involvement of some neural pathways for diverse aggressive behaviors. Besides, human pathological aggression is characterized by a variety of inappropriate and uncontrollable impulsive attacks (Coccaro, 2012) that may break out in the absence of an external threat, implying that some circuits in the brain may encode a general aggressive response. However, whether such synergistic circuits exist in the brain and how they regulate the internal state and behavioral activity patterns that together comprise diverse aggressive behaviors remain unclear.

The amygdala is a key node for regulating aggression-related emotional states (McCloskey et al., 2016, Davidson et al., 2000). However, the role of subdivisions of the amygdala in diverse aggressive behaviors and aggressive states remains elusive. The substantia innominata (SI), a sub-region of the basal forebrain that extends >2 mm rostrocaudally in the mouse brain (https://bbp.epfl.ch/nexus/cell-atlas/), can be divided into the anterior SI (aSI) and the posterior SI (pSI) (Figure S1), which has also been classified as the extended amygdala (Heimer et al., 1997, Grove, 1988a). Remarkably, the SI is also involved in the regulation of motivation and fear behaviors (Yu et al., 2017, Cui et al., 2017), the aggression-related emotional state.

Using electrophysiological recording and Ca^2+^ imaging, we found that mouse pSI neurons exhibited increased activity with differential intensity and dynamics. This graded neuronal activity was correlated with the biological significance (threatening/attack-provoking or neutral) of the different social-contextual cues and predicted the topography of diverse aggressive behaviors. Acute optogenetic or pharmacogenetic manipulation of pSI neurons projecting to PAG demonstrated that the pSI-PAG circuit universally controls diverse aggressive behaviors.

## RESULTS

### Posterior SI Neurons Are Highly Active During Inter-male Aggression

To explore the role of SI in mouse aggressive behaviors, we first examined the c-Fos expression in the SI and nearby regions induced by inter-male aggression (Figures 1A and S1). The number of c-Fos+ neurons in the pSI under aggressive conditions was ∼3.4-fold greater than that under the control investigating condition, while less of an increase occurred in the aSI, nearby central amygdala (CeA), and globus pallidus (GP) (Figures 1B, 1C).

**Figure 1.**
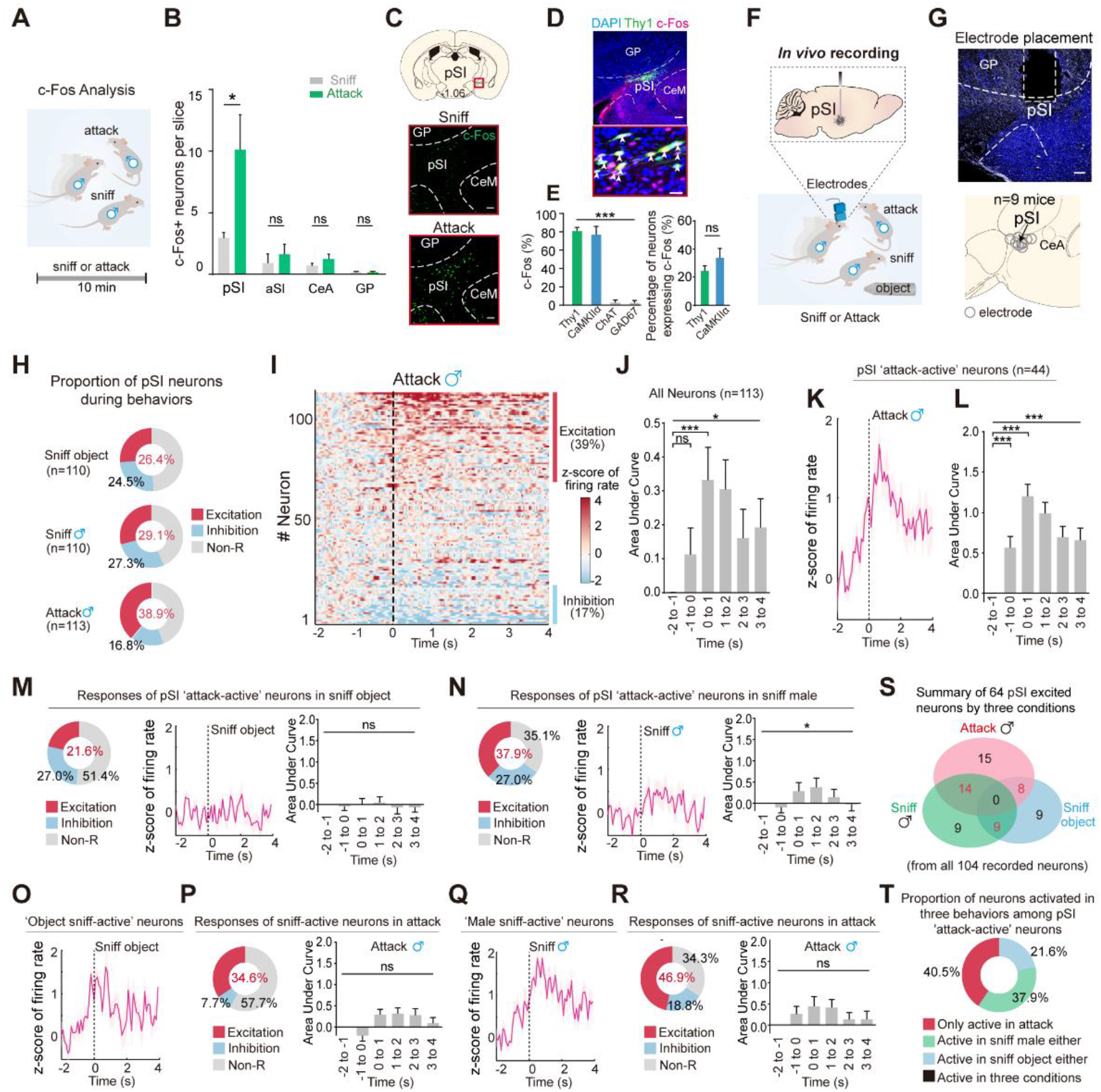
Elevated Neuronal Activity in the pSI during Inter-male Aggression. (A) Schematic of c-Fos analysis after sniff or attack. (B) c-Fos+ neurons per slice in the pSI, aSI, CeA, and GP. (C) Example images of c-Fos+ neurons (green) in the pSI after sniff (middle) or attack (bottom) (scale bars, 100 μm). CeM, medial division of CeA. (D) Upper, example image of c-Fos+ (magenta) and Thy1+ (green) neurons in the pSI after aggression (scale bar, 100 μm); Lower, magnified image (scale bar, 20 μm). (E) Left, percentage of c-Fos+ neurons co-localized with Thy1, CaMKIIα, ChAT, or GAD67 after an attack. Right, percentage of Thy1+ or CaMKIIα+ neurons expressing c-Fos after an attack. (F) Schematics of *in vivo* recording of pSI neuronal activity during social behaviors. (G) Representative image of electrode placement (upper, scale bar, 100 μm) and overlay of electrode placements in the pSI (lower). (H) Relative proportions of neurons excited (red), inhibited (blue), or unaffected (gray) while contacting an object, sniffing, or attacking a male mouse. (I) Heatmap of normalized firing rates of pSI neurons related to aggression. (J) Mean population activity (Area under the curve, AUC) before, during, and after an attack for all recorded neurons in all attacks examined. (K) Z-score of the firing rate of the neurons excited by aggression. (L) AUC per second before, during, and after an attack for neurons excited by aggression. (M-N) Relative proportions of ‘attack-active’ neurons excited (red), inhibited (blue), or unaffected (gray) while contacting an object (M) or a male (N) (left), Z-score of firing rate (middle) and AUC per second (right) of the ‘attack-active’ neurons excited by sniffing an object (M) or a male (N) for the ‘attack-active’ neurons. (O-R) Z-score of the firing rate of the ‘object sniff-active’ (O) or ‘male sniff-active’ (Q) neurons excited by sniffing an object (O) or a male (Q) (left). Relative proportions of ‘object sniff-active’ (P) or ‘male sniff-active’ (R) neurons excited (red), inhibited (blue), or unaffected (gray) by aggression (left), AUC per second before, during, and after an attack for ‘object sniff-active’ (P) or ‘male sniff-active’ (R) neurons (right). (S) Relative proportions of pSI neurons excited during three different behavioral activities. (T) The proportion of neurons excited in three distinct behaviors among pSI ‘attack-active’ neurons. Data are presented as the mean ± SEM. See also **Figure S1**.

We found that approximately 81% of the aggression-induced c-Fos+ neurons in the pSI were thymus cell antigen 1-positive (Thy1+), and 77% were calcium/calmodulin-dependent protein kinase IIα-positive (CaMKIIα+) (Figures 1D, 1E). We then made *in vivo* electrophysiological recordings of the pSI neuronal activity during non-aggressive and aggressive social behaviors (Figures 1F, 1G). The average firing rate of all the recorded pSI neurons (n = 113) was increased during aggression (Figures 1H-1J), with 39% showing increased activity during an attack (Figure 1H), which was much higher than that during a sniff (Figure 1H). Notably, activity in these aggression-excited pSI neurons started to increase during the sniff preceding an attack (Figures 1K, 1L). The pSI neurons excited with aggression were less activated when sniffing an object (22%, Figure 1M) than when sniffing a male mouse (38%, Figure 1 N). Moreover, we found that pSI neurons excited by sniffing a male, a provoking cue for aggression, were activated more in attacks than the excitation when sniffing an object (Figures 1O-1R; 47% *vs* 35%). Interestingly, none of the recorded neurons showed an increased firing rate under all three conditions (Figures 1S, 1T). In summary, we found that pSI neurons are highly active in inter-male aggression, and these neurons may encode the aggression and aggression-provoking cues.

### Differential Ca^2+^ Dynamics in pSI^Thy1^ Neurons Associated with the Different Aggressive States in Inter-male Aggression

To further explore the correlation between the activity in identified subtypes of pSI neurons and aggression, we injected viruses encoding GCaMP6m into the pSI of male Thy1-Cre mice (FVB/N) and used fiber photometry to record Ca^2+^ signaling in pSI^Thy1^ neurons during various social behaviors (Figures 2A, 2B, and S2A, S2B). Male Thy1-Cre mice exhibited robust aggressive behaviors (mean duration, 1.01 ± 0.06 s; Figures S2C, S2D). The activity of pSI^Thy1^ neurons started to increase before an attack (∼–1.2 s, pre-attack), peaked midway through the attack (duration of attacks, ∼1.0 s; peak magnitude (ΔF/F): 23.43%) and was sustained for some time after termination of the attack (∼4.8 s; post-attack, Figures S2E-S2H). Remarkably, distinct Ca^2+^ dynamics of pSI^Thy1^ neurons were found in attacks of different patterns or different episodes of continuous attacks (Figures S2I-S2P).

**Figure 2.**
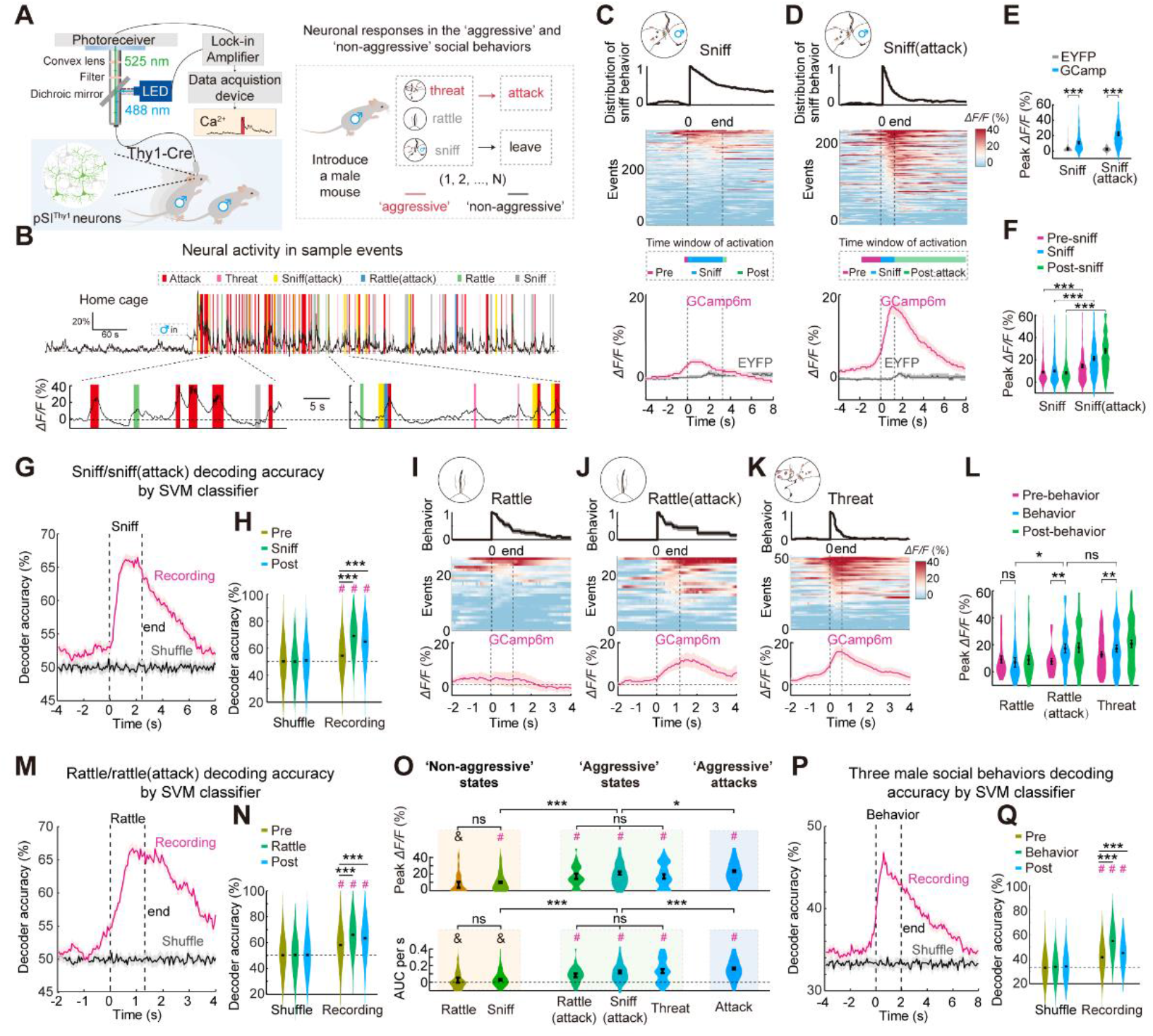
Graded Ca^2+^ Responses in pSI^Thy1^ Neurons under the Different Aggressive States. (A) Setup for fiber photometric recording in pSI^Thy1^ neurons during social behaviors. (B) Representative GCaMP6m signals during various social behaviors. (C, D) Distribution of episodes of behavior (upper), heatmaps of GCaMP6m Δ*F/F* signals from pSI^Thy1^ neurons, time windows of pSI^Thy1^ neuronal activation, and Δ*F/F* of the EYFP and GCaMP6m signals (lower) before, during, and after a sniff without an attack (C) and a sniff preceding an attack (D). (E) Peak Δ*F/F* of GCaMP6m and EYFP signals during a sniff with and without a subsequent attack. (F) Peak Δ*F/F* of GCaMP6m signals before, during, and after a sniff with or without a subsequent attack. (G,H) Sample decoding accuracy (G) and averaged sample decoding accuracy (H) of pSI^Thy1^ neuronal activity in the sniff/sniff (attack) trials under shuffled (black) and recording (red) conditions. (I-K) Distribution of episodes of behavior (upper), heatmaps of GCaMP6m Δ*F/F* signals from pSI^Thy1^ neurons (middle), and Δ*F/F* of GCaMP6m signals (lower) before, during, and after a rattle without an attack (I), a rattle preceding an attack (J), and threat behavior (K). (L) Peak Δ*F/F* of GCaMP6m signals before, during, and after a rattle without a subsequent attack, a rattle preceding an attack, and a threat behavior. (M,N) Decoding accuracy (M) and averaged sample decoding accuracy (N) of the pSI^Thy1^ neuronal activity in rattle/rattle (attack) trials. (O) Peak Δ*F/F* (upper) and AUC/s (lower) of GCaMP6m signals during non-aggressive states, aggressive states, and aggressive attacks. ‘#’ and ‘&’ above each bar indicate that the value during each behavior period was (*P* < 0.05) and wasn’t (*P* > 0.05) significantly changed compared with the EYFP control group. (P,Q) Decoding accuracy (P) and averaged sample decoding accuracy (Q) of the pSI^Thy1^ neuronal activity in three social behavior trials (sniff *vs* sniff (attack) *vs* attack). Data are presented as the mean ± SEM. See also **Figures S2, S3**.

Differential Ca^2+^ dynamics in the pSI^Thy1^ neurons were also recorded during other types of social behaviors such as sniffs and rattles, depending on whether these behaviors were followed by attack behavior (Figures 2A, 2B). Ca^2+^ activity in the pSI^Thy1^ neurons increased more during the sniff/rattle preceding an attack (peak Δ*F/F*: 21.23%/17.37%) than the simple sniff/rattle without an attack (9.90%/7.03%; Figures 2C-2F, and 2I-2J). Ca^2+^ levels associated with these sniffs or rattles predicted whether or not an actual attack would follow (Figures 2G-2H and 2M-2N). When mice threatened male intruders, the pSI^Thy1^ neuronal activity also increased to a high level (17.06%), similar to that in ‘aggressive’ rattles (Figures 2K, 2L).

Collectively, pSI^Thy1^ neurons showed a graded increase in Ca^2+^ activity in various inter-male social behaviors, with the least increase during ‘non-aggressive’ social behaviors such as a simple sniff or rattle, relatively high activity during ‘aggressive’ inter-male social behaviors such as sniff-attack or rattle-attack, and the highest activity in inter-male attacks (Figures 2O and S2). These behaviors in different aggressive states were largely predicted by the associated pSI^Thy1^ neuronal activity (Figures 2P, 2Q, and S2).

### Graded Ca^2+^ Activity in pSI^Thy1^ Neurons Associated with Various Aggression-provoking Cues and Different Aggressive Behaviors

Next, we investigated whether aggressive behaviors in male Thy1-Cre mice could be evoked by different social-contextual cues (Figure S4A) and how pSI^Thy1^ neurons responded to various cues that may or may not trigger aggression (Figure S4E). We found that various imminent non-social threat cues but not neutral or appetitive stimuli significantly activated the pSI^Thy1^ neurons (Figures S4F-S4I) and moderately evoked aggressive behaviors (Figures S4B-S4D). Moreover, introducing a conspecific into the home cage of a tested male mouse (Figure S4C) frequently evoked aggressive behaviors and immediately induced even higher activity in pSI^Thy1^ neurons than non-threat social cues (Figures S4J-S4N). Interestingly, the introduction of different types of the conspecific evoked differential intensity of pSI ^Thy1^ neuronal activity in the order (from weak to strong) female C57 < male C57 < male CD1 < male C57 with a subsequent attack (Figure S4O).

We next investigated whether and how Ca^2+^ activity in pSI^Thy1^ neurons correlates with different types of aggressive behavior (Figure 3A). First, we recorded the neural activity during male-female attacks when Thy1-Cre male mice occasionally attacked female mice (observed in 22.7% of mice, Figure 3B). We found that pSI^Thy1^ neuronal activity was increased during a male-female attack with a time window from –1.7 s to 7.5 s (Figure 3C) and peak magnitude similar to that in inter-male attacks (25.62% *vs* 23.43%, Figures 3D and S2F, S2G). When introduced to the home-cage of male CD-1 mice and attacked by the latter, the Thy1-Cre mice in turn exhibited defensive attacks. Interestingly, the increased neural activity during inter-male defensive attacks (peak Δ*F/F*: 30.23%) was even higher than that during inter-male offensive attacks (Figures 3E-3G). In contrast, although the pSI^Thy1^ neuronal activity was also increased in predatory attacks on crickets, the increase was much less and shorter in sustained time (from –0.2 s to 3.3 s) than that in inter-male attacks (Figures 3H-3L). The increased pSI^Thy1^ neuronal activity in pup-directed attacks had a similar magnitude but a longer time window (from –2.1 s to 4.8 s) than that in a predatory attack (Figures 3M-3Q). The female aggression in virgin female mice toward a male mouse, although seldom observed and lasting for a shorter time (duration in females *vs* males: 0.42 s *vs* 1.01 s) (Figures 3T, 3V and S2E), was associated with increased pSI^Thy1^ neuronal activity at a similar amplitude but an even longer time window (from –4.0 s to 7.5 s) than with predatory or pup-directed aggression (Figures 3R-3V). The pSI^Thy1^ neuronal Ca^2+^ properties in cricket-directed, pup-directed, and female social behaviors were used to nicely classified the sniffs, sniffs (attack), and attacks (Figures 3K, 3P, 3U) and predicted whether or not an actual attack would follow (Figures S3L-S3AA). Interestingly, the decoding accuracy in the 4 types of attack and approach (attack) was much higher than those in the 4 types of approach (Figures S3X, S3Y). Moreover, pSI^Thy1^ neurons in the tested mice showed graded increases in activity in the order (from weak to strong) predatory aggression < pup-directed aggression < female aggression < male offensive aggression < male defensive aggression (Figures S3Z, S3AA).

**Figure 3.**
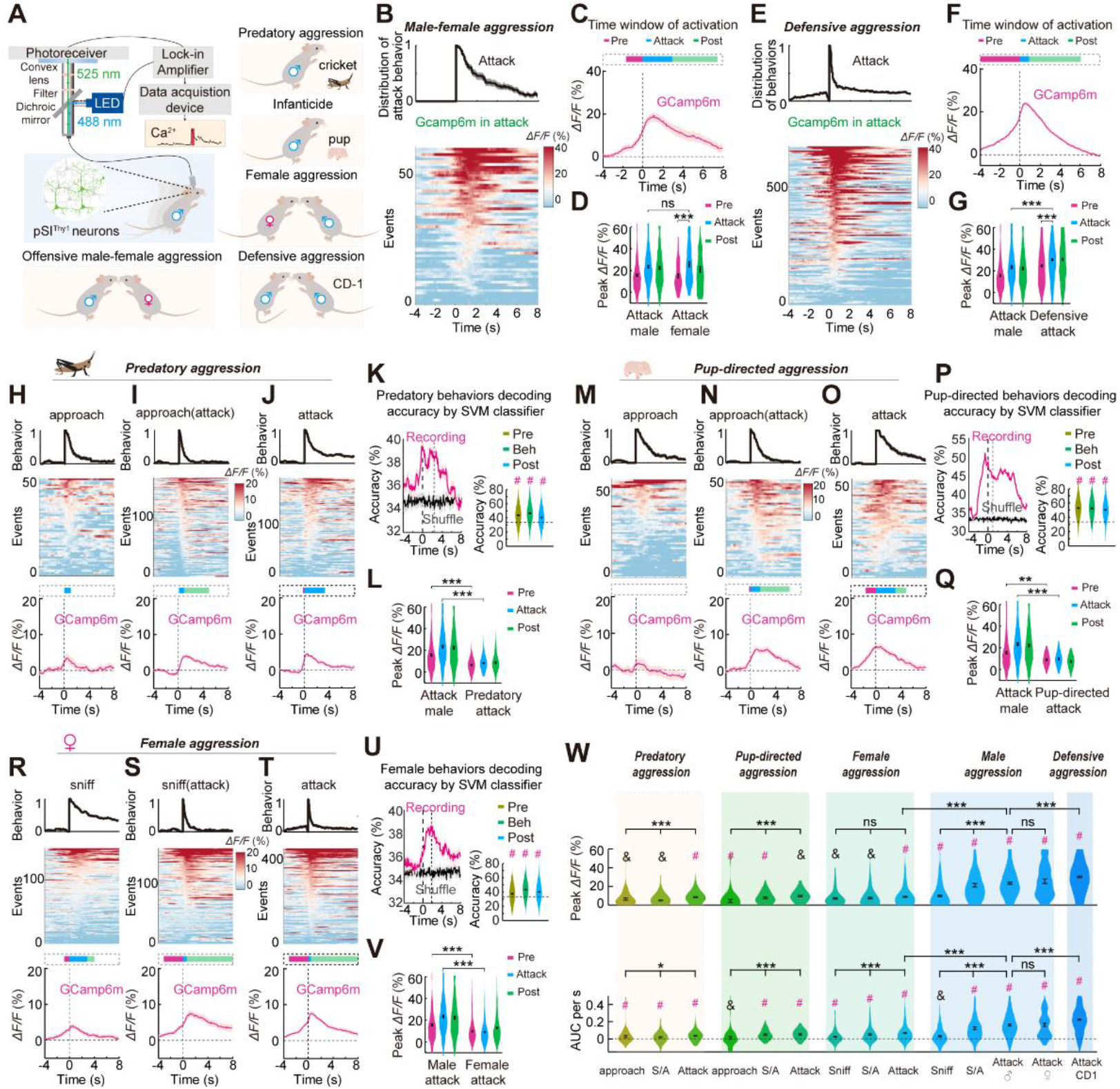
Graded Ca^2+^ Responses in pSI^Thy1^ Neurons under Diverse Aggressive Behaviors. (A) Setup for fiber photometric recording of pSI^Thy1^ neuronal activity during diverse aggressive behaviors. (B-G) Distribution of attack episodes (upper) and heatmap of GCaMP6m Δ*F/F* signals from pSI neurons (lower) in male-female (B) or defensive aggression (E). Time window of the pSI^Thy1^ neuronal activation (upper) and Δ*F/F* of GCaMP6m signals (lower) before, during, and after a male-female (C) or a defensive attack (F). Peak Δ*F/F* of GCaMP6m signals before, during, and after male-male and male-female attacks (D), or male-male offensive and defensive attacks (G). (H-V) From top to bottom: distribution of episodes of behavior, heatmaps of GCaMP6m Δ*F/F* signals from pSI neurons, the time window of pSI^Thy1^ neuronal activation, and Δ*F/F* of GCaMP6m signals of approach without an attack (H), approach preceding attack (I), and attack (J) in predatory aggression, without attack (M), approach preceding attack (N), and attack (O) in pup-directed aggression, or without attack (R), sniff preceding attack (S), and attack (T) in female-male aggression. Decoding accuracy of pSI neuronal Ca^2+^ signals in the approach/sniff, approach/sniff (attack), and attack trials (left); averaged sample decoding accuracy of pSI neuronal activity under recording conditions (right) for predatory attack (K), pup-directed attacks (P), or female-male attacks (U). Peak Δ*F/F* of GCaMP6m signals of a male-male attack and a predatory attack (L), male-male attacks and pup-directed attacks (Q), or male-male and female-male attacks (V). (W) Peak Δ*F/F* (upper) and AUC/s (lower) of GCaMP6m signals during diverse aggressive behaviors. Data are presented as the mean ± SEM. See also **Figures S3, S4**.

In summary, the perception of various aggression-provoking cues and the topography of aggressive behaviors are critical elements of the magnitude and dynamics of pSI^Thy1^ neuronal activity.

### Activation of pSI Neurons Drives Inter-male Aggression and Enhances Autonomic Arousal

We then asked whether the activation of pSI^Thy1^ neurons is sufficient to elicit aggression. The hM3Dq-encoding virus injected male mice were intraperitoneally injected (i.p.) with clozapine-N-oxide (CNO) to pharmacogenetically activate the pSI^Thy1^ neurons (Figures 4A, 4B). CNO treatment robustly induced aggressive behaviors in mice (Figures 4C and S5S). Interestingly, no mating behavior was observed (Figure 4C). Using the anterograde trans-synaptic tracing approach, we found that ventral lateral/lateral PAG (VL/LPAG), a region involved in the motor control of social behaviors (Behbehani, 1995), was densely innervated by pSI neurons (Figures S5A-S5D) (also see http://www.brain-map.org experiment id: 120281646) (Grove, 1988a). We further labeled the input neurons to the PAG by injecting a retrograde AAV-Retro-EGFP virus into the VL/LPAG (Figure 4D). A dense infection of upstream neurons was found in the pSI, but not or few in the aSI, GP, or MeA (Figures 4D, 4E). Most of the pSI neurons projecting to the PAG (pSI^-PAG^ neurons) were Thy1-positive or CaMKIIα-positive (∼87%/77%, Figures S5E-S5H), and chemogenetic activation of the pSI^-PAG^ neurons (Figures S5I-S5O) robustly elicited inter-male aggression, but no mating behavior was observed (Figures S5P-S5R).

**Figure 4.**
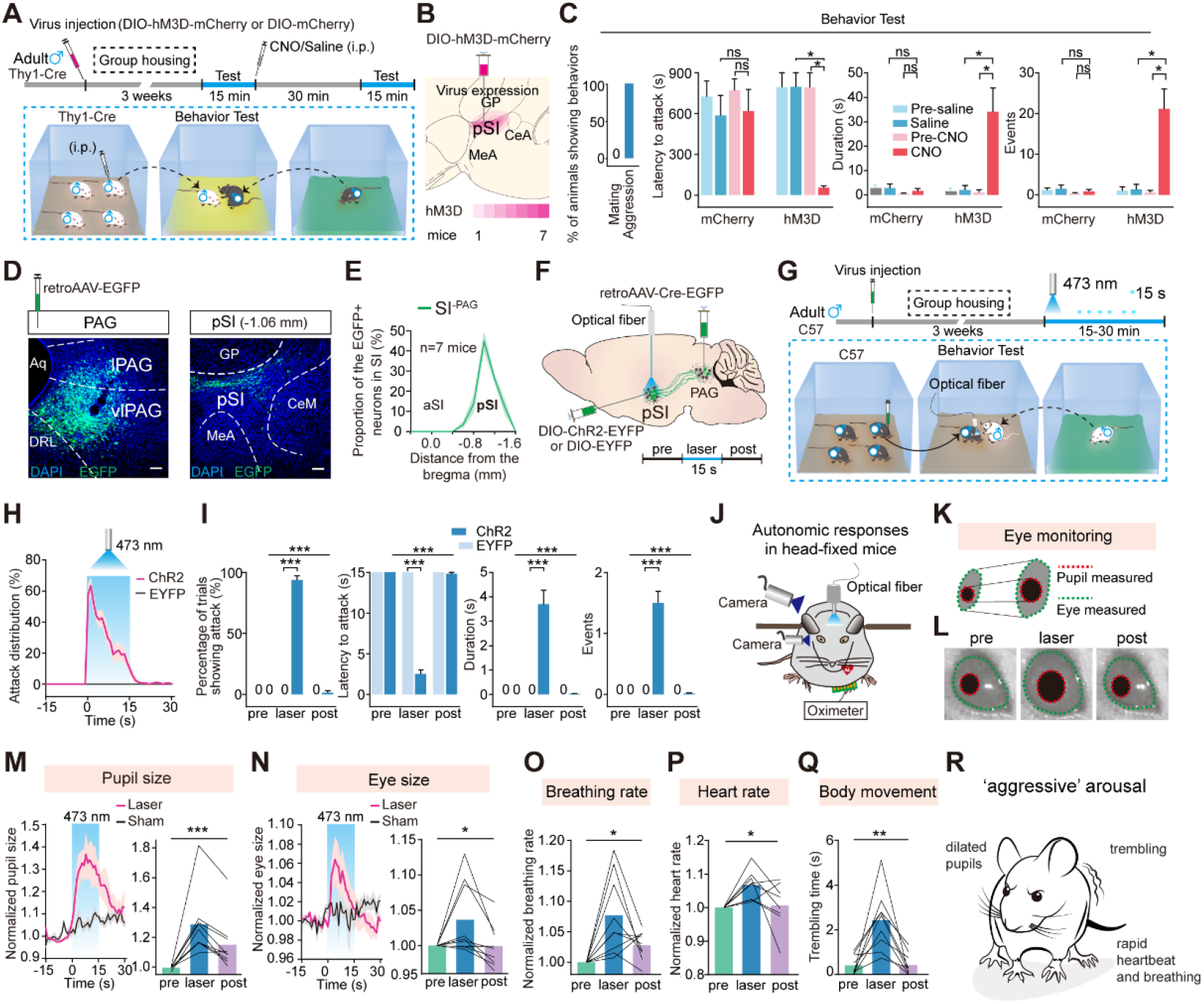
Activation of pSI^Thy1^ or pSI^-PAG^ Neurons Promotes Inter-male Aggression and Autonomic Arousal. (A) Experimental design and schematic of the viral injection strategy. (B) Overlay of DIO-hM3Dq-mCherry expression in pSI. (C) Percentage of animals showing aggressive or mating behaviors (left), latency, duration, and events of attacks before (pre-) and after injection with saline or CNO in the control and hM3Dq groups (right). (D) Representative images of retrograde AAV-retro-EGFP infection (green) in the PAG (left) and pSI (right) (scale bars, 100 μm). Aq, aqueduct; DRL, dorsal raphe nucleus, lateral part. (E) Proportion of retrogradely-labeled SI^-PAG^ neurons distributed in the SI. (F) Strategy for photoactivation of pSI^-PAG^ neurons in male mice. (G) The behavioral paradigm of aggression test. (H) Distribution of attack episodes during photostimulation. (I) Percentage of trials, latency, duration, and events of attacks in the ChR2 and control groups in pre-laser, laser, and post-laser phases. (J) Schematic of assessment of autonomic arousal in head-fixed mice with photostimulation of pSI^-PAG^ neurons. (K) Schematic for eye and pupil size measurements. (L) Representative images of the computer-detected pupil (red circle) and eye (green outline) before, during, and after photostimulation. (M, N) Left, changes of normalized pupil size (M) and eye size (N) in the laser and sham stimulation groups. Right, averaged pupil size (M) and eye size (N) before, during, and after photostimulation. (O-Q) Normalized breathing rate (O), normalized heart rate (P), and duration of body trembling (Q) before, during, and after photostimulation. (R) Cartoon of the increased aggressive arousal during the activation of pSI^-PAG^ neurons. Data are presented as the mean ± SEM. See also **Figures S5**, **S6**, **S7**, and **Movies S1**, **S2**.

We next used an optogenetic approach to establish a causal relationship between pSI^-PAG^ neuronal activity and aggressive behaviors with more precise temporal resolution (Figures 4F and S6A-S6C). A socially-housed male that exhibited minimal natural inter-male aggression (Figure 4G) immediately attacked a singly-housed male (Figures 4H and S6G; Movie S1) when pSI^-PAG^ neurons were optogenetically activated (Figures S6D-S6F). Notably, the light-evoked attack was characterized by biting an opponent’s dorsal surface at a mean latency of ∼3 s (Figure 4I), resembling the most violent natural aggressive behavior (Figure S2). Optogenetic activation of pSI^CaMKIIα^ neurons (Figures S6P-S6T) or PAG projecting pSI^Thy1^ neurons (Figure S7) similarly increased male aggression.

Diverse types of aggression may be primarily classified into proactive and affective forms, characterized by hypoarousal and hyperarousal, respectively (Siegel and Victoroff, 2009). We next determined whether the photoactivation of pSI^-PAG^ neurons modulates arousal (Figures 4J, 4K) (Wang et al., 2015). Photoactivation of the pSI^-PAG^ neurons resulted in a significant (∼30%) increase in pupil size (Figures 4L, 4M; Movie S2), a response that was sustained for >15 s after the stimulation ceased. The eyes were also wider during pSI stimulation (Figure 4N; Movie S2). Moreover, pSI^-PAG^ activation caused an increase in the breathing rate (Figure 4O) and heart rate (Figure 4P), as well as body trembling (Figure 4Q; Movie S2). However, the photostimulation of pSI^-PAG^ neurons did not affect the overall level of anxiety (Figures S6I, S6J), nor trigger the freezing or flight response (Figures S6K-S6O). In summary, optogenetic activation of pSI^-PAG^ neurons immediately induced robust attacks accompanied by an increased arousal state (Figure 4R), a response consistent with affective aggression (Siegel and Victoroff, 2009).

### Diverse Aggressive Behaviors Controlled by Differential Activity of pSI^-PAG^ Neurons

We reasoned that the increased arousal induced by pSI activation might establish a ‘heightened aggressive arousal’ that overcomes the internal and external requirements for evoking multiple types of aggression (Moyer, 1968). In the subsequent experiments, we determined whether the photo-activation of pSI^-PAG^ neurons induces a variety of aggressive behaviors under various conditions (Figure 5A).

**Figure 5.**
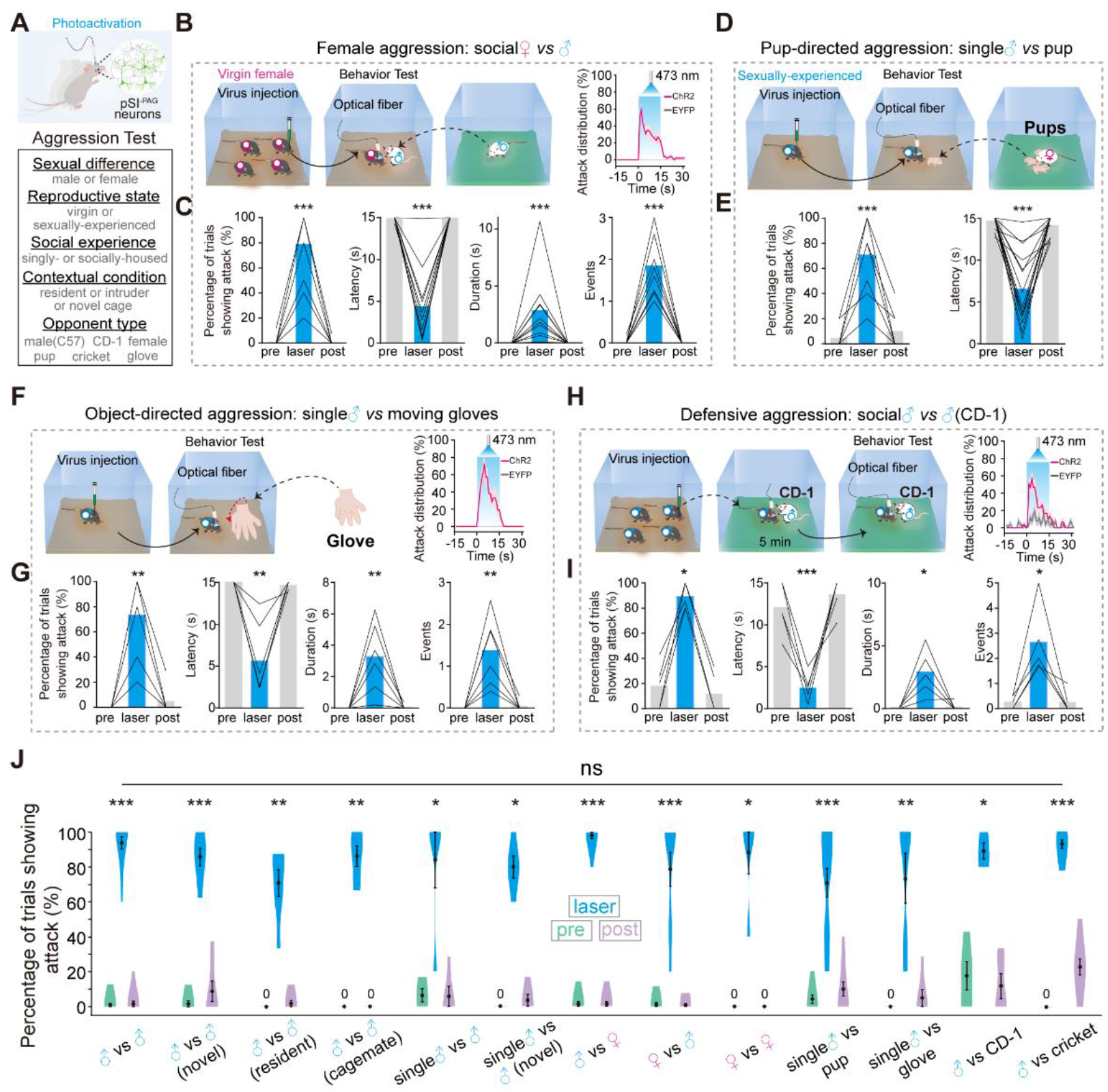
Activation of pSI^-PAG^ Neurons Promotes Diverse Aggressive Behaviors. (A) Schematic of the behavioral design to test aggression induced by photostimulation of pSI^-PAG^ neurons under various conditions. (B-I) Left, schematic of female (B), pup-directed (D), object-directed (F), and defensive aggression (H) with photostimulation of pSI^-PAG^ neurons. Right, distribution of attack episodes during photostimulation (B, F, H). Percentage of trials, latency, duration, and events of induced female (C), pup-directed (E, no duration and events), object-directed (G), and defensive aggression (I) by photostimulation of pSI^-PAG^ neurons. (J) Probability of light-induced aggression in thirteen aggression paradigms. Data are presented as the mean ± SEM. See also **Figures S8**, **S9**, **S10, S11**, and **Movies S3**, **S4**.

Offensive inter-male aggression is expressed distinctly when animals are under different social-contextual conditions (Yang et al., 2017, Zelikowsky et al., 2018). Surprisingly, photoactivation of pSI^-PAG^ neurons not only facilitated typical territorial inter-male aggression in both singly-housed (Figure 5J) and socially-housed (Figure 4) resident mice in their home cages but also in a novel cage (Figure 5J) or even when a socially-housed male encountered its male cage mate (Figure 5J). Notably, photoactivation of pSI^-PAG^ neurons even induced inter-male aggression when the optogenetically activated male mice intruded into the home cage of another mouse (Figure 5J). Under normal conditions, male mice do not attack females (Lin et al., 2011). Interestingly, photoactivation of pSI^-PAG^ neurons reliably induced male aggression against females to an extent similar to attacks against males (Figure 5J). Strikingly, pSI-activation in virgin females, which rarely express aggression, immediately initiated attacks against males (Figures 5B-5C; Movie S3) or females (Figure 5J).

We found that photoactivation of pSI^-PAG^ neurons in sexually-experienced males evoked aggression toward pups (Figures 5D, 5E), an infanticidal behavior usually committed by adult virgin males but not sexually-experienced males (Isogai et al., 2018). Photoactivation of pSI^-PAG^ neurons increased predatory attacks in male mice toward crickets (Figure 5J). Interestingly, photoactivation of pSI^-PAG^ neurons also induced object-directed aggression (Figures 5F, 5G): a moving glove evoked a rage-like response in male mice that never occurred in control mice (Lin et al., 2011). A socially-housed male that had been previously socially defeated by a more aggressive CD-1 male for 5 min would flee or freeze without apparent aggression (Blanchard et al., 2003) when encountering the same CD-1 mouse later in the home cage of that CD-1 mouse (Movie S4). Surprisingly, photoactivation of pSI^-PAG^ neurons in the defeated mouse immediately shifted the flight or freeze into a defensive attack towards the aggressive CD-1 male (Figures 5H-5I; Movie S4). No increased mating behavior was observed under the above conditions. Optogenetic or pharmacogenetic activation of pSI^CaMKIIα/Thy1^ neurons similarly increased the rarely-observed male or female aggression (Figure S8). Furthermore, photoactivation of the pSI^- PAG^ or pSI^CaMKIIα^ neuronal projection terminals in the PAG, but not in the other two downstream target areas related to aggression, bed nucleus of the stria terminalis (BNST) or VMH (Lin et al., 2011) (Figure S9), triggered diverse aggressive behaviors (Figure S10).

Notably, the extent of an attack was similar among the above thirteen types of aggressive behaviors (Figure 5J) induced by the photoactivation of pSI^-PAG^ neurons at the same intensity of laser stimulation (2.7 mW). Considering that graded neuronal responses of pSI neurons were recorded during natural aggressive behaviors (Figure 3), we investigated whether multiple aggressive behaviors could be elicited by different activities of pSI^-PAG^ neurons (Figure 6A). We found that the strength of photoactivation was strongly correlated with the attack probability or latency to attack onset among all aggressive behaviors tested (Figures 6B-6G). However, distinct dose-response curves, obtained by non-linear regression fit of photostimulation intensity with the probability and latency of attack, were created from these five types of evoked aggression (Figures 6F-6G). Low-intensity photostimulation (<0.2 mW) mostly initiated predatory and pup-directed aggression, with less evoked female aggression and the least triggered male aggression (Figures 6F-6H). As the stimulation intensity increased to intermediate light intensities (0.2 - 1 mW), female-male, male-female, and inter-male aggression were more frequently evoked (Figures 6F, 6G). Moreover, under conditions of high-intensity photostimulation (>1 mW), all five types of aggressive behaviors were triggered (Figures 6F-6G and 5J). Thus, diverse aggressive behaviors are differentially initiated by scalable activation of pSI^-PAG^ neurons, with the different threshold in the order: predatory aggression < pup-directed aggression < female aggression < inter-male aggression (Figure 6H). We next traced the upstream of pSI^-PAG^ neurons (Figure S11) and found that pSI^-PAG^ neurons receive direct innervations from many areas (such as VMH, MeA, BNST) involved in aggressive or social cue-induced behaviors (Lin et al., 2011). Interestingly, the threshold of aggressive behaviors induced by activation of pSI^-PAG^ neurons was elevated after the acute social defeat (Figure S12), suggesting the pSI^-PAG^ neurons can be modulated by social stress.

**Figure 6.**
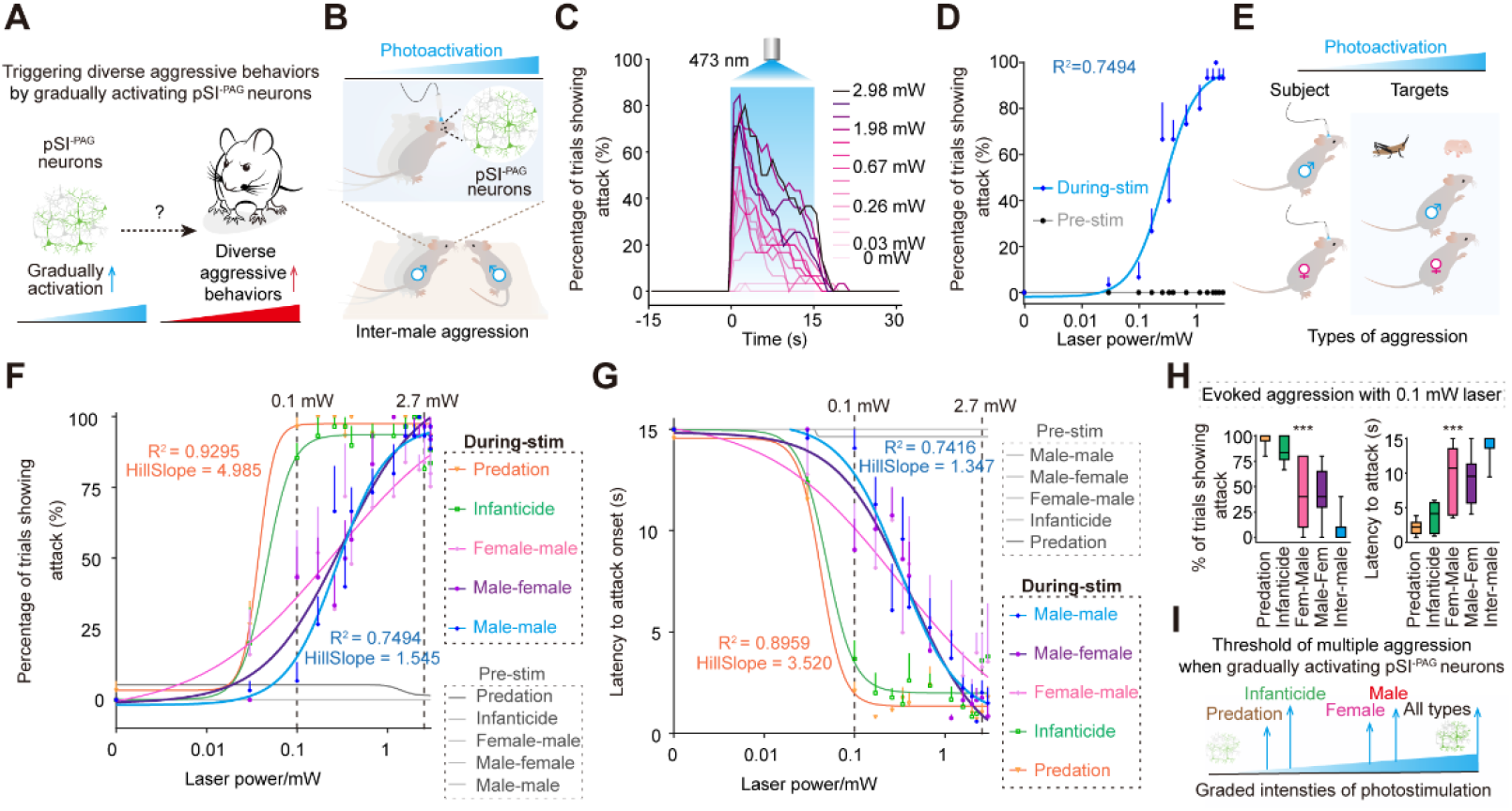
Intensity-Response Relationships of Photostimulation of pSI^-PAG^ Neurons and Success Rate of Evoked Attacks for Different Aggressive Behaviors. (A) Schematic for testing the relationship between the activation levels of pSI^-PAG^ neurons and different types of aggression. (B) Behavioral paradigm of the inter-male aggression test during photoactivation of pSI^-PAG^ neurons at different stimulation intensities. (C) Distribution of attack episodes during photoactivation of pSI^-PAG^ neurons at 13 graded stimulation intensities. (D) Nonlinear regression of the probabilities of evoked inter-male aggression with different intensities of photostimulation, before (gray) and during (blue) laser stimulation. Lines are non-linear fits (pooled across all mice and binned light intensities). (E) Behavioral paradigm of diverse aggressive behaviors during photoactivation of pSI^-PAG^ neurons at different stimulation intensities.| (F, G) Nonlinear regression of the probabilities (F) and latencies (G) of attacks for the five types of induced aggression with different photostimulation intensities before and during laser stimulation. (H) Probabilities of trials showing attack (left) and latency to attack onset (right) of the five types of induced aggression during laser stimulation with a lower intensity (0.1 mW). (I) Threshold model for the relationship between the activation level of pSI^-PAG^ neurons and evoked attacks for diverse aggressive behaviors. Data are presented as the mean ± SEM. See also **Figure S12.**

Together, the causal relationship of the graded pSI^-PAG^ neuronal activity with diverse aggressive behaviors, the dynamics (Figure 3), and anatomical evidence of pSI^-PAG^ neurons suggest that pSI activity represents a threshold variable for computing diverse aggressive behaviors.

### pSI Is Required for Diverse Aggressive Behaviors but not for Mating

We then asked whether the pSI is also necessary for these natural aggressive behaviors. We first optimized a behavioral paradigm in which a singly-housed male in its home cage robustly attacked a socially-housed male intruder (Figure 7A). Pharmacogenetically inhibiting the pSI^Thy1^ neurons (Figures 7B-7E and S13H-S13L) and the pSI^-PAG^ neurons (Figures S13A-S13G) largely blocked the aggression but not mating.

**Figure 7.**
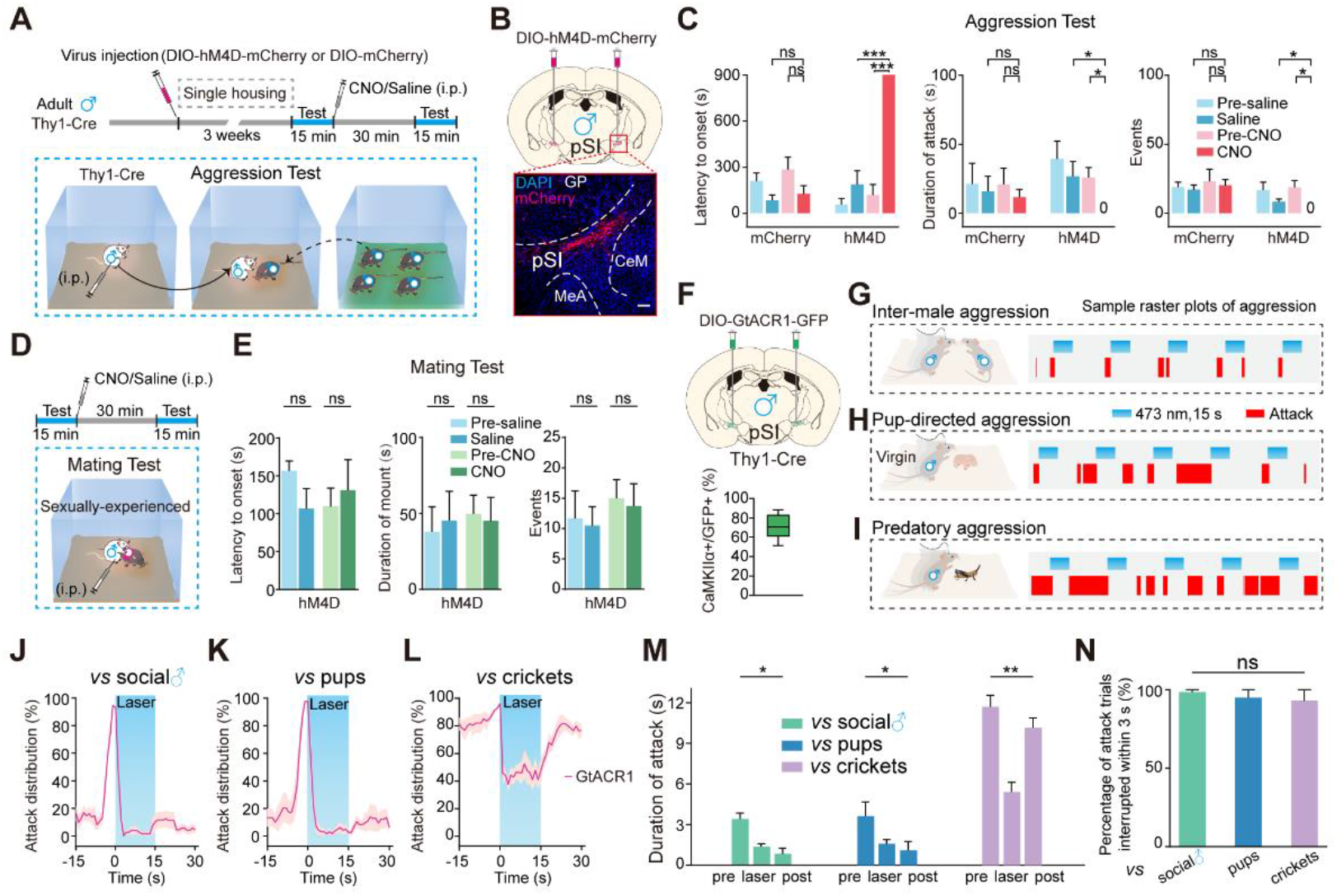
Inactivation of pSI^Thy1^ Neurons Interrupts Diverse Aggressive Behaviors but not Mating. (A) Experimental design and viral injection strategy for the expression of hM4Di-mCherry in pSI^Thy1^ neurons in male Thy1-Cre mice. (B) Schematic of virus injection and a representative histological image showing the expression of hM4Di-mCherry (red) in pSI^Thy1^ neurons (scale bar, 100 μm). (C) Latency (left), duration (middle), and events (right) of attack before and after injection of saline or CNO in the control and hM4Di groups. (D) Mating behavioral paradigm. (E) Latency to mounting onset, mounting duration, and mounting events before and after injection of saline or CNO in Thy1-Cre mice. (F) Upper, schematic of bilateral injection of DIO-GtACR1-GFP virus into the pSI in male Thy1-Cre mice. Lower, the co-localization ratio of GFP expression with CaMKIIα in the pSI. (G-I) Behavioral paradigms (left) and example raster plots (right) of optogenetic inhibition of pSI^Thy1^ neurons on offensive inter-male (G), pup-directed (H), and predatory aggression (I). (J-L) Distribution of attack episodes interrupted by optogenetic inhibition of pSI^Thy1^ neurons in inter-male (J), pup-directed (K), and predatory aggression (L). (M) Attack duration by optogenetic inhibition of aggressive behaviors. (N) Percentage of attack trials interrupted within 3 s during the laser phase by optogenetic inhibition in aggressive behaviors. Data are presented as the mean ± SEM. See also **Figures S13**, **S14**, and **Movie S5**.

Strikingly, photo-inhibition of pSI^Thy1^ neurons (Figures 7F, and S13H-S13J) by targeting the Cre-dependent expression of *Guillardia theta* anion channelrhodopsin 1 protein (GtACR1) immediately terminated inter-male offensive aggression when a singly-housed male in its home cage encountered a socially-housed male (Figures 7G, 7J, 7M; Movie S5). Optogenetically inhibiting pSI^Thy1^ neurons decreased the pup-directed aggression in singly-housed virgin adult males (Figures 7H, 7K, 7M). Similarly, photo-inhibition of pSI^Thy1^ neurons reduced the predatory aggression when a socially-housed male was attacking crickets (Figures 7I, 7L, 7M). The probability of interruption was similar among these aggressive behaviors blocked by pSI-inhibition (Figure 7N). Silencing pSI^Thy1^ neurons by photo-inhibition also significantly reduced the male-female and female-male aggression (Figures S14A-S14H), but running and feeding behaviors were not affected (Figures S14I-S14P).

## DISCUSSION

We have identified the pSI as a previously-unappreciated key center for regulating diverse aggressive behaviors in mice. Upon perception of various social-contextual cues, pSI neurons increased their activity with graded intensity and differential dynamics in a manner that nicely predicted and determined the states and topography of diverse aggressive behaviors.

### The pSI Circuit Controls a Scalable Aggressive State

Social behaviors like aggression are believed to be tightly associated with internal states (Anderson, 2016, Asahina et al., 2014, Zelikowsky et al., 2018). Several lines of evidence indicated that the pSI circuit controls a scalable aggressive state by integrating various attack-related social-contextual cues, and activity-dependently drives multiple aggressive behaviors. First, the activity of pSI^Thy1^ neurons increased in the following situations: 1. when mice were exposed to aggression-provoking aversive cues, but not with appetitive and neutral stimuli (Figure S4); 2. when mice actively contacted intraspecific and interspecific targets (sniff-attack and rattle-attack; Figures 2 and 3); and 3. before, during, and after the mice executed attacks (Figures 2, 3 and S2). Second, the initiation, termination, and topography of an attack were decoded by the pSI^Thy1^ neuronal activity (Figures 2, 3 and S2, S3). Third, the increased neuronal activity induced by various cues was graded in intensity and differed in dynamics, depending on the saliency of various attack-related cues, such as the ‘threat’ level of the cues (Figure S4) and the nature of the attacked targets (Figure 3). Finally, the activation of pSI^-PAG^ neurons not only triggered aggression but also increased autonomic arousal (Figure 4), an internal state correlated with aggressive behavior in flies and mice (Anderson, 2016, Asahina et al., 2014), defensive rage in cats (Siegel and Victoroff, 2009), and impulsive aggression in humans (Coccaro, 2012). An increased internal aggressive state may explain our finding that attacks induced by the activation of pSI neurons were directed at a variety of targets within the visual field, including a moving glove, crickets, and conspecific male or female mice (Figures 5, 6).

Notably, pSI shows a graded neuronal response that scales with multiple behavioral states rather than a ‘binary state’ (of the neurons that are only active during aggression and not in other contexts). This property may help the pSI to orchestrate other complex behaviors so as to decide whether attack or not to attack. Similar neural responses have been observed recorded in other behavioral state-defined neurons, such as ‘hunger’ neurons in the arcuate nucleus (Chen et al., 2015) and ‘parenting’ neurons in the preoptic area (Kohl et al., 2018).

### The pSI Circuit Governs Multiple Forms of Aggression

Various aggressive behaviors with specific displays are crucial ways to resolve conflicts for survival and reproduction and may have distinct neural circuitry mechanisms (Han et al., 2017, Yang et al., 2017, Hashikawa et al., 2017, Chen et al., 2019). For example, animals may respond with different levels of attacks when encountering a social threat, including displaying substantial offensive or defensive attack response initially and shifting aggression to defensive freezing or flight after the social defeat (Blanchard and Blanchard, 1988, Blanchard et al., 2003). We found that activation of pSI in a defeated mouse immediately shifted the defensive response to a robust attack pattern (Figure 5). Stronger stimulation of the pSI was required for an offensive attack after successive social defeats (Figure S12). These together highlight the conclusion that pSI neurons are critical for coping with a social threat by expressing attacks and can be modulated by past social stress.

VMH initiated-attacks in mice are affected by the singly- or socially-housing conditions (Yang et al., 2017), genetic background, and reproductive status (Hashikawa et al., 2017). Neurons in the central amygdala (Han et al., 2017) or zona incerta (Shang et al., 2019, Zhao et al., 2019) triggers predatory but not intra-conspecific aggression. Here, we demonstrate that the activation of pSI^-PAG^ neurons universally controls diverse aggressive behaviors, bypassing genetic background, reproductive state, social-contextual conditions, or opponent type required for aggression evoked by the activation of the dedicated circuits for controlling specific types of aggressive behaviors reported previously. It is plausible that the pSI-PAG circuit acts further downstream of the aggression-controlling pathway than the dedicated circuits for controlling specific attack types. Alternatively, pSI neurons may have a more global effect in the brain than other specifically-dedicated circuits to override the factors required for evoking aggressive behaviors.

### Identification of the pSI-Midbrain Circuit Specifically Responsible for Aggression

Aggression plays an essential role in survival and reproduction. Many reports are consistent with Tinbergen’s model (Tinbergen, 1951), which assigns aggression to a “reproductive” behavioral hierarchy (Lin et al., 2011, Anderson, 2012, Lee et al., 2014). For example, VMH (Lee et al., 2014) and MeA (Hong et al., 2014) neurons control several social behaviors, including attack, social investigation, and mounting in an intensity-dependent manner. Also, medial preoptic area neurons (Wu et al., 2014, Park et al., 2018) and MeA neurons (Chen et al., 2019) have been correlated with both aggression and parenting behaviors. Notably, we found that the pSI^-PAG^ neurons specifically regulate various aggressive behaviors but not mating, consistent with SI is much higher activated after aggression than after mating (Lin et al., 2011). Given the heterogeneous cell types in the pSI (http://mouse.brain-map.org/), it will be intriguing to know whether and how different populations of pSI neurons may encode opposing internal states of aggression and other social behaviors.

Many hypothalamic, amygdala, and midbrain regions directly connect with the pSI circuit (Figures S9 and S11) (Grove, 1988b, Grove, 1988a, Cui et al., 2017). For example, the pSI sends strong output to the aggression-related PAG and nearby BNST (Lin et al., 2011, Hashikawa et al., 2017, Mos et al., 1982, Padilla et al., 2016). Interestingly, our results indicate that the VL/LPAG, but not the BNST or VMH, is a downstream pathway through which the pSI can execute aggression and may function as a circuit dedicated to the finely-tuned, dynamically-controlled behavioral components of executing attack actions (Arber, 2012, Tovote et al., 2016, Falkner et al., 2020). It will be of great interest to explore whether the pSI directly receives and integrates convergent aggression-related information from these aggression-related circuits and sends out divergent signals to downstream pathways to execute these behaviors.

### Potential Relevance to Pathological Aggression

Abnormal aggressive arousal and an inability to control aggressive behaviors appropriately, such as “Intermittent Explosive Disorder” in humans, are serious social problems that still lack efficient interventional approaches (Siegel and Victoroff, 2009, Davidson et al., 2000). Various inappropriate aggressive behaviors induced by the pSI in mice could serve as an animal model of pathological aggression. Interestingly, abnormalities of the human amygdala are closely associated with pathological aggression (McCloskey et al., 2016, Nelson and Trainor, 2007, Coccaro, 2012, Davidson et al., 2000). As a conserved sub-region of the human amygdala (see http://atlas.brain-map.org/) and a necessary node for mouse aggressive behaviors, the pSI circuit may thus provide a therapeutic target for the suppression of human pathological aggression.

## ACKNOWLEDGMENTS

We thank Xiaohong Xu, Bo Li, Hailan Hu, Dayu Lin, and Nirao M Shah for discussions and reading the manuscript. We thank Hui-Fang Lou, Li-Ya Zhu, Xiaohong Xu, Hao-Ran Wang, Wei-Qian Jiang, and Hai-Shan Yao for technical support. This work was supported by the National Key Research and Development Program (2016YFA0501000 and 2016YFC1306700), the National Natural Science Foundation of China (81821091, 31771167, 31970939, and 81527901), the Non-profit Central Research Institute Fund of the Chinese Academy of Medical Sciences (2018PT31041), Science and Technology Planning Project of Guangdong Province (2018B030331001 and 2019B030335001), The Key Research and Development Program of Zhejiang Province (2020C03009), and Fundamental Research Funds for the Central Universities (2019FZA7009).

## AUTHOR CONTRIBUTIONS

Conceptualization, Z.G., Y.Y.Q., and D.S.; Methodology, Z.G., Y.Y.Q., and D.S.; Software, Z.G.; Formal Analysis, Z.G., M.L., Z.X., and M.X.; Investigation, Z.G., M.Q., M.L., P.L., L.K., W.J., Y.H., L.S., H.Y., M.W., F.X., L.S., and H.S.; Visualization, Z.G., M.Q., M.L., P.L., and L.K.; Writing Original Draft, Z.G., Y.Y.Q., and D.S.; Writing Review & Editing, Z.G., L. H.Z., Y.Y.Q., and D.S.; Funding Acquisition, Y.Y.Q., and D.S.; Resources, Y.Y.Q., and D.S.; Supervision, Z.G., Y.Y.Q., and D.S.

## DECLARATION OF INTEREST

The authors declare that they have no competing financial interests.

## STAR*METHODS

Detailed methods are provided in the online version of this paper and include the following:

- **CONTACT FOR REAGENT AND RESOURCE SHARING**
- **EXPERIMENTAL MODEL AND SUBJECT DETAILS**

- Animals
- **METHOD DETAILS**

- Viral injection and stereotaxic surgeries
- Anterograde and retrograde viral tracing
- Behavior measurements and video analysis
- Autonomic responses
- Anxiety and fear response measurements
- Feeding behavior measurements
- HM4Di-mediated neural silencing and hM3Dq-mediated neural activation
- ChR2-mediated neural activation
- ChR2-mediated neural activation before and after acute social defeat
- Optogenetic inhibition of neural activity
- Fiber photometry
- Decoder analysis by the support vector machine (SVM) algorithm
- Single-unit recording and data analysis
- *In vitro* electrophysiology
- Histology
- **QUANTIFICATION AND STASTISTICAL ANALYSIS**
- **KEY RESOURCES TABLE**

### CONTACT FOR REAGENT AND RESOURCE SHARING

Further information and requests for resources and reagents should be directed to and will be fulfilled by the Contact, Yan-qin Yu.

### KEY RESOURCES TABLE

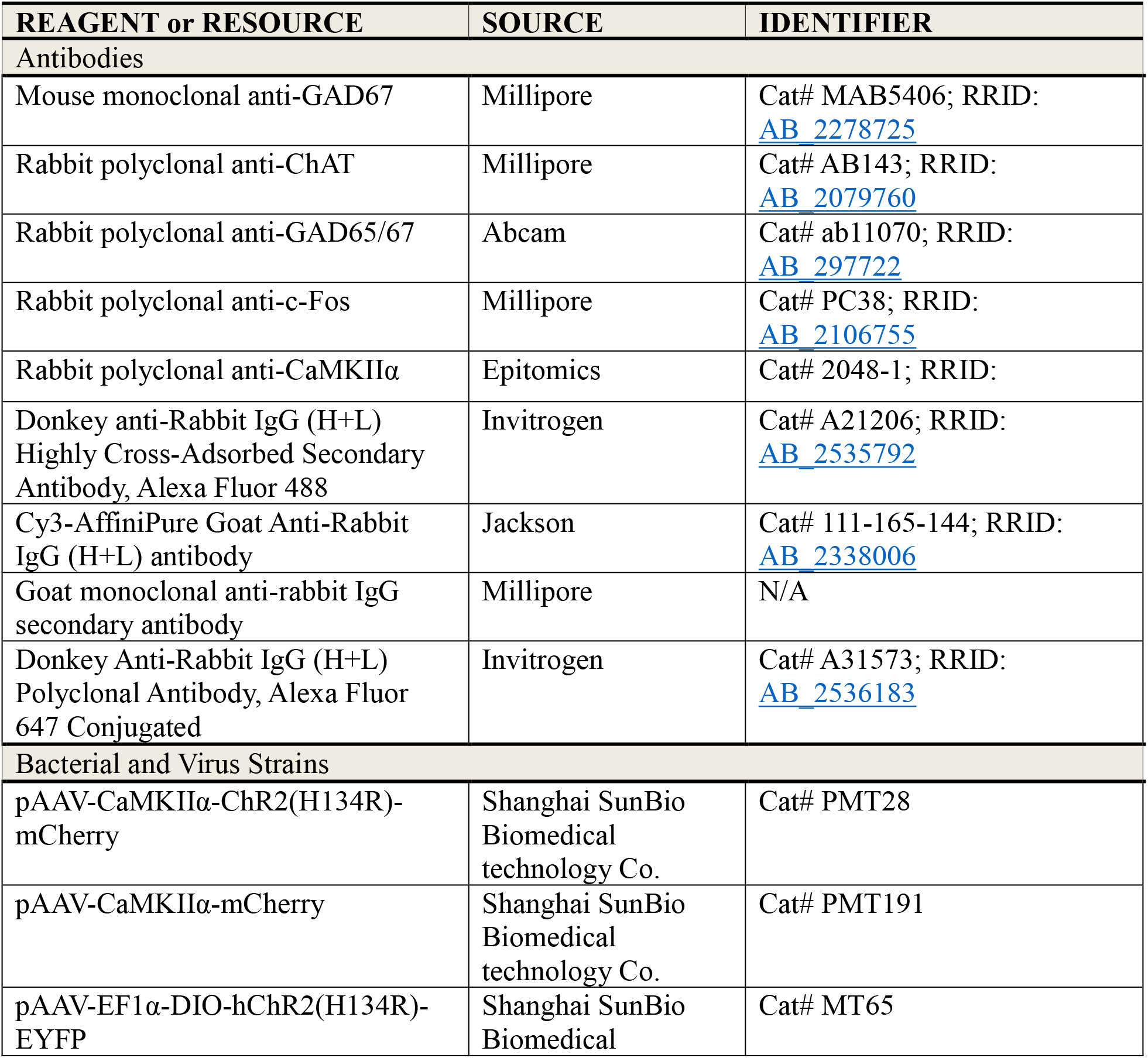
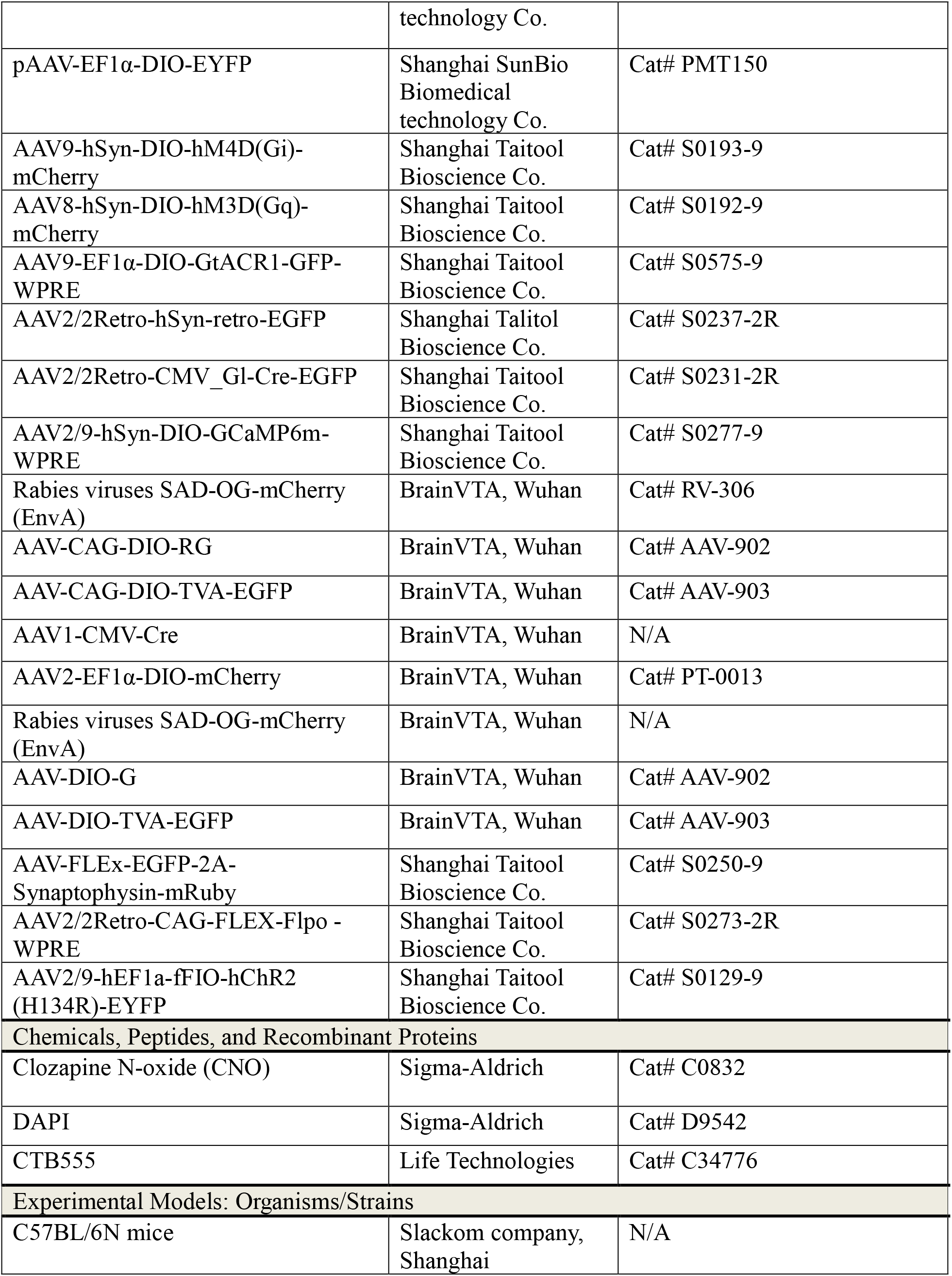
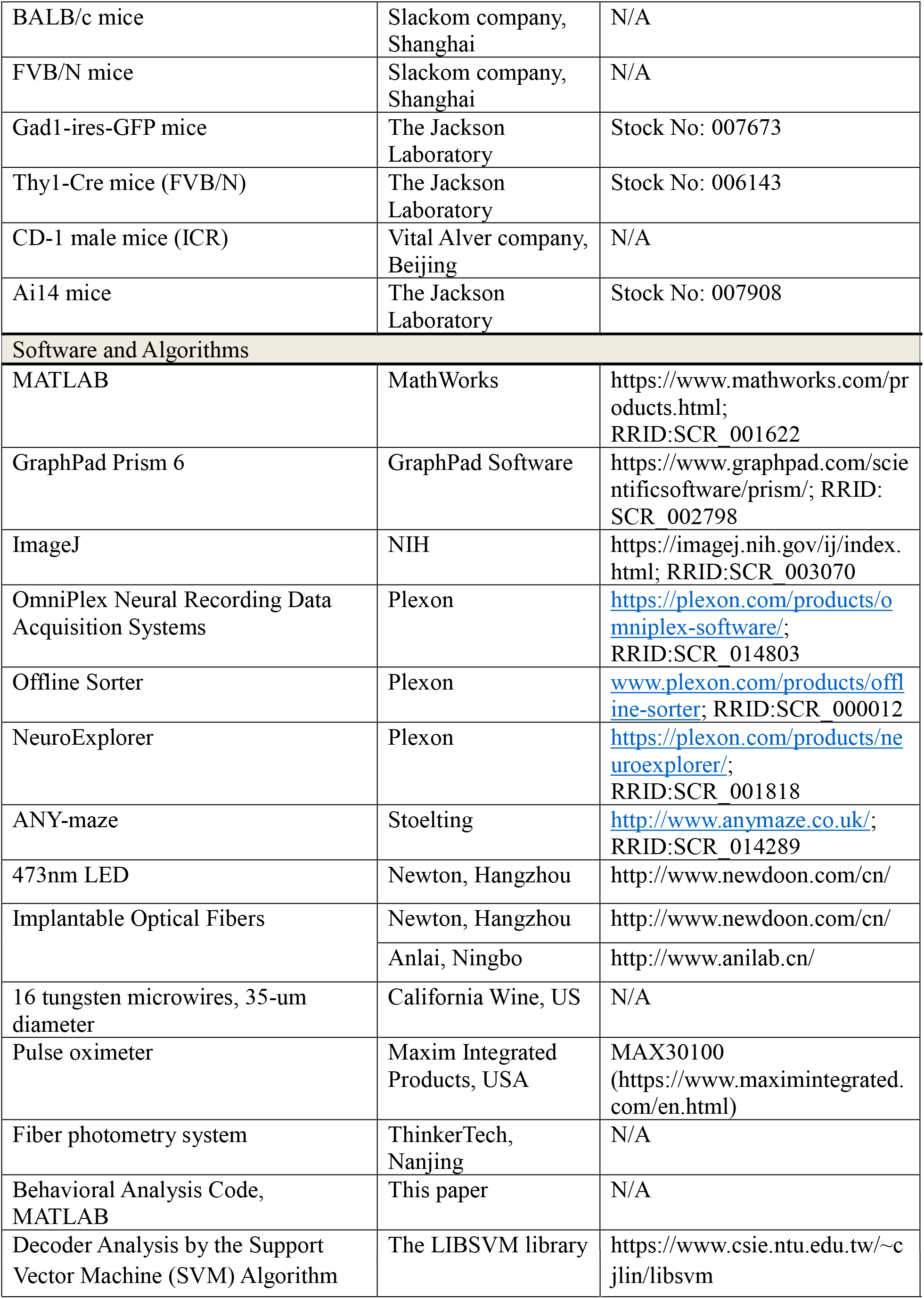

### EXPERIMENTAL MODEL AND SUBJECT DETAILS

#### Animals

Male GAD1 (GAD67)-GFP mice, male Ai14 mice, male CD-1 (ICR) mice, male FVB/N mice, Thy1-Cre (FVB/N), C57BL/6J, and BALB/c mice of both sexes were used. For behavior tests in wild type mice and functional manipulation using *in vivo* fiber photometry and *in vivo* multi-unit recording experiments, 8- to 20-week-old mice and 1- to 3-day-old naive pups were used. Mice were housed at 22 ± 1°C and 55 ± 5% humidity with food and water *ad libitum*. Animal experiments were conducted in accordance with the Guidelines for the Care and Use of Laboratory Animals of Zhejiang University and approved by Zhejiang University.

### METHOD DETAILS

#### Viral Injection and Stereotaxic Surgeries

AAV-CaMKIIα-hChR2 (H134R)-mCherry (AAV2/9, 7.57×10^12^ genomic copies/ml), AAV-CaMKIIα-mCherry (AAV2/9, 1.00×10^12^ genomic copies/ml), AAV-EF1α-DIO-ChR2 (H134R)-EYFP (AAV2/9, 2.42×10^12^ genomic copies/ml), AAV-EF1α-DIO-EYFP (AAV2/8, 2.50×10^12^ genomic copies/ml), AAV-hSyn-DIO-hM3Dq-mCherry (AAV2/9, 2.04×10^12^ genomic copies/ml), and AAV-hSyn-DIO-hM4Di-mCherry (AAV2/8, 1.13×10^12^ genomic copies/ml) were made by Shanghai SunBio Biomedical technology Co., Ltd. (Shanghai, China). AAV2/2Retro-CMV-Cre-EGFP (1.75×10^13^ genomic copies/ml), AAV2/2Retro-CMV-EGFP (2.28×10^13^ genomic copies/ml), AAV2/9-CAG-DIO-GCaMP6m-WPRE (1.00×10^13^ genomic copies/ml), and AAV-EF1α-DIO-GtACR1-P2A-GFP-WPRE (3.15×10^13^ genomic copies/ml) were made by Shanghai Taitool Bioscience Co., Ltd. (Shanghai, China). AAV1-CMV-Cre-WPRE-pA (1.40×10^13^ genomic copies/ml), and AAV-EF1α-DIO-mCherry (AAV2/9, 1.00×10^13^ genomic copies/ml) were provided by BrainVTA (Wuhan) Co., Ltd. (Wuhan, China).

Mice were anesthetized with 1% sodium pentobarbital before stereotaxic injection of a volume of 50-300 nl virus solution (depending on viral titer and expression strength). We injected 0.05-0.1 μl of the virus into each location at 0.01 μl/min, and the syringe was not removed until 5-10 min after the end of infusion to allow diffusion of the virus. A heating pad maintained core body temperature at 36°C. The coordinates of viral injection sites included the PAG (AP, –4.70 mm; ML, –0.8 mm; DV, –2.8 mm) and the pSI (AP, –0.90 mm; ML, –2.3 mm; DV, –4.5 mm). The coordinates of the cannula or optical fiber placement sites in the PAG and pSI were located 50-400 μm above the virus injection sites, and the coordinates of the optical fiber placement site in the BNST were AP, –0.22 mm; ML, –0.9 mm; DV, –3.5 mm, and the coordinates in the VMH were AP, –1.70 mm; ML, –0.8 mm; DV, –5.5 mm.

The pSI is a relatively little-studied region in the mouse brain, so we have better defined the pSI here (also see Figure S1), by comparing its relative coordinates and identifying the histological and chemical differences between the pSI and its surrounding structures (Grove, 1988a; Heimer et al., 1997). First, we registered the SI to the averaged template of the mouse brain atlas (Franklin and Paxinos, 2008) [we also referred to the Allen mouse brain atlas (http://atlas.brain-map.org/atlas) and https://bbp.epfl.ch/nexus/cell-atlas/)], and identified the pSI as the posterior part of the SI –0.7 mm to –1.6 mm from bregma. Second, the pSI lies deep to the globus pallidus (GP), lateral to the lateral hypothalamus (LH), and above the optic tract to the amygdala (dorsal to the MeA, dorsomedial to the CeA) (Grove, 1988a; Heimer et al., 1997). The boundary between the GP and pSI can be observed as differential contrast during confocal imaging, and the pSI is below the GP and internal capsule but within ∼300 μm. The optic tract is white matter, so the boundary between the optic tract and the pSI is even clearer. The MeA is immediately below the optic tract and the LH is above the optic tract, we separated the pSI with MeA, LH by comparing the relative position and sizes of the optic tract/GP. Moreover, we co-stained ChAT around the injection site in the virus injection and cannula implantation experiments and found strong ChAT expression in the virus-infected and cannula-implanted sites of the pSI. With these features and boundaries, relative coordinates, and histological differences, we confirmed that the virus injection, cannula implantation, and slice recording experiments were in the pSI and not in surrounding structures.

### Anterograde and Retrograde Viral Tracing

To reveal the outputs of the pSI, anterograde trans-synaptic AAV1-CMV-Cre virus (50 nl/side) was unilaterally injected into the pSI in male Ai14 mice (Zingg et al., 2017). To specifically label the connection from the pSI to the VL/LPAG, AAV1-CMV-Cre virus (50 nl/side) and AAV-DIO-mCherry virus (100 nl/side) was unilaterally injected into the pSI, and AAV-DIO-mCherry virus (100 nl/side) was unilaterally injected into the VL/LPAG in C57 mice. To retrogradely trace the input neurons to the VL/LPAG neurons, retrograde AAV2/2Retro-CMV-EGFP virus (60 nl/side) was used to unilaterally infect PAG neurons in C57 male mice (Tervo et al., 2016). The infection of upstream neurons from the VL/LPAG was examined in the aSI and pSI.

To retrogradely trace the monosynaptic presynaptic neurons to the pSI^-PAG^ neurons, we used a pseudorabies virus retrograde tracing strategy to trace the presynaptic targets of pSI^-PAG^ neurons in male mice. On day 1, we first injected AAV2/2-Retro-CMV-Cre-EGFP (60 nl/side) into the PAG to retrogradely express Cre in the pSI neurons. Then the Cre-dependent AAV carrying TVA, RG, SADΔG-EGFP (EnvA) and rabies virus (100 nl/side) was then injected into the pSI neurons at the same site to trace the upstream neurons. On the 15th day, the pseudorabies virus was injected at the same site in the pSI. We checked the range of viral infection in the pSI, and then examined the mCherry+ neurons in the whole brain of the male mice.

To specifically label the downstream targets from the pSI^-PAG^ neurons, retrograde AAV2/2Retro-CMV-EGFP virus (60 nl/side) was used to unilaterally infect PAG neurons in C57 male mice (Tervo et al., 2016). AAV-FLEx-EGFP-2A-Synaptophysin-mRuby (60 nl/side) was then unilaterally injected into the pSI, and the downstream targets of the pSI^-PAG^ neurons were examined by checking the Syp: mRuby+ presynaptic termini in the pSI and the regions in the whole brain areas.

AAV1-CMV-Cre virus (50 nl/side) and AAV-DIO-mCherry virus (100 nl/side) was unilaterally injected into the pSI, and AAV-DIO-mCherry virus (100 nl/side) was unilaterally injected into the VL/LPAG in C57 mice. To retrogradely trace the input neurons to the VL/LPAG neurons, retrograde AAV2/2Retro-CMV-EGFP virus (60 nl/side) was used to unilaterally infect PAG neurons in C57 male mice (Tervo et al., 2016). The infection of upstream neurons from the VL/LPAG was examined in the aSI and pSI.

#### Behavioral Measurements and Video Analysis

##### Aggression tests

Behavioral parameters such as the probability of trials with an induced attack, number of attack events, latency to the first attack, and duration of attack in animals with aggressive behaviors were analyzed in most behavioral experiments. Aggression parameters such as the percentage of interrupted attacks in trials, latency to attack offset, and duration of the attack were analyzed in the optogenetic inhibition experiments. The distribution of behavior episodes in Figures 2-7 was calculated as the percentage of trials showing the behavior at different time points, the y-axis of the plot means the percentage of trials showing this behavior. The features of aggression (Moyer, 1968) considered were: (a) various opponents for the tested mice: moving gloves, 1- to 3-day-old naive C57 pups (Wu et al., 2014, Isogai et al., 2018), crickets (Li et al., 2018, Han et al., 2017), and socially- and singly-housed adult C57 or BALB/c mice of both sexes (Yang et al., 2017, Hashikawa et al., 2017), and aggressive male CD-1 mice (see below for details for different forms of aggression); (b) various social-contextual conditions: tested mice were singly-housed for 3 days (Figures 4 and 5), singly-housed for 3 weeks (Figures 2-3 and 7), or socially-housed for at least 3 weeks before behavioral tests. During aggression tests, a tested mouse was introduced into the cage of another mouse, into a novel cage, or was resident in its home cage as described in the results; (c) both sexes and mice in different reproductive states were used (Hashikawa et al., 2017), that is, except for the sexually-experienced adult males used for the data are shown in Figures 5D-E and 7D-E, adult virgin males and females were used as tested mice; and (d) offensive or defensive aggression: offensive aggression was defined as biting, wrestling, tumbling, or chasing the conspecific, usually targeted toward the back and flanks of the opponent (Blanchard et al., 2003). All the above intra-conspecific attack conditions were considered as offensive aggression. A defensive attack was only considered when the mouse bit the snout of the conspecific or displayed upright postures (further details below).

##### Pup-directed aggression

One- to three-day-old naive C57 pups were used as standard pup intruders in all these behavior assays (Wu et al., 2014, Isogai et al., 2018). The pups were not related to the tested animals. A naive C57 pup was introduced into the home cage of each tested mouse and placed at the farthest corner from the resident’s resting nest. The pup was returned to its mother at the end of each assay. The probability of trials with an evoked attack and the latency to attack onset were scored in the optogenetic activation experiments (experiments were stopped if an attack caused actual wounds to the pup). In the optogenetic inactivation experiments and *in vivo* fiber photometry recording, the probability of trials with an interrupted attack within 3 s, latency to attack offset, and duration of attacks was scored.

##### Predatory aggression

House crickets (*Gryllus domesticus*) were purchased from pet food providers. Five to ten crickets per trial were used in the optogenetics experiments (Han et al., 2017, Li et al., 2018). In the neural activation experiments, the mouse was located in one corner of the cage and crickets were released near the diagonally-opposite corner. Mice fed *ad libitum* were placed in a clean cage and left for ∼10 min before laser stimulation. Behavioral parameters (percentage of trials with an induced attack, latency to first attack, attack events, and duration of attack) were analyzed only from animals displaying a predatory attack, such as chasing and biting the crickets. In the neural inhibition experiments and *in vivo* fiber photometry recording, mice were fasted for 24 h. The probability of trials with an interrupted attack within 3 s, latency to attack offset, and duration of attacks was scored.

##### Defensive aggression

Defensive aggression was defined as a reaction to prevent being bitten or threatening a conspecific (Blanchard et al., 2003). A C57 or Thy1-Cre mouse encountered an aggressive CD-1 male in a testing session. Behavioral parameters were analyzed only from animals with aggressive behaviors like upright postures and bites toward the snout. In the optogenetic activation experiments, a 5-min session was used to allow social interactions between the tested mouse and the CD-1 male. Since CD-1 mice are much stronger and more aggressive than the tested C57 or Thy1-Cre mice, tested mice would be socially defeated by CD-1 males during this session. Recording for defensive aggression by the tested mice was then started.

##### Male mating behavior

A sexually-receptive, adult, socially-housed, C57 female was introduced into the home cage of a sexually-experienced, singly-housed, Thy1-Cre male before and after the delivery of CNO or saline. Each test ran for 15 min and was videotaped and scored for latency to first mounting, mounting events, and duration of mounting (Yang et al., 2013, Yang et al., 2017).

All videos were recorded at 30 frames per second, and manual annotations were made by an observer blind to the experimental conditions. The annotations were then processed and analyzed using customized MATLAB programs to characterize and quantify behavioral episodes as previously described (Xu et al., 2012, Wei et al., 2018).

#### Autonomic Responses

**Heart rate and breathing rate measurements.** Heart rate was recorded in freely-moving mice using a pulse oximeter (MAX30100, Maxim Integrated, San Jose, CA). The abdominal area was shaved to enable the red light to pass through to the detector. Recordings were obtained for ∼10 min with intermittent blue laser stimulation in each mouse. Data were collected and analyzed using a custom-written MATLAB program. Breathing rate and episodes of bodily trembling were counted manually, based on chest movements captured on the video (Wang et al., 2015).

##### Pupil and eye measurement

Mice were adapted to constant, ambient room light for 1 h. Pupillary responses and eye size were measured under infrared light when the animals were head-fixed. The pupil and eye were video-recorded under constant light conditions before, during, and after photostimulation. A blue laser was interleaved and delivered to the pSI (10 times each). Control mice were treated the same except for sham stimulation. The size of the pupil and eye in each frame was determined using a custom-written MATLAB program and the normalized pupil diameter, eye outline, and relative pupil size were later measured and calculated (Wang et al., 2015).

#### Anxiety and Fear Response Measurements

##### Elevated plus maze test

The maze consisted of four arms arranged around a central platform (5 × 5 cm^2^) that allowed access to all arms. Mice were placed in the central platform facing the corner between a closed arm and an open arm, and were allowed to explore the maze for 5 min. The test was recorded with a video camera mounted above the maze and connected to a computer. The times spent in the open and closed arms were measured and automatically analyzed by ANY-maze (Stoelting Co., Wood Dale, IL).

##### Open-field test

Mice were placed in the center of a 40 × 40 × 40 cm polystyrene enclosure and videotaped individually. The center area was defined as the central 20 × 20 cm. To assess the effect of optogenetic activation of pSI neurons on anxiety and locomotor activity, mice were tested in 15-min sessions (consisting of 5-min light OFF, 5-min light ON, and 5-min light OFF periods). Blue laser light (473 nm, 5 ms, 40 Hz, 15 s) was delivered bilaterally during the light ON phase. To assess the immediate effect of optogenetic activation of pSI neurons on the flight and freezing response, a plastic shield cover (nest) was introduced into the corner of the enclosure, mice were tested in 15-min sessions, consisting of 5-min light OFF, 5-min light ON (15 s off, 15 s laser on, and 15 s off), and 5-min light OFF periods. Latency to the nest was counted manually. The time spent in the center area and the velocity of mice were automatically analyzed by ANY-maze.

To investigate the effect of optical inhibition of pSI^Thy1^ neurons on general locomotion, mice were tethered to an optic fiber, habituated in the home cage for 15 min, and then individually placed in the center of a novel plastic arena (48 × 48 × 48 cm) at the start of the session and allowed to freely explore for 9 min in three 3-min epochs (Pre-Laser-Post). During the laser epoch, mice received continuous bilateral optogenetic inhibition with the light intensity same as that optimized for inhibiting attacks on previous testing (3-4 mW). The video was captured with an overhead mounted camera. The movement was tracked and measured using automated video-tracking software (ANY-maze).

#### Feeding Behavior Measurements

To assess feeding behavior in response to optogenetic inhibition of pSI^Thy1^ neurons, mice were food-restricted for 24 h before the experiment. The next day, the mice were habituated in the behavioral context for 15 min after tethering to the bilateral optic fibers. Each inhibition test ran for ∼10 min and was videotaped and scored for the following parameters: latency to feeding offset, probability of feeding trials interrupted within 3 s, duration of feeding, and feeding events. While they were eating, the mice received 15-s photoinhibition with light intensity the same as that optimized for inhibiting attacks in previous tests (3-4 mW, ∼20 times, 15 s ON/OFF). The control test used the same procedure, but the mice received sham blue laser inhibition (0 mW, ∼20 times, and 15 s ON/OFF).

#### HM4Di-mediated Neural Silencing and HM3Dq-mediated Neural Activation

To label and chronically silence pSI^-PAG^ neurons (Armbruster et al., 2007), retrograde AAV-Retro-Cre-EGFP (60 nl/side) was used to bilaterally infect PAG neurons, and AAV-hSyn-DIO-hM4Di-mCherry (a Cre-dependent pharmacogenetic silencing virus, 150 nl/side) was used to bilaterally infect Cre+ pSI neurons in singly-housed male C57 mice. The control singly-housed males were bilaterally injected with AAV-Retro-Cre-EGFP (60 nl/side) in the PAG and AAV-hSyn-DIO-mCherry (150 nl/side) in the pSI. To silence the pSI^Thy1^ neurons, AAV-hSyn-DIO-hM4Di-mCherry (a Cre-dependent pharmacogenetic silencing virus, 150 nl/side) was directly used to bilaterally infect the pSI^Thy1^ neurons in singly-housed male Thy1-Cre mice. Control male Thy1-Cre mice were bilaterally injected with AAV-hSyn-DIO-mCherry (150 nl/side) in the pSI. Three weeks later, a randomly-selected socially-housed adult C57 male intruder (20-30 g) was introduced and left for 15 min to evaluate the aggression level of the tested mouse. Mice in the test group were then intraperitoneally injected with 50 μl saline or CNO (1 mg/kg, Sigma, C0832, stored at –20°C and dissolved in 0.9% sterile saline to a volume of 0.5 mg/ml before use). Thirty minutes after injection, the same behavioral paradigm was repeated for 15 min to evaluate the aggression level of the tested mice again. The neural activation by natural aggression was assessed by c-Fos staining on the next day.

To label and chronically activate the pSI^-PAG^ neurons, retrograde AAV-Retro-Cre-EGFP virus (60 nl/side) was used to unilaterally infect PAG neurons, and AAV-hSyn-DIO-hM3Dq-mCherry (a Cre-dependent pharmacogenetic activating virus, 100 nl/side) was used to unilaterally infect the Cre+ pSI neurons in socially-housed male C57 mice. The controls were unilaterally injected with AAV-Retro-Cre-EGFP (60 nl/side) in the PAG and AAV-hSyn-DIO-mCherry (100 nl/side) in the pSI. To activate the pSI^Thy1^ neurons, AAV-hSyn-DIO-hM3Dq-mCherry (a Cre-dependent pharmacogenetic activating virus, 100 nl/side) was directly used to unilaterally infect the pSI^Thy1^ neurons in socially-housed male Thy1-Cre mice. The control male Thy1-Cre mice were unilaterally injected with AAV-hSyn-DIO-mCherry (100 nl/side) in the pSI. Three weeks later, the mice were intraperitoneally injected with 25 μl saline or CNO (0.5 mg/kg), and a randomly-selected singly-housed adult C57 male intruder (20-30 g) was introduced into the home cage of the tested mouse and left for 15 min to evaluate the aggression level of the pharmacogenetically-activated and control mice. Thirty minutes after injection, the same behavioral paradigm was run for 15 min to evaluate the aggression level of the mice again. The neural activation by CNO (0.5 mg/kg) was assessed by c-Fos staining on the next day.

#### ChR2-mediated Neural Activation

To label the pSI^-PAG^ neurons and activate them in a time-locked manner (Boyden et al., 2005), retrograde AAV-Retro-Cre-EGFP (60 nl/side) was used to unilaterally infect PAG neurons, and AAV-Ef1α-DIO-hChR2 (H134R)-EYFP (a Cre-dependent optical activating virus, 150 nl/side) was used to unilaterally infect the Cre+ pSI neurons in male and female C57 mice. The controls were unilaterally injected with AAV-Retro-Cre-EGFP (60 nl/side) in the PAG and AAV-Ef1α-DIO-EYFP (100 nl/side) in the pSI. To optogenetically activate pSI^CaMKIIα^ neurons, C57 males and females were unilaterally injected with AAV-CaMKIIα-hChR2 (H134R)-mCherry (60 nl/side) in the pSI. The control animals were unilaterally injected with AAV-CaMKIIα-mCherry (60 nl/side) in the pSI. After injection, mice were allowed 3-4 weeks to recover. For *in vivo* optogenetic manipulation, an optical fiber cannula (200 μm) was implanted into the right pSI. To acutely manipulate the pSI-PAG, pSI-BNST or pSI-VMH circuit, an optical fiber cannula (200 μm) was implanted into the right VL/LPAG, BNST, or VMH. The optical fiber was connected to a laser source using an optical fiber sleeve. The light intensity was the same as that optimized for eliciting attacks on previous testing days (5 ms, 5-40 Hz, 0-3.50 mW, 20 times, 15 s ON and 15 s OFF). Randomly-selected socially- or singly-housed adult males or females were introduced into the home cage of the tested mouse, one at a time and each for 10-20 min. Mice with missed injection or cannula placements were excluded. Control mice expressing only EYFP/mCherry underwent the same procedure and received the same intensity of laser stimulation. For the intensity modulation assay, the pSI^-PAG^ neurons were photostimulated with 14 different light intensities (0, 0.03, 0.10, 0.17, 0.26, 0.34, 0.40, 0.68, 1.17, 1.58, 1.98, 2.24, 2.70, and 2.98 mW) in diverse aggressive behaviors in male and female mice (Figure 6). Different laser intensities were delivered in a pseudo-random order and each intensity of laser (5 ms, 40 Hz, 15 s ON and 30 s OFF) for five types of aggression were tested for 5-8 trials. 1- to 3-day-old naive C57 pups, crickets, adult C57 or BALB/c mice of both sexes were introduced as the opponents, the tested male and female mice were socially-housed for at least 3 weeks before behavioral tests (see the above description of aggression tests). A plot of log (photostimulation intensity) vs aggressive behaviors was fitted with the nonlinear sigmoidal curve (yielded a nonlinear relationship).

To optogenetically activate the pSI^Thy1^ neurons projecting to the PAG, Thy1-Cre male mice were unilaterally injected with rAAV-CAG-retro-FLEX-Flpo mixed with CTB555 (Dilution ratio = 4:1, 100 nl /side) in the PAG and AAV-fDIO-hChR2 (H134R)-EYFP (100 nl/side) in the SI. Three weeks later, we used the same aggression behavioral protocol (see above) to test the role of pSI^Thy1^ neurons projecting to the PAG in inter-male aggression and predatory aggression.

#### ChR2-mediated Neural Activation before and after Acute Social Defeat

To optogenetically activate pSI^-PAG^ neurons before and after ASD by CD1 mice, the right PAG of C57 males was unilaterally injected with retrograde AAV-Retro-Cre-EGFP (80 nl/side), and AAV-Ef1α-DIO-hChR2 (H134R)-EYFP (100 nl/side) was used to unilaterally infect the right Cre+ pSI neurons. For *in vivo* optogenetic manipulation, an optical fiber cannula (200 μm) was implanted into the right pSI. Before testing, the virus-injected mice were socially-housed. On the test day, the tested mice habituated in the behavioral context for 15 min after tethering optic fibers bilaterally. Randomly-selected socially-housed adult male intruders were introduced into the home cage of the tested mouse, one time for each intruder for 10-20 min. The pSI^-PAG^ neurons were photostimulated with 7 different light intensities (0.00, 0.33, 0.54, 0.75, 1.51, 2.72, and 4.28 mW) in the first behavior test (before ASD) for ∼30 min (see the above description of aggression tests). Different laser intensities were delivered in a pseudo-random order and each intensity of laser (5 ms, 40 Hz, 15 s ON and 30 s OFF) was tested for 5-8 trials. To explore the acute impacts of social defeat on the level of the attack triggered by photostimulation of pSI neurons, the tested mouse was then exposed to a CD-1 aggressor for 10 min. During the exposure, the tested mouse was physically defeated by the aggressor for ∼15 times. After 10-min defeat, the tested mouse was returned to its home cage, followed by the next behavior test for ∼30 min (after ASD).

#### Optogenetic Inhibition of Neural Activity

To label and silence in a time-locked manner the pSI^Thy1^ neurons or the pSI^Thy1^ neural terminals in the VL/LPAG, AAV-EF1a-DIO-GtACR1-P2A-GFP-WPRE (a Cre-dependent optogenetic silencing virus, 150 nl/side) was used to bilaterally infect the pSI^Thy1^ neurons in singly-housed male Thy1-Cre mice. For *in vivo* optogenetic manipulation in awake behaving animals, optical fiber cannulas (diameter, 200 μm) were bilaterally implanted into the pSI or the VL/LPAG. Three weeks after viral infection, a randomly-selected socially-housed adult male intruder was introduced into the home cage of the tested mouse and left for ∼15 min to evaluate the aggression level of the singly-housed adult male Thy1-Cre mouse. To assess the effects of optogenetic inhibition of pSI^Thy1^ neurons, five types of robust natural aggressive behaviors were used (offensive inter-male aggression, when a singly-housed Thy1-Cre male in its home cage attacked a socially-housed C57 male mouse; offensive male-female aggression, when a singly-housed Thy1-Cre male in its home cage attacked a socially-housed C57 female mouse; offensive female-male aggression, when a singly-housed Thy1-Cre female in its home cage attacked a socially-housed C57 male mouse; pup-directed aggression, when a singly-housed Thy1-Cre virgin male in its home cage attacked a pup; and predatory aggression, when a singly-housed Thy1-Cre virgin male in its home cage attacked crickets, see Figure 7. Each inhibition test ran for ∼15 min and was videotaped and scored for the following parameters: latency to attack offset, probability of attack trials interrupted within 3 s, duration of the attack, and attack distribution. The light intensity was the same as that optimized for inhibiting attacks on previous testing days (3-4 mW, ∼20 times, and 15 s ON/OFF). Mice with the missed injection or cannula placements were excluded. Control mice underwent the same procedure and received the same intensity of laser stimulation.

#### Fiber Photometry

Following AAV-EF1a-DIO-GCamp6m or AAV-EF1a-DIO-EYFP injection in Thy1-Cre males, an optical fiber (230 μm OD, 0.37 NA) was placed in a ceramic ferrule (2.5 mm OD, 126 μm ID) and inserted toward the pSI. The ferrule was affixed with a skull-penetrating M1 screw and dental acrylic. To enable recovery and AAV expression, mice were housed individually for at least 10 days after virus injection. A fiber photometry system (Thinker Tech Nanjing Biotech Ltd, Nanjing, China) was used for recording (Gunaydin et al., 2014, Li et al., 2018). To record fluorescence signals, the beam from a 488-nm laser (OBIS 488LS; Coherent, Santa Clara, CA, USA) was reflected with a dichroic mirror and focused with a 10× objective lens (NA = 0.3; Olympus, Tokyo, Japan). An optical fiber (230 μm OD, NA = 0.37; 2 m long) guided the light between the commutator and the implanted optical fiber. To minimize GCaMP bleaching, the laser power at the tip of the optical fiber was adjusted to a low level (0.03–0.04 mW). The GCaMP6 fluorescence was filtered with an EYFP bandpass filter and collected by a photomultiplier tube (R3896; Hamamatsu Photonics). An amplifier converted the photomultiplier current output to a voltage signal, which was further filtered through a low-pass filter (40 Hz cut-off; Brownlee 440). The analog voltage signals were digitized at 500 Hz (Power 1401 digitizer, Cambridge Electronic Design) and sampled with software (TDMS). Fiber-photometric recording data were exported as MATLAB files for further analysis, and fiber photometry-related aggression behavioral data were analyzed using MATLAB. All the raw fluorescence data (*F*) were smoothed with a moving average filter (10-ms span) and then segmented and aligned to the onset of behavioral events within individual trials or bouts. The fluorescence change values (Δ*F/F*) were calculated as (F–F0)/(F0–V_offset_), where F0 is the baseline fluorescence signal averaged over a 2-s/4-s time-window prior to a trigger event and V_offset_ is the fluorescence signal recorded before the cannula was connected to the optical fiber above the pSI. Δ*F/F* values are presented as heatmaps or average plots with a shaded area indicating the SEM.

On testing days, a water port and food were left inside the tested mouse’s cage for ∼10 min. During that period, the mouse was free to approach, lick, and/or eat. A non-aversive stimulus consisting of an object (a cotton ball without liquid), or a cotton ball with 10 μl DMSO (dimethyl sulfoxide), male urine, or TMT (2, 5-dihydro-2, 4, 5-trimethylthiazoline) was then left for ∼10 min in the home cage of a subject Thy1-Cre mouse. The aversive stimuli were air-puffs (delivered by a hand-pumped compressor with its opening ∼3 cm from the mouse’s nares; one press of the pump generated a gentle ∼0.5-s puff), tail suspension (the tail was gripped and the mouse lifted off the floor of its cage for ∼4 s, with at least 30 s between each lift), and a flying object (a flattened glove was introduced into the tested mouse’s cage above its head, and then approached its body). The onset of each stimulus triggered data alignment. Social and non-social entry refers to the threat period after an intruder mouse (female C57, male C57, male CD-1, or male C57 entry preceding attack) or object (a 10-ml plastic centrifuge tube) was first introduced into the home cage of a subject Thy1-Cre mouse. The time of each entry triggered data alignment. Aggressive social behaviors of these subject Thy1-Cre mice were recorded during the application of various contextual cues.

We calculated the average Δ*F/F* during the following stimuli and aggressive behaviors: offensive inter-male attack (all attack), first attack (an attack not followed by another attack within 4 s), continuous attack (an attack following or followed by another attack within 4 s), last attack (a continuous attack not followed by another attack within 4 s), sniff without attack (a sniff not followed by an attack within 4 s), sniff preceding attack, rattle without attack (a rattle not followed by an attack within 4 s), rattle preceding attack, threat (subject male mouse threatened other males with boxing postures but no bite), male-female attack, defensive inter-male attack, predatory approach and attack, pup-directed approach and attack, female-male sniff and attack, and multiple social and non-social cues based on the peristimulus time histograms (PSTHs). The non-parametric Wilcoxon signed-rank test or t-test was used to determine whether the AUC per second before, during and after events were significantly different from control (2/4 s before a trigger event).

#### Decoder Analysis by the Support Vector Machine (SVM) Algorithm

Population decoding analysis was used to evaluate whether diverse aggressive behaviors could be extracted based on the pSI^Thy1^ neuronal Ca^2+^ profile of individual trials. Analysis with the support vector machine (SVM) algorithm was performed with the LIBSVM library (https://www.csie.ntu.edu.tw/~cjlin/libsvm) according to the library’s documented instructions.

To study the correlation of behavioral performance and population neuronal decoding, stimulus identity was decoded from the activity of all pSI neurons using the cross-validated SVM decoder. The total number of trials of types of social behaviors ranged from 26 to 680. To ensure that the result came from sampling all trials rather than a limited number of them, we alternated the number of trials (from 10 to 150) in the calculations and used the classification accuracy test with the SVM with the linear kernel or the radial basis function (RBF) kernel. The classification accuracy from the linear kernel was better than the RBF kernel with default parameters, and the sampling results suggested the classification accuracy of a pre-selected 50 trials was relatively higher and suitable to the test in the social behaviors with limited collected trials. Thus, 50 trials of the Ca^2+^ signals for each type of social behavior were randomly selected for the classification accuracy test with the linear kernel, and the Ca^2+^ signals for each behavior trial were binned into 10-ms windows and grouped by social behavior type. The Ca^2+^ signals were normalized to the [0, 1] range. In each calculation, 90% of the trials were randomly partitioned to a training set, and the remaining 10% was used as the test set, with no overlap between the training and testing sets. All trials were then randomly re-partitioned, and the training and testing repeated 10 times. The SVM was trained with the training set, and the trained SVM was used to classify the test set. Each of the results was labeled as correct if the classified label matched the behavioral type, or incorrect otherwise. Because the training and testing set never overlapped and all trials were likely to be used in both the training and testing sets due to the large repetition number, the average decoding accuracy was equal to the cross-validation accuracy in a resampled form. For the recording group and shuffled control group, the procedure was repeated 1000 times with the behavioral label for each trial randomly re-partitioned and redistributed.

Bar plots of decoder accuracy were generated using held-out test data for a cross-validated SVM decoder trained on the time-averaged responses of population neuronal activity in a window from 4 s before to 8 s after stimulus presentation. Time-evolving plots of decoder accuracy were constructed by training a separate cross-validated decoder on the time-averaged activity of population neuronal activity in 10-ms sliding steps from 4 s before to 8 s after stimulus presentation. Decoder performance is reported as the average prediction accuracy on the held-out test data; chance accuracy is 1/2 for the two-class decoder and 1/3 for the three-class decoder.

#### Single-unit Recording and Data Analysis

Modified electrode arrays were implanted dorsal to the pSI (Lin et al., 2007). The electrodes consisted of 32 single microwires (California Fine Wire Co., Grover Beach, CA, USA). One ground wire was soldered to a 32-channel connector (Omnetics Connector Corp., Minneapolis, MN, USA). Mice were allowed to recover for at least 7 days, and then the electrodes were connected to a 32-channel preamplifier head-stage (Plexon Inc., Dallas, TX, USA). Twenty-four hours after recording the activity of a pSI neuron, a behavioral test was given to examine the firing pattern of the same neuron. During the recording sessions, all signals recorded from each electrode were amplified, filtered between 0.1 Hz and 10 kHz, and spike waveforms were digitized at 40 kHz. Spikes were sorted using the software Offline Sorter (Plexon). Units were accepted only if a distinct cluster was visible in a two-dimensional plot of the largest two principal components. In total, 9 mice were implanted with electrodes and used for data collection. Neurons with mean firing rates > 0.5 Hz were included in the analysis. Then analysis was performed using MATLAB, referencing neural activity to behavior. A given neuron that significantly responded to a defined behavior would be reflected by its reliable responses across different trials. The responses of each neuron were averages of 2-20 behavior trials. We then calculated the averaged response during attack, investigation of object, and male, based on the z-scored PSTHs. For each recorded unit, PSTHs were z-score transformed by subtracting the mean firing rate and dividing by the standard error of each unit’s firing rate. To test the significance of changes in firing rate, we used individual unit analysis. A non-parametric Wilcoxon signed-rank test was performed on each unit to determine whether the mean firing rate during the event (e.g., attack, sniffing object, and sniffing male) was significantly different from baseline (from –2 s to –1 s), and units were classified into three populations: excited, inhibited, or not changed. The responses and relative proportions of attack-excited, attack-inhibited, and attack-no-response units before and during the events were compared using the t-test.

#### *In Vitro* Electrophysiology

Mice were deeply anesthetized with pentobarbital sodium (100 mg/kg, i.p.) and decapitated. The whole brain was quickly dissected into ice-cold oxygenated artificial cerebrospinal fluid (aCSF), then cut coronally into 300-μm slices on a microtome (VTA-1200S; Leica). Slices containing the pSI were transferred to an incubation chamber filled with aCSF and incubated for at least 1 h at room temperature (RT) (24-26°C). At RT, the slices were transferred to a recording chamber on the stage of a fluorescence microscope (BX51WI; Olympus). Patch electrodes were pulled from borosilicate capillaries (BF150-86-10; Sutter Instrument Co, Novato, CA, USA). Recordings were made with a MultiClamp 700B amplifier (Molecular Devices). Signals were low-pass filtered at 10 kHz and digitized at 10 kHz (MICRO3 1401, Cambridge Electronic Design). Data were acquired and analyzed using Spike2 7.04 software (Cambridge Electronic Design). For photostimulation of pSI neurons, an optical fiber (diameter 200 µm) was coupled to a 473-nm LED. The fiber was glued to a stainless-steel tube (ID 250 µm, OD 480 µm). The tip of the fiber was trimmed and polished, submerged in aCSF, and positioned above the stimulation site. The blue light was controlled by a pulse stimulator (PG-4000A; Cygnus Technologies, Southport, NC, USA). Whole-cell patch-clamp recordings under both voltage and current clamp were performed in neurons that expressed EYFP/mCherry fluorescence. To test for the effects of photostimulation, LED-generated blue light pulses at different frequencies (5, 10, 20, and 40 Hz) were applied to recorded neurons. For recordings from virus-infected neurons, pipettes were filled with a K^+^-based low Cl^−^ internal solution containing (in mM) 145 KGlu, 10 HEPES, 0.2 EGTA, 1 MgCl2, 4 Mg-ATP, 0.3 Na2-GTP, 10 Na2-Phosphocreatine (pH 7.3 adjusted with KOH; 295 mOsm). Membrane potentials were corrected for ∼10 mV liquid junction potential. To test for the effects of pharmacogenetic activation or inhibition of pSI^Thy1^ neurons, 5 or 10 µM CNO was added to the aCSF perfusion. To test the effects of optogenetic inhibition of pSI^Thy1^ neurons, LED-generated blue light pulses at constant frequencies were applied to the recorded GtACR1-expressing pSI^Thy1^ neurons. To test the excitatory effects of photoactivation of pSI^Thy1^ projection terminals on VL/LPAG neurons (Figure S7), LED-generated blue light pulses at a constant frequency were applied to the recorded PAG neurons. The light-evoked currents in the VL/LPAG neurons by photostimulating axonal projections from the pSI to these neurons were recorded in the absence or presence of the bath-applied glutamate receptor antagonist CNQX (20 μM), and the GABA receptor antagonist Picrotoxin (100 μM).

#### Histology

After experiments, the mice were anesthetized and perfused with saline followed by 4% paraformaldehyde in 0.1 M phosphate buffer (PBS). The brain was then removed and placed in 4% paraformaldehyde buffer at 4°C for overnight fixation, after which the brain was cryoprotected in 30% sucrose (wt/vol) at 4°C. Coronal sections were cut at 40 μm on a cryostat (Leica CM1900) for imaging. After rinsing with 0.3% Triton-X 100 (vol/vol) in 0.1 M PBS (30 min) or ice-cold methanol (10 min) and blocking with 10% (wt/vol) normal bovine serum for 1 h at room temperature, sections were incubated with the following primary antibodies (12-24 h at 4°C): anti-GAD67 (1:500, mouse, Millipore), anti-GAD65/67 (1:400, rabbit, Abcam), anti-ChAT (1:200, rabbit, Millipore), anti-CaMKII (1:200, rabbit, Abcam), and anti-c-Fos (1:2500, rabbit, Calbiochem). After rinsing, sections were incubated with fluorophore-conjugated secondary antibody (1:1000; Millipore) for 2 h at RT. Antibodies were diluted in PBS containing 4% bovine serum albumin and 0.2% Triton X-100. To induce natural aggression in males for c-Fos quantification, a socially-housed adult male intruder (C57) was introduced into the home cage of a highly sexually-experienced adult male C57 resident mouse or male Thy1-Cre resident mouse (AAV-DIO-EYFP virus was injected into the pSI in the Thy1-Cre mice two weeks before) for 10 min of aggression. Resident mice showing the only investigation but not attack toward intruders in the 10-min test period were used as a control group. To assess the activation of pSI neurons by optogenetics or pharmacogenetics, adult male mice were singly housed in their home cage on the first day, and the mice were stimulated with light for 10 min (5 ms, 40 Hz, 1-2 mW, 20 times, 15 s ON and 15 s OFF) or were intraperitoneally injected with CNO (0.5 mg/kg) on the next day. These mice were perfused 1.5 h later, and sections were cut as described above. After the anti-c-Fos immunohistochemistry reaction, nuclei were stained with DAPI, and confocal images were captured using 10×, 20×, and 60× objectives (Olympus FV-1200). Cell counting was carried out manually using Fiji (NIH).

### QUANTIFICATION AND STASTISTICAL ANALYSIS

No statistical methods were used to pre-determine sample sizes. All data were randomly collected. We first determined whether the data values came from a normal distribution with the D’Agostino–Pearson omnibus, Shapiro–Wilk, and Kolmogorov–Smirnov normality tests. In experiments with paired samples, we used a two-tailed paired t-test and the Wilcoxon matched-pairs signed-rank test for parametric and non-parametric data, and the Friedman test for nonparametric data. In all other experiments, we used a t-test, the t-test with Welch’s correction for unequal standard deviation, or ANOVA for parametric data, and a Mann-Whitney or Kruskal-Wallis test for non-parametric data. All values were calculated with Prism 6.0 or MATLAB.

### KEY RESOURCES TABLE

## Supplemental Figure Legends

**Figure S1.**
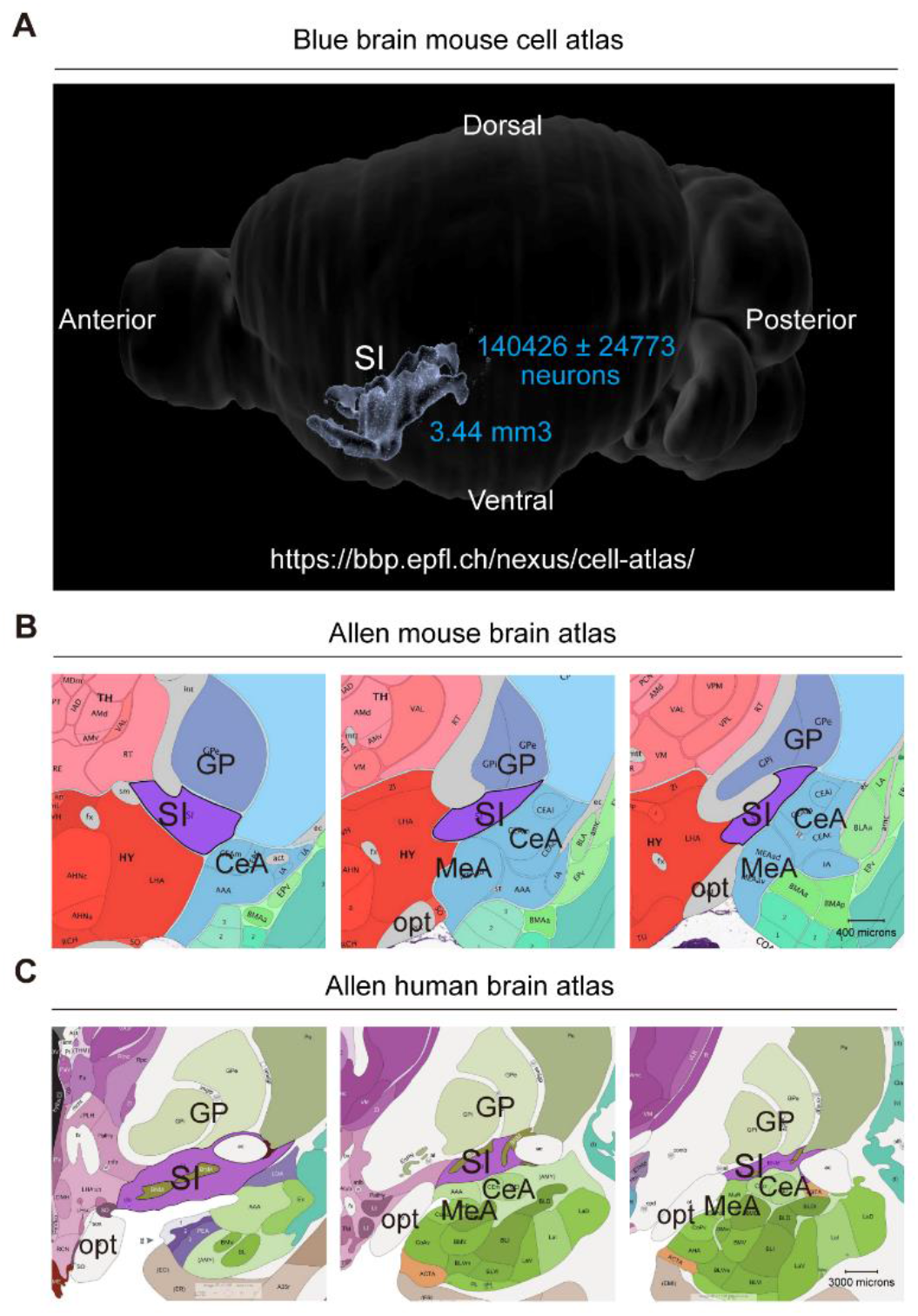
Substantia Innominata in the Mouse and Human Brain Reference Atlases, Related to Figure 1 (A) Structure of the substantia innominata (SI) as shown in the EPFL Blue Brain Reference Atlas (2019) (https://bbp.epfl.ch/nexus/cell-atlas/). (B) The SI as shown in the Allen Mouse Brain Reference Atlas (2011, http://mouse.brain-map.org/static/atlas<u>). The anatomical reference atlas illustrates the SI (purple) in the adult mouse brain in coronal planes (from left to right, planes 60, 63, and 66).</u> (C) The SI as shown in the Allen Human Brain Reference Atlas (using modified Brodmann annotation, http://atlas.brain-map.org/atlas?atlas=265297126). The anatomical reference atlas illustrates the SI (purple) in an adult human brain (32 years old) in coronal planes (from left to right, planes 27, 32, and 37)

**Figure S2.**
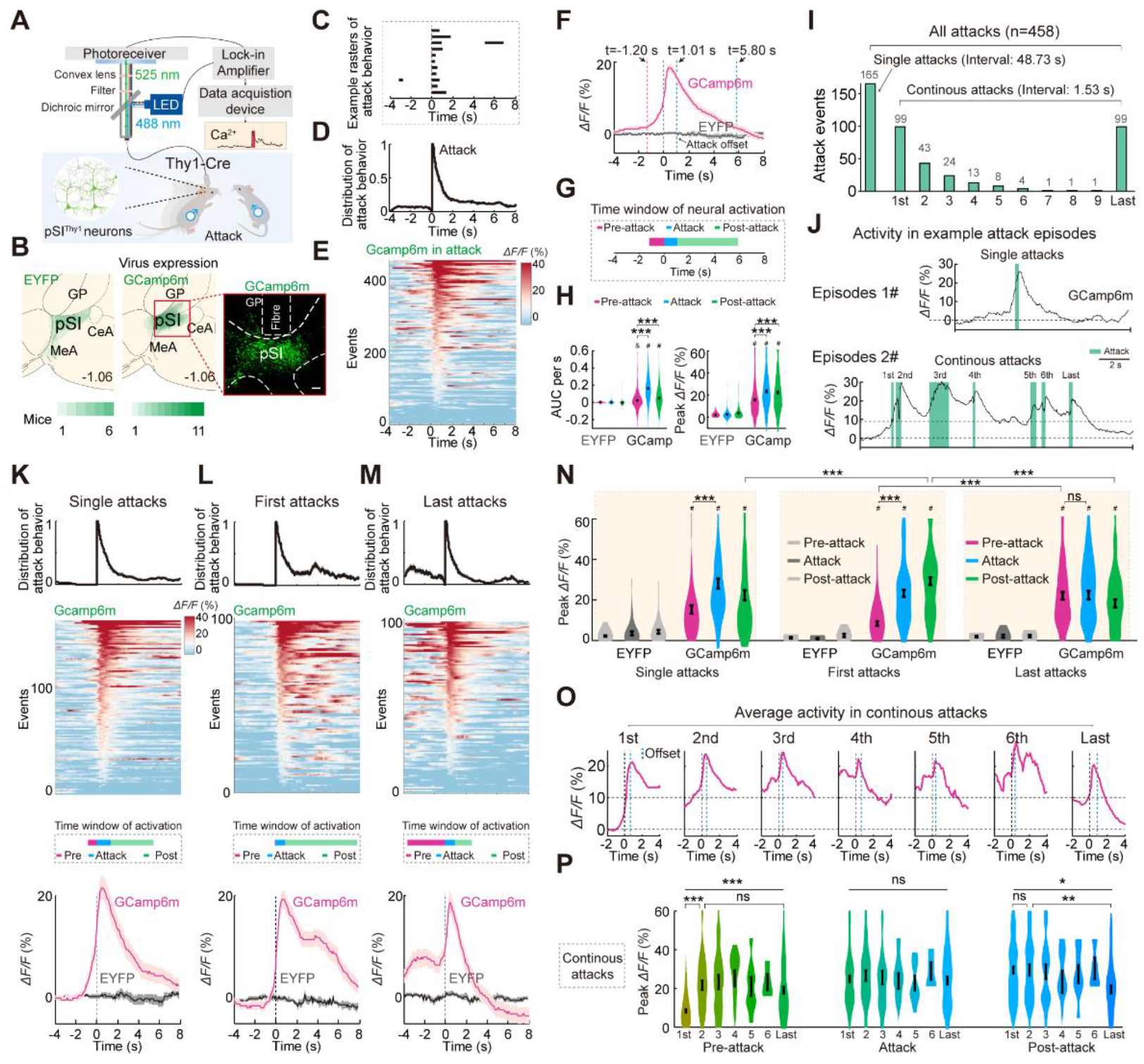
Correlation of pSI^Thy1^ Neuronal Ca^2+^ Dynamics with Attack Patterns, Related to Figure 2 (A) Setup for fiber photometric recording in pSI^Thy1^ neurons during inter-male aggression. (B) Overlay of EGFP (left) and GCaMP6m (middle) expression in the pSI. Right panel, a representative image (scale bar, 100 μm) of GCaMP6m (green) and the optical fiber track. (C) An example raster plot of single attack distribution from 11 mice. (D) Summarized distribution of attack episodes plotted by the percentage of trials showing attack at different time points (n = 458 attacks). (E) Heatmap of GCaMP6m Δ*F/F* signals from pSI neurons in aggression (n = 458 attack trials). (F) Representative Δ*F/F* of GCaMP6m (magenta) and EYFP (gray) signals before, during, and after an attack. Note that the elevation of Ca^2+^ signaling (t = –1.2 s) preceded the attack behavior (onset at t = 0 s). (G) Time windows of pSI^Thy1^ neuronal activation before, during, and after attacks. (H) Area under the curve (AUC) per second (left) and peak Δ*F/F* (right) of GCaMP6m and EYFP signals before, during, and after an attack. Mann-Whitney test and Wilcoxon matched-pairs signed-rank test; \*\*\**P* < 0.001 between the groups indicated. ‘#’ and ‘&’ above each bar indicate that the AUC per second (left) or peak Δ*F/F* (right) during each behavior period was (*P* < 0.001) and wasn’t (*P* > 0.05) significantly changed compared with the EYFP group, determined by Mann-Whitney test. (I) Distribution of the incidence of single attacks and continuous attacks with different episode orders. (J) Representative GCaMP6m signals during a single attack (upper) and continuous attacks with different episode orders (lower). Green bars, episodes of attack. (K-M) From top to bottom: distribution of attack behavior episodes, heatmaps of GCaMP6m Δ*F/F* signals from pSI^Thy1^ neurons, time windows of pSI^Thy1^ neuronal activation, and Δ*F/F* of the EYFP and GCaMP6m signals from pSI^Thy1^ neurons before, during, and after single attacks (K, n = 165 trials) and the first (L, n = 99 trials) and last (M, n = 99 trials) episodes of continuous attacks. (N) Peak Δ*F/F* of EYFP and GCaMP6m signals before, during, and after single attacks (left) and the first (middle) and last (right) episodes of continuous attacks. The EYFP and GCaMP6m signals in each type of attack behavior were compared using the Mann-Whitney test, Wilcoxon matched-pairs signed-rank test, and two-tailed paired t-test; from left to right, \*\*\**P* < 0.001, \*\*\**P* < 0.001, \*\**P* = 0.0049, \*\*\**P* < 0.001, \*\*\**P* < 0.001, *P* = 0.5349. The GCaMP6m signals in the three types of attack behavior were compared using the Mann-Whitney test; \*\*\**P* < 0.001. ‘#’ above each bar indicates that peak ΔF/F (right) during each behavior period was significantly (\*\**P* < 0.01) changed compared with the EYFP group, determined by Mann-Whitney test. (O) Averaged Δ*F/F* of GCaMP6m dynamic signaling of continuous attacks with different episode orders. (P) Peak Δ*F/F* of GCaMP6m signals before, during, and after continuous attacks with different episode orders. Mann-Whitney test, unpaired t-test, and Kruskal-Wallis test; from left to right, \*\*\**P* < 0.001, \*\*\**P* <0.001, *P* = 0.2623, *P* = 0.9688, *P* = 0.9867, \**P* = 0.0189, \*\**P* = 0.005. ns, not significant. Data are presented as the mean ± SEM.

**Figure S3.**
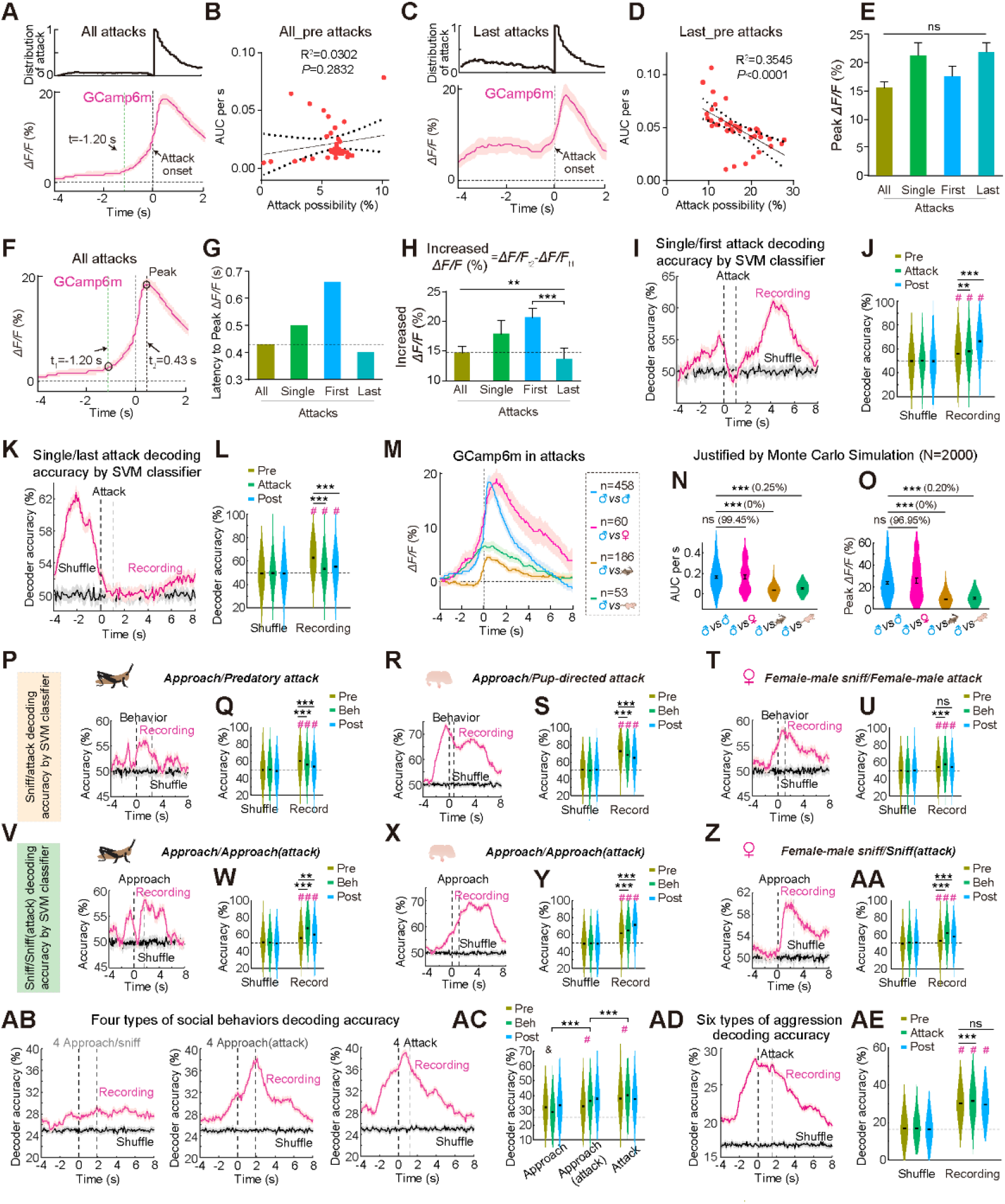
Correlation of Different Aggressive Behaviors with the Differential Calcium Activity Levels in pSI^Thy1^ Neurons, Related to Figures 2-3, and S2 (A) Distribution of attack episodes (n = 458 attacks, upper) and representative Δ*F/F* of GCaMP6m signals before and during attacks (n = 458 attacks, lower). Note that elevation of the Ca^2+^ signal (t = –1.2 s) preceded the attack behavior (onset at t = 0 s). (B) Linear regression of the AUC per second of GCaMP6m signals prior to the attack behavior and probability of all_pre attacks (attacks during the 4s window prior to all episodes of attack behavior, n = 458). (C) Distribution of last attack episodes (n = 99 attacks, upper) and representative Δ*F/F* of GCaMP6m signals before, during, and after the last episode of continuous attacks (n = 99 attacks, lower). (D) Linear regression of the AUC per second of GCaMP6m signals and probability of last_pre attacks (attacks during the 4s window prior to the last episode of continuous attacks, n = 99). (E) Peak Δ*F/F* of GCaMP6m in all attack behavior, single attack behavior, and the first and last episodes of attack behavior. *P* = 0.1270, Kruskal-Wallis test. (F) Sample decoding accuracy of the pSI neuronal Ca^2+^ signals in single and the first episode of continuous attack trials. (G) Averaged decoding accuracy of pSI neuronal activity before, during, and after attack trials under recording and shuffled conditions. ‘#’ above each bar indicates that the decoding accuracy during each behavior period was significantly increased (*P* <0.05) compared with that in the shuffled period, assessed by the Mann-Whitney test. Decoding accuracy in recording differences before, during, and after attack trials was assessed by the Mann-Whitney test. From left to right, \*\*\**P* <0.001, \*\*\**P* <0.001. (H) Sample decoding accuracy of pSI neuronal Ca^2+^ signals in single and the last episode of continuous attack trials. (I) Averaged decoding accuracy of pSI neuronal activity before, during, and after attack trials under recording and shuffled conditions. Decoding accuracy in recording differences before, during and after attack trials was assessed by the Mann-Whitney test. From left to right, \*\*\**P* <0.001, \*\*\**P* <0.001. (J) Averaged ΔF/F of Ca^2+^ signals in pSI neurons before, during, and after male-male, female-male, predatory, and pup-directed attacks. (K) Peak ΔF/F of Ca^2+^ signals during diverse attacks. Comparison of diverse attacks was determined by Monte Carlo simulation with a subset of the male-male data 2000 times. *P*-value of comparison of subset of Ca^2+^ data associated with the diverse attacks was determined by the Wilcoxon rank-sum test. ‘ns (96.95%)’ indicates that these two samples were not significantly different (*P* >0.05). Monte Carlo simulation showed that 96.95% of the 2000 comparisons were from populations with identical distributions. ‘*** (0.20%)’ indicates that these two samples was significantly different (*P* <0.001), and Monte Carlo simulation showed that 0% of 2000 comparisons were from populations with identical distributions. (L-Q) Sample decoding accuracy of the pSI neuronal Ca^2+^ signals and averaged sample decoding accuracy of pSI neuronal activity under recording and shuffled conditions compared between the cricket-directed approach and predatory attack trials (L, M), the pup-directed approach and pup-directed attack trials (N, O), and the female-male sniff and female-male attack trials (P, Q). ‘#’ above each bar indicates that decoding accuracy during each behavior period was significantly higher (*P* <0.05) than that in the shuffles period, determined by the Mann-Whitney test. Decoding accuracy in recording differences before, during, and after attack trials was determined by the Mann-Whitney test. From left to right, \*\*\**P* <0.001, \*\*\**P* <0.001. (R-W) Sample decoding accuracy of the pSI neuronal Ca^2+^ signals and averaged sample decoding accuracy of pSI neuronal activity under recording and shuffled conditions compared between the cricket-directed approach and approach (attack) trials (R, S), the pup-directed approach and approach (attack) trials (T, U), and the female-male sniff and sniff (attack) trials (V, W). From left to right, \*\*\**P* <0.001, \*\*\**P* <0.001. (X) Sample decoding accuracy of the pSI neuronal Ca^2+^ signals and averaged sample decoding accuracy of pSI neuronal activity under recording and shuffled conditions in cricket-directed, pup-directed, female-male, and inter-male approaches or sniffs (left), approaches or sniffs preceding an attack (middle), and attack trials (right). (Y) Averaged sample decoding accuracy in four types of approaches/sniffs, approaches/sniffs preceding attacks, and attacks under recording conditions as shown in (AB). ‘#’ or ‘&’ above each bar indicates that the decoding accuracy during each behavior period was (*P* <0.05) or was not (*P* >0.05) significantly higher than that before each behavior period, determined by the Mann-Whitney test. Decoding accuracy in recording differences in the three behaviors was determined by the Mann-Whitney test. From left to right, \*\*\**P* <0.001, \*\*\**P* <0.001. (Z) Sample decoding accuracy of the pSI neuronal Ca^2+^ signals under recording and shuffled conditions in predatory, pup-directed, female-male, inter-male, male-female, and inter-male defensive attack trials). (AA) Averaged sample decoding accuracy under recording and shuffled conditions. ‘#’ above each bar indicates that the decoding accuracy during each behavior period was significantly (*P* <0.05) higher than that before each behavior period, determined by the Mann-Whitney test. Decoding accuracy in recording differences was determined by the Mann-Whitney test. From left to right, \*\*\**P* <0.001, \*\*\**P* <0.001. ns, not significant. Data are presented as the mean ± SEM.

**Figure S4.**
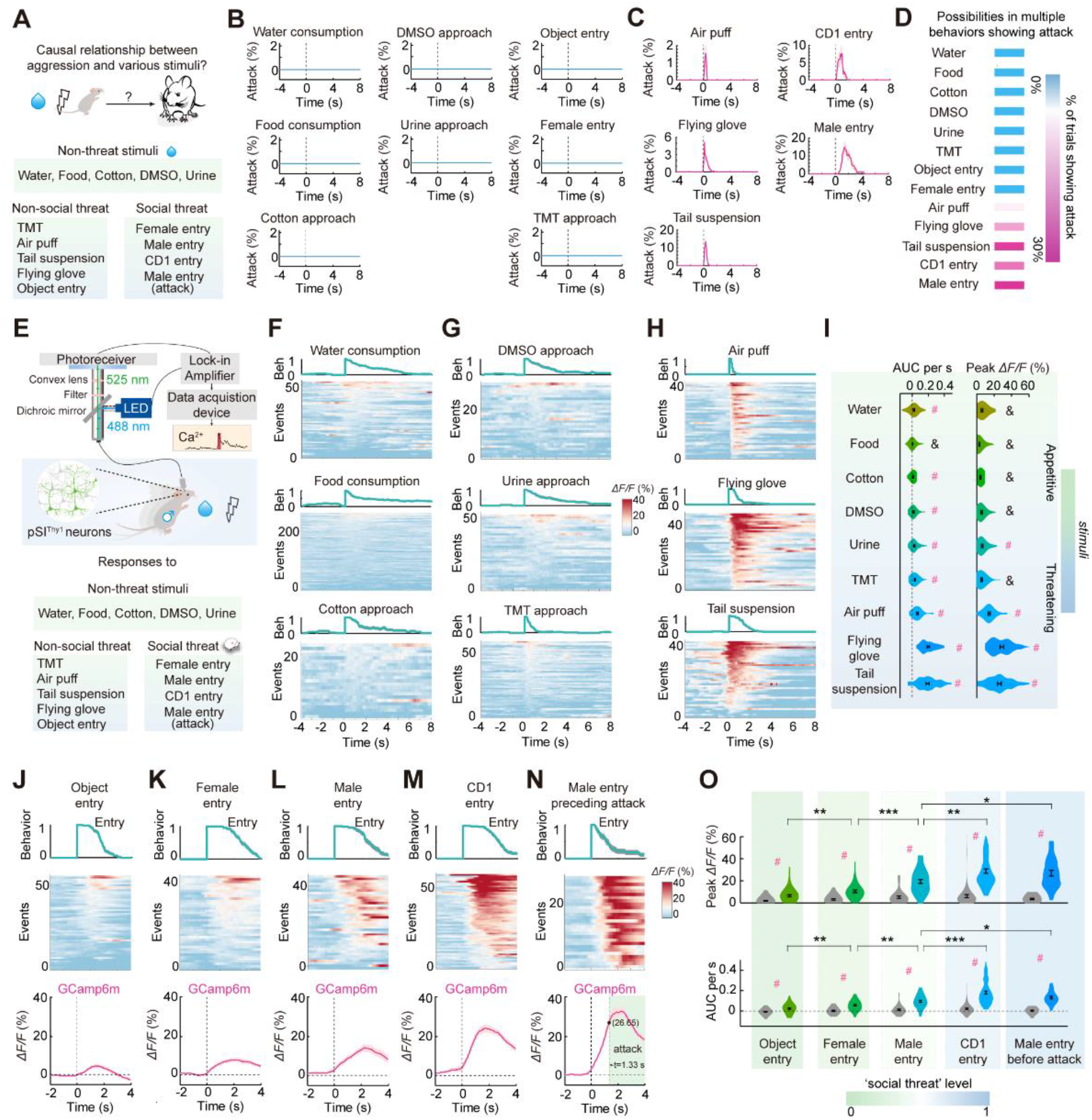
pSI^Thy1^ Neurons Are Differently Modulated by Non-social and Social Contextual Cues, Related to Figure 3 (A) Setup for aggressive behaviors recording in response to diverse non-social and social stimuli. (B) Distribution of episodes of attack behavior during water consumption (n = 52 trials), food consumption (n = 260 trials), approach a cotton ball (n = 23 trials), during detection of DMSO (n = 30 trials), approach a urine cotton ball (n = 36 trials), object entry (n = 39 trials), female C57 entry (n = 39 trials), and approach a TMT cotton ball (n = 57 trials). (C) Distribution of episodes of attack behavior in response to air puff (n = 322 trials), the threat of a flying glove (n = 567 trials), tail suspension (n = 142 trials), male CD-1 entry (n = 119 trials), and male C57 entry (n = 108 trials). (D) Possibilities of behavior trials showing attack during diverse stimuli as shown in (B) and (C). (E) Setup for fiber photometric recording in pSI^Thy1^ neurons in response to diverse non-social and social stimuli. (F) Distribution of episodes of behavior and heatmaps of GCaMP6m Δ*F/F* signals from pSI neurons during water consumption (upper, n = 52 trials), food consumption (middle, n = 260 trials), and approach a cotton ball (lower, n = 23 trials). (G) Distribution of episodes of behavior and heatmaps of GCaMP6m Δ*F/F* signals from pSI neurons during detection of DMSO (upper, n = 30 trials), urine (middle, n = 36 trials), and TMT (lower, n = 57 trials) by approach. (H) Distribution of episodes of behavior and heatmaps of GCaMP6m Δ*F/F* signals from pSI neurons in response to air puff (upper, n = 40 trials), the threat of a flying glove (middle, n = 40 trials), and tail suspension (lower, n = 40 trials). (I) AUC/s (left) and Peak Δ*F/F* (right) of GCaMP6m signals during diverse stimuli. ‘#’ and ‘&’ above each bar indicate that AUC per second (left) or peak Δ*F/F* (right) during each behavior period was (*P* < 0.05) and wasn’t (*P* > 0.05) significantly increased compared with baseline pre-behavior, determined by Wilcoxon matched-pairs signed-rank test and two-tailed paired t-test. (J-N) Distribution of episodes of behavior, heatmaps of GCaMP6m Δ*F/F* signals, and Δ*F/F* of GCaMP6m (magenta) signals from pSI neurons with object entry (J, n = 53 trials in female entry (K, n = 39 trials), male C57 entry (L, n = 39 trials), male CD-1 entry (M, n = 58 trials), and male C57 entry preceding an attack (N, n = 28 trials). (O) Peak Δ*F/F* (upper), and AUC/s (lower) of GCaMP6m signals during different entries. ‘#’ above each bar indicate that peak Δ*F/F* (upper) or AUC per second (lower) during each behavior period was significantly (*P* < 0.001) increased compared with baseline pre-behavior, determined by Wilcoxon matched-pairs signed-rank test and two-tailed paired t-test. The significance of differences in peak Δ*F/F* was determined by the Mann-Whitney and Kruskal-Wallis tests, unpaired t-test, and paired t-test: from left to right, \*\**P* = 0.005, \*\*\**P* < 0.001, \*\**P* = 0.004, \**P* = 0.0131. Significance of differences in the AUC/s was determined by the Mann-Whitney and Kruskal-Wallis tests, unpaired t-test, and paired t-test: from left to right, \*\**P* = 0.007, \*\**P* = 0.0083, \*\*\**P* < 0.001, \**P* = 0.0239. ns, not significant. Data are presented as the mean ± SEM.

**Figure S5.**
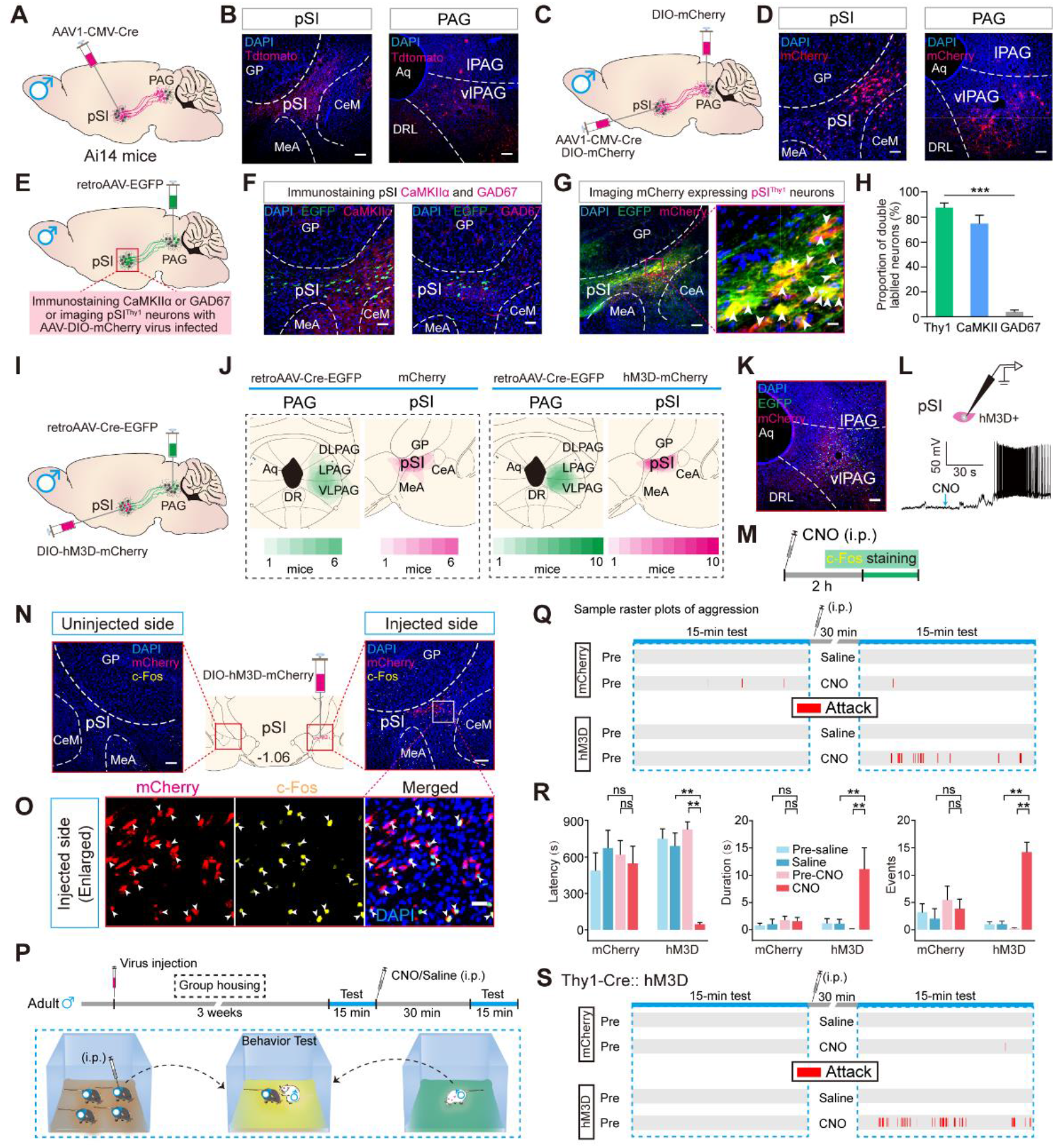
Anterograde Virus Tracing of the pSI-PAG Connection, and Pharmacogenetic Activation of pSI^Thy1^ or pSI^-PAG^ Neurons Promoted Inter-male Aggression, Related to Figure 4 (A) Schematic for injection of anterograde trans-synaptic virus into the pSI to trace pSI-PAG connections in Ai14 male mice. (B) Representative images of pSI (left) and VL/LPAG (right) neurons infected with anterograde trans-synaptic AAV1-CMV-Cre Tdtomato (scale bars, 50 μm). (C) Schematic of the strategy for anterograde virus tracing. (D) Representative images showing the expression of trans-synaptic anterograde AAV1-CMV-Cre and AAV-DIO-mCherry (red) in the pSI (left, scale bar, 50 μm) and the VL/LPAG (right, scale bar, 100 μm). Aq, aqueduct; DRL, dorsal raphe nucleus, lateral part. (E) Schematic showing injection of retrograde tracing virus into the PAG and immunostaining/imaging of pSI^-PAG^ neurons. (F) Example images showing the overlap of retrogradely-labeled pSI^-PAG^ neurons (green) with staining for DAPI (blue) and CaMKIIα (left, red) or GAD67 (right, red). Scale bars, 50 μm. (G) Left panels, example images showing the overlap of retrogradely-labeled pSI^-PAG^ neurons (green) with staining for Thy1 (red). Scale bars, 100 μm. Right panels, magnified images of the boxed areas in the left panels; Scale bars, 10 μm. Arrowheads indicate retrogradely-labeled neurons co-localized with mCherry. (H) Percentage overlap of EGFP-expressing neurons with Thy1, CaMKIIα or GAD67 (KW = 11.65, \*\*\**P* < 0.001; Kruskal-Wallis test, n = 5/5/6 mice). (I) Schematic of the strategy for retrograde tracing from the PAG and Cre-dependent pharmacogenetic activation of pSI^-PAG^ neurons. (J) Left, overlay of EGFP and mCherry expression from mice unilaterally injected with retrograde AAV-Retro-Cre-EGFP virus into the PAG and anterograde AAV-DIO-mCherry virus into the pSI. Right, overlay of EGFP and hM3D-mCherry expression from mice unilaterally injected with retrograde AAV-Retro-Cre-EGFP virus into the PAG and anterograde AAV-DIO-hM3D-mCherry virus into the pSI. (K) Representative image of the PAG showing neurons infected with retrograde AAV-Retro-Cre-EGFP (green) and mCherry terminals (red) projected from the pSI (scale bar, 100 μm). (L) Upper, schematic of whole-cell electrophysiological recordings from hM3D-expressing pSI neurons in slice preparations. Lower, sample recording trace from an hM3D-expressing pSI neuron in response to perfusion with CNO. (M) Timing for the examination of c-Fos expression in the pSI after intraperitoneal injection of CNO. (N) Middle, schematic showing unilateral injection of hM3Dq-mCherry virus into the pSI. Left, representative histological image showing c-Fos expression (yellow) on the uninjected side. Right, representative histological image of c-Fos expression on the side of the pSI injected with hM3Dq-mCherry virus (red). Scale bars, 100 μm. (O) High-magnification images of the boxed area in (N) (scale bar, 10 μm; arrowheads indicate pSI neurons immunopositive for c-Fos antibody). (P) Behavioral paradigm of aggression test. (Q) Raster plots of attack behavior recorded during 15-min encounters. (R) Summary of aggression during the pre- and post-injection phases in the control and hM3Dq groups (n = 7/10 mice) with saline or CNO injection. Red blocks indicate duration of attack episodes. Pre-CNO *vs* CNO in the hM3D group: from left to right, \*\**P* = 0.002, \*\**P* = 0.002, \*\**P* = 0.002; saline *vs* CNO in the hM3D group: from left to right, \*\**P* = 0.002, \*\**P* = 0.002, \*\**P* = 0.002, Wilcoxon matched-pairs signed-rank test. (S) Raster plots of attack behavior recorded during 15-min encounters. ns, not significant. Data are presented as the mean ± SEM.

**Figure S6.**
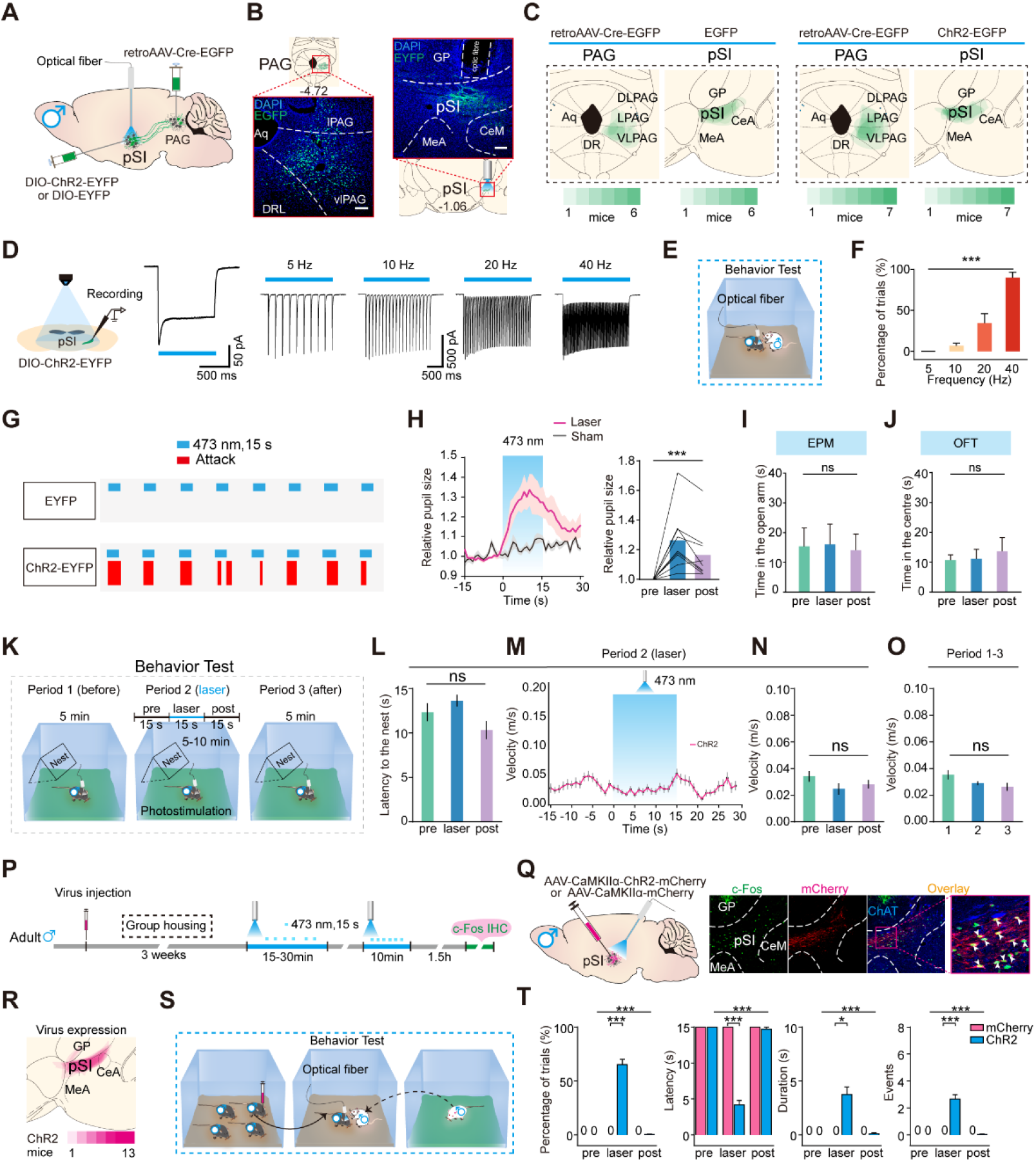
Photoactivation of pSI^-PAG^ Neurons or pSI^CaMKIIα^ Neurons Time-locked to Evoked Inter-male Aggression and Autonomic Arousal, but not Anxiety, Related to Figure 4 (A) Strategy for virus injection and photoactivation of pSI^-PAG^ neurons in male mice. (B) Schematics and representative images of AAV-Retro-Cre-EGFP expression in the PAG (left, scale bar, 100 μm) and AAV-DIO-ChR2-EYFP expression in upstream pSI neurons (right, scale bars, 50 μm). (C) Left, overlay of EGFP expression from mice unilaterally injected with retrograde AAV-Retro-Cre-EGFP virus into the PAG and anterograde AAV-DIO-EGFP virus into the pSI. Right, overlay of EGFP and ChR2-EGFP expression from mice unilaterally injected with retrograde AAV-Retro-Cre-EGFP virus into the PAG and anterograde AAV-DIO-ChR2-EGFP virus into the pSI. (D) Left, schematic of whole-cell recordings from ChR2-expressing pSI^-PAG^ neurons in slice preparations. Middle, the light-induced inward current recorded in a ChR2+ pSI neuron (blue bar, application of a LED-generated blue light pulse). Right, action potentials recorded from a ChR2-expressing pSI neuron induced by 5, 10, 20, and 40 Hz light stimulation. (E) Behavioral paradigm of inter-male aggression test. (F) Success rates of light-induced attacks in response to different frequencies of light stimulation at a fixed intensity (2.7 mW). 5 Hz *vs* 10 Hz *vs* 20 Hz *vs* 40 Hz in the ChR2 group: 0.0 ± 0.0 *vs* 6.7 ± 3.3 *vs* 34.4 ± 11.7 *vs* 90.0 ± 6.7 (%), \*\*\**P* < 0.001, n = 9 trials, Friedman test. (G) An example of behavioral output during photostimulation in the ChR2 and control groups. (H) Left, peristimulus time histogram of relative pupil size (pupil size/eye size) aligned with light onset in the laser or sham stimulation group (n = 9/8 mice). Right, relative pupil size before, during, and after photostimulation (\*\*\**P* < 0.001, n = 9 mice, Friedman test). (I) Average percentage of time in the open arms of the elevated plus maze before, during, and after photostimulation of pSI^-PAG^ neurons (Friedman test, FM = 0.7429, *P* = 0.7407, n = 9 mice). (J) Average percentage of time in the center of the open field before, during, and after photostimulation of pSI^-PAG^ neurons (one-way ANOVA, F (1.608, 12.87) = 0.2183, *P* = 0.7599, n = 9 mice). (K) Behavioral paradigm of open field test with photoactivation of pSI^-PAG^ neurons in socially-housed male C57 mice (n = 6 mice), with a shield cover (nest) inside the open field box. (L-N) Latency to the nest (L, *P* = 0.0721, Friedman test), averaged velocity in each photostimulation period (M, from −15 s to 30 s), and velocity in the pre (1), during (2), and post-laser (3) period (N, *P* = 0.3379, one-way ANOVA). (O) Averaged velocity during period 1, 2, and 3 (*P* = 0.0875, one-way ANOVA). (P) Schematic of the timing and behavioral paradigm with optical activation of pSI^CaMKIIα^ neurons in male mice. (Q) Left, schematic of pSI injection of virus-carrying AAV-CaMKIIα-ChR2-mCherry or AAV-CaMKIIα-mCherry and light stimulation of pSI^CaMKIIα^ neurons. Right, representative histological images of light-induced c-Fos expression (green) in neurons expressing AAV-CaMKIIα-ChR2-mCherry (magenta). Arrowheads indicate neurons co-expressing AAV-CaMKIIα-ChR2-mCherry and c-Fos. (R) Overlay of CaMKIIα-ChR2-mCherry expression in pSI (−1.06 mm from bregma). (S) Behavioral paradigm of a socially-housed male mouse in its home cage attacking a singly-housed male intruder. (T) Percentage of trials with attacks, total latency to attack, attack duration, and number of attack events in response to pSI photostimulation in the ChR2 group and controls (mCherry group) (Wilcoxon matched-pairs signed-rank test and two-tailed unpaired t-test: from left to right, \*\*\**P* < 0.001, \*\*\**P* < 0.001, \**P* = 0.0313, \*\*\**P* < 0.001, n = 15/6 mice) during the pre-laser, laser, and post-laser phases when a socially-housed male mouse initiated an attack in its home cage on a singly-housed male intruder (Friedman test: from left to right, \*\*\**P* < 0.001, \*\*\**P* < 0.001,\*\*\**P* < 0.001, \*\*\**P* < 0.001, n = 15 mice). ns, not significant. Data are presented as the mean ± SEM.

**Figure S7.**
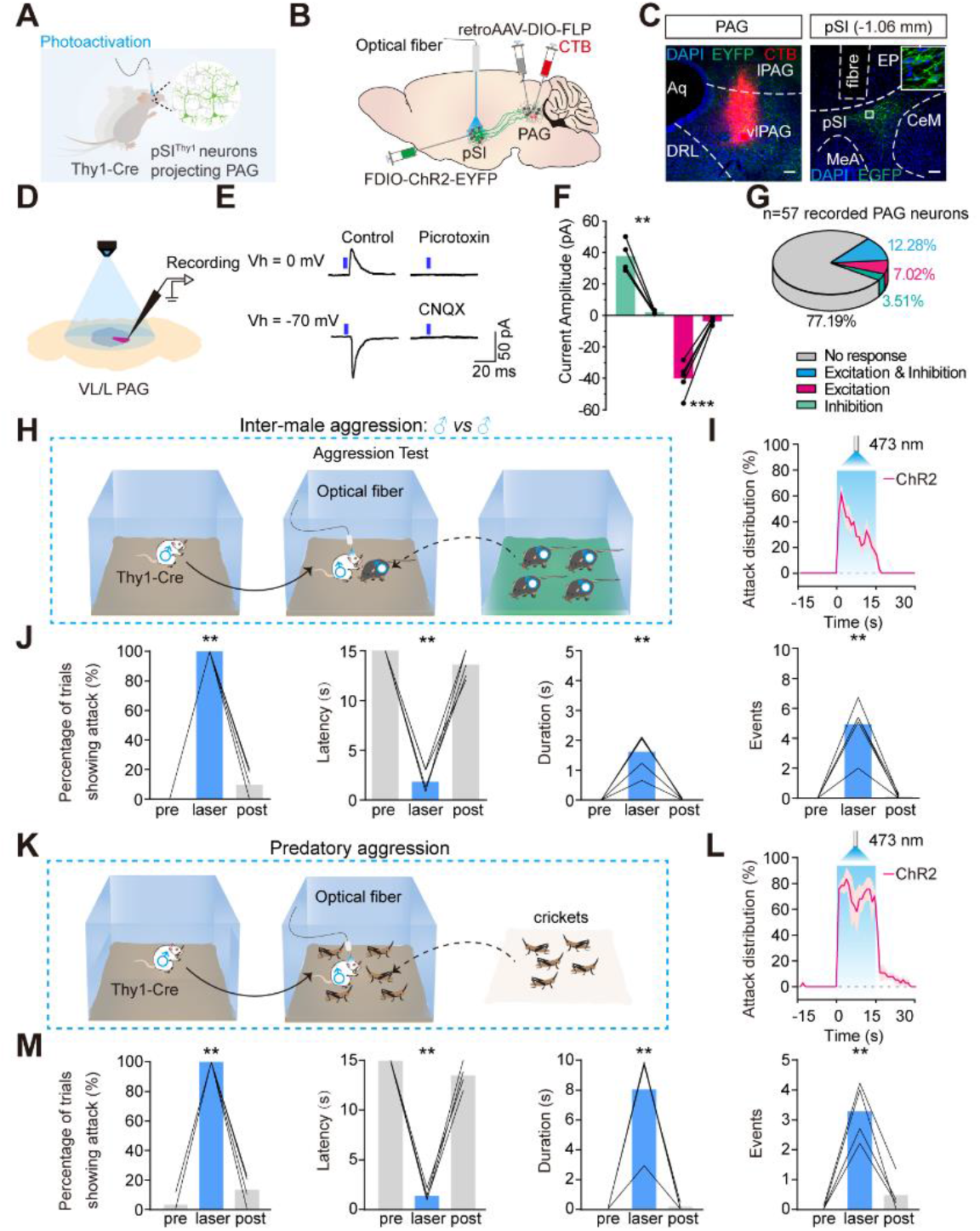
Optogenetic Activation of pSI^Thy1^ Neurons Projecting to the PAG Promotes Inter-male Aggression, Related to Figure 4 (A, B) Experimental design: (A) photostimulation of pSI^Thy1^ neurons projecting to the PAG and (B) schematic of the injection strategy of virus and CTB555. (C) Left, representative image of retrograde AAV-CAG-retro-FLEX-Flpo and CTB555 infection (red) in the PAG (scale bar, 100 μm); Aq, aqueduct; DRL, dorsal raphe nucleus, lateral part; MeA, medial amygdaloid nucleus; CeM, central amygdaloid nucleus, medial division. Right, representative image of FDIO-ChR2-EYFP (green) and ChAT (yellow) in the pSI (scale bar, 100 μm). Insets, magnified images of the boxed areas in the right panels; scale bar, 10 μm. (D) Slice electrophysiological recording of the activity of VL/LPAG neurons induced by photoactivation of pSI^Thy1^ projection terminals in the VL/LPAG. (E) Representative traces of the light-evoked currents in the PAG neurons by photostimulating axonal projections from the pSI to VL/LPAG neurons in the absence or presence of the GABA receptor antagonist Picrotoxin and the glutamate receptor antagonist CNQX; Vh, holding voltage. (F) Amplitudes of light-evoked excitatory and inhibitory postsynaptic currents in the absence (left) or presence (left) of Picrotoxin and CNQX (from left to right, \*\**P =* 0.0043, \*\*\**P =* 0.0006, n = 4/5 neurons). (G) Proportions of PAG neurons (n = 57) showing light-evoked excitatory and inhibitory postsynaptic currents by photostimulating axonal projections from the pSI. (H) Schematic of inter-male aggression test when a singly-housed male Thy1-Cre mouse encounters a socially-housed male C57 mouse in its home cage. (I) Distribution of attack episodes during photostimulation. (J) Summary of male aggression induced by photostimulation of pSI^Thy1^ neurons projecting to the PAG: percentage of trials showing attacks (\*\**P* = 0.0031), latency to attack onset (\*\**P* = 0.0031), attack duration (\*\**P* = 0.0031), and attack events (\*\**P* = 0.0031) in pre-laser, laser, and post-laser phases. n = 5 mice, Friedman test. (K) Schematic of the predatory aggression test when a singly-housed male Thy1-Cre mouse encounters crickets in its home cage. (L) Distribution of attack episodes during photostimulation. (M) Summary of predatory aggression induced by photostimulation of pSI^Thy1^ neurons projecting to the PAG: percentage of trials showing attacks, latency to attack onset, attack duration, and attack events (\*\**P* = 0.0093 for all) in pre-laser, laser, and post-laser phases, n = 4 mice, Friedman test. Data are presented as the mean ± SEM.

**Figure S8.**
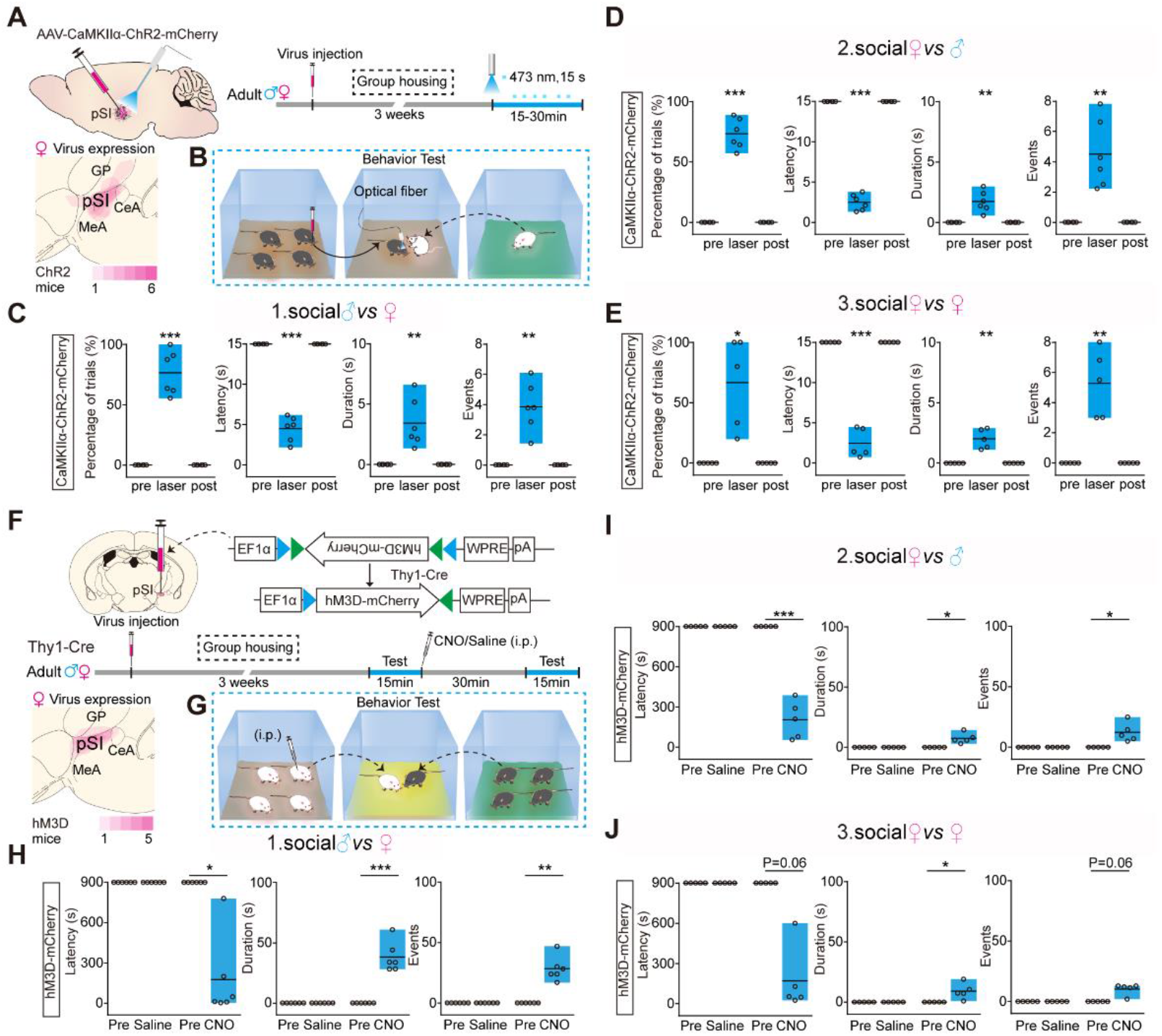
Optical Activation of pSI^CaMKIIα^ Neurons or Pharmacogenetic Activation of pSI^Thy1^ Neurons Evoked Aggressive Behaviors under Various Conditions, Related to Figure 5 (A) Upper left, schematic of pSI injection of virus-carrying AAV-CaMKIIα-ChR2-mCherry and optical activation of pSI^CaMKIIα^ neurons. Upper right, schematic of the timing and behavioral paradigm with optical activation of pSI^CaMKIIα^ neurons in male or female mice. Lower, overlay of CaMKIIα-ChR2-mCherry expression in the pSI (–1.06 mm from bregma) of female mice. (B) Behavioral paradigm of a socially-housed mouse in its home cage attacking a singly-housed intruder induced by optical activation of pSI^CaMKIIα^ neurons. (C-E) Percentage of trials with attacks, latency to attack onset, attack duration, and attack events with photostimulation of pSI^CaMKIIα^ neurons in the pre-laser, laser, and post-laser phases. A socially-housed male mouse in its home cage initiated an attack on a socially-housed female intruder (C, from left to right, \*\*\**P* < 0.001, \*\*\**P* < 0.001, \*\**P* = 0.0092, \*\**P* = 0.0022, n = 6 mice), and a socially-housed female in its home cage initiated an attack on a socially-housed male (D, from left to right, \*\*\**P* < 0.001, \*\*\**P* < 0.001, \*\**P* = 0.0046, \*\**P* = 0.0047, n = 6 mice) or female intruder (E, from left to right, \**P* = 0.0168, \*\*\**P* < 0.001, \*\**P* = 0.005, \*\**P* = 0.0063, n = 5 mice). Significance was determined by one-way ANOVA. (F) Upper, schematic of the timing and behavioral paradigm with pharmacogenetic activation of pSI^Thy1^ neurons through Cre-dependent expression of AAV-DIO-hM3d-mCherry in Thy1-Cre male and female mice. Lower, overlay of hM3d-mCherry expression in pSI (−1.06 mm from bregma) of the female Thy1-Cre mice. (G) Behavioral paradigm in which the pharmacogenetic activation of pSI^Thy1^ neurons in a socially-housed Thy1-Cre male mouse in a novel cage promoted attacks on a socially-housed female. (H-J) Latency to attack, attack duration, and attack events during the pre- and post-injection phases of the pharmacogenetic activation of pSI^Thy1^ neurons. A socially-housed male mouse encountered a socially-housed female (H, from left to right, \**P* = 0.0313, \*\*\**P* < 0.001, \*\**P* = 0.001, n = 6 mice) and a socially-housed female encountered a socially-housed male (I, from left to right, \*\*\**P* < 0.001, \**P* = 0.0176, \**P* = 0.0264, n = 5 mice) or female (J, from left to right, *P* = 0.0625, \**P* = 0.0378, *P* = 0.0625, n = 5 mice) in a novel cage. Significance was determined by the Wilcoxon matched-pairs signed-rank test and two-tailed paired t-test. ns, not significant. Data are presented as the mean ± SEM.

**Figure S9.**
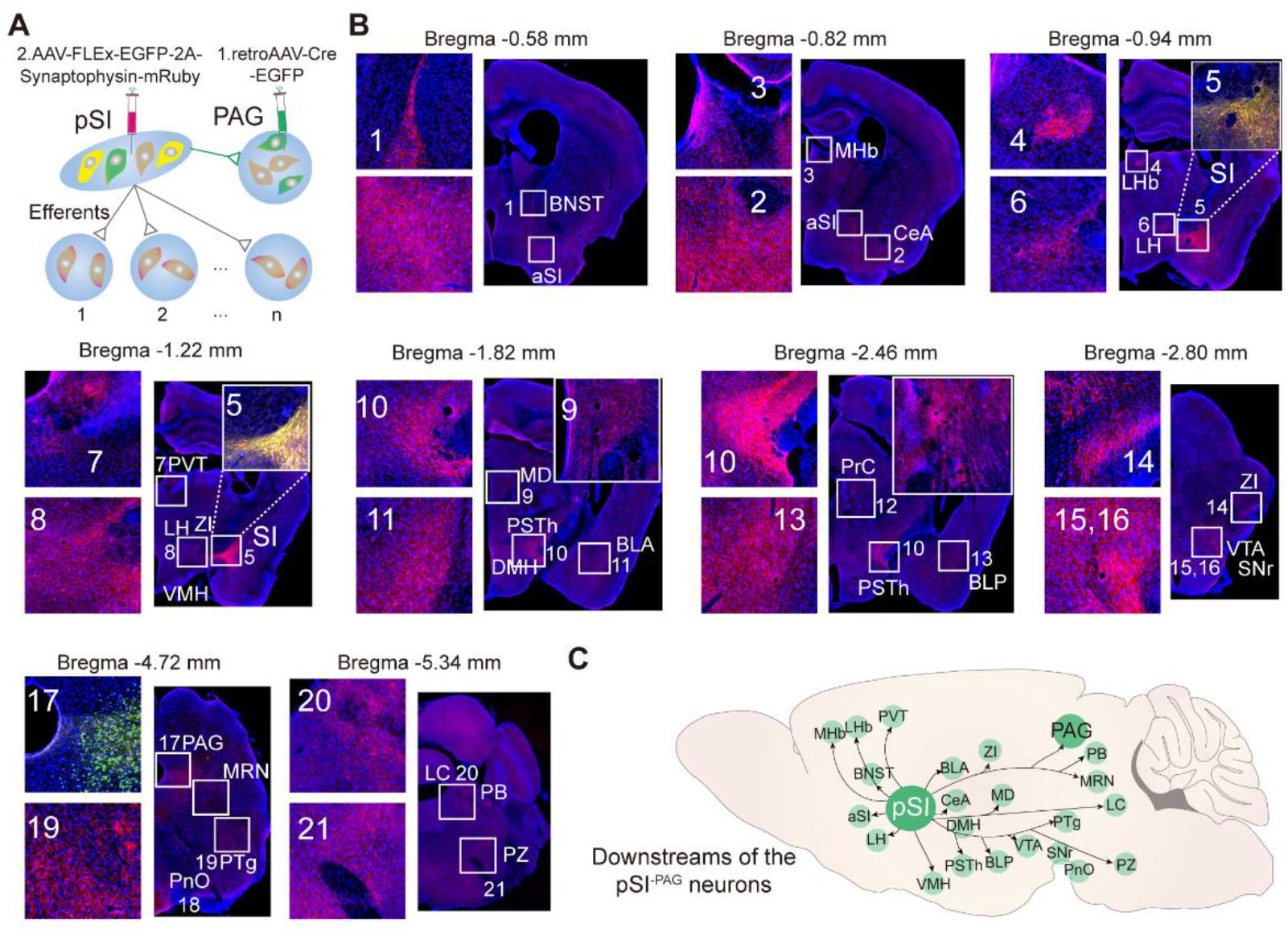
Parallel Output Connections of pSI^-PAG^ Neurons, Related to Figure 5 (A) Strategy of anterograde tracing of pSI^-PAG^ neurons. (B) Horizontal section series showing the brain-wide distribution of the output connected neurons with pSI^-PAG^ neurons revealled by anterograde transsynaptic tracing experiments. Green, EGFP; magenta, mCherry from mRuby. (C) Overview of the outputs from the pSI^-PAG^ neurons as shown in (B).

**Figure S10.**
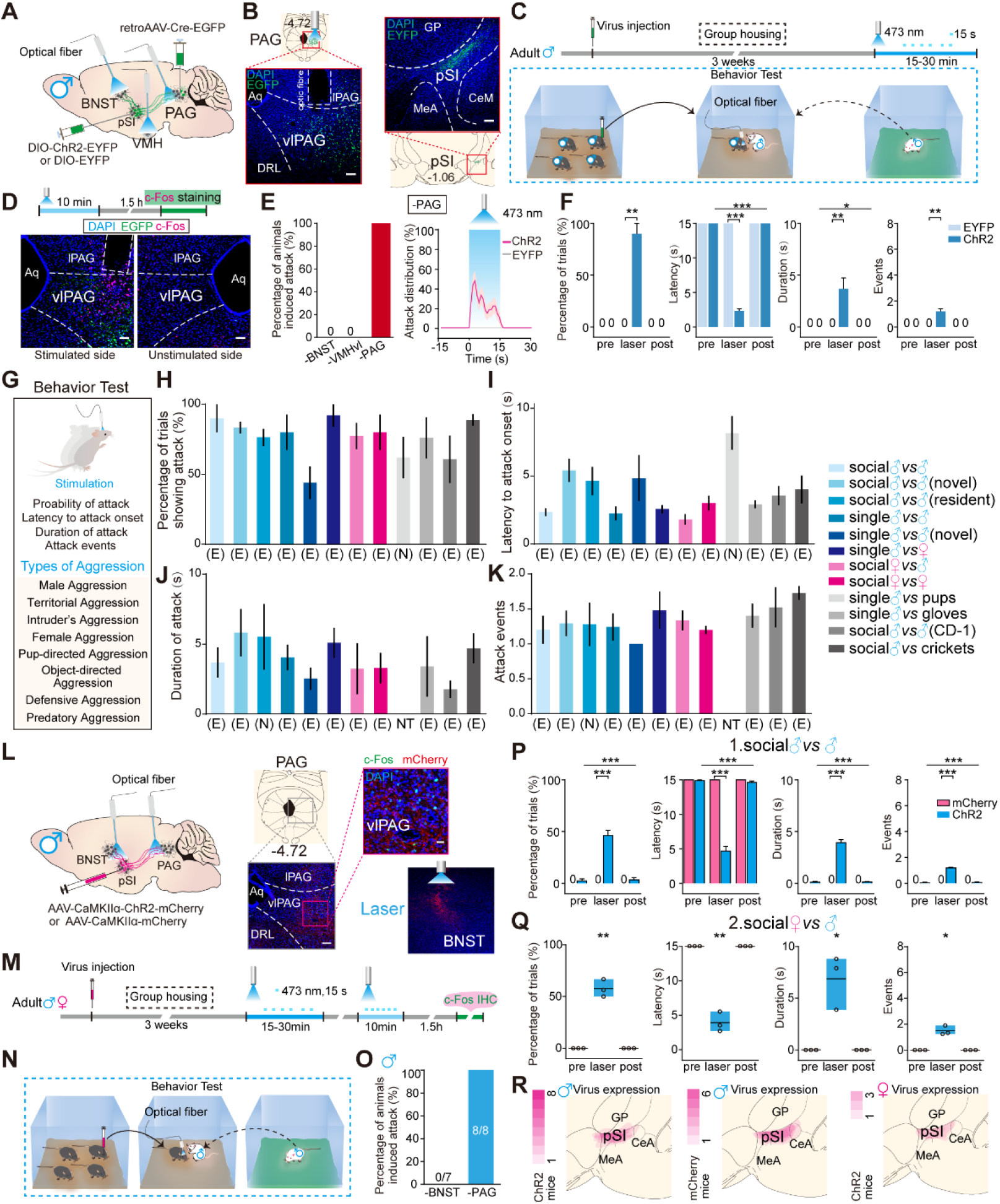
The Projection from pSI^CaMKIIα^ Neurons to the PAG, but not to the BNST or VMH, Universally Drove Diverse Aggressive Behaviors, Related to Figure 5 (A) Strategy for virus injection and optical activation of the terminals of pSI^-PAG^ neurons in the PAG and BNST in male mice. (B) Schematics and representative images of AAV-Retro-Cre-EGFP expression in the VL/LPAG (left; scale bar, 100 μm) and AAV-DIO-ChR2-EYFP expression (green) upstream from pSI neurons (right; scale bar, 100 μm). (C) Behavioral paradigm of the attack test of a socially-housed male in its home cage towards a singly-housed male intruder. (D) Upper, schematic of c-Fos examination of PAG neural activation induced by light stimulation of pSI^-PAG^ terminals in male mice. Lower, representative histological images of the PAG stained with c-Fos (red) after photostimulation of pSI^-PAG^ terminals expressing ChR2-EGFP (green) on the stimulated and unstimulated sides (scale bars, 50 μm). (E) Left, percentage of animals showing attacks by photostimulation of the pSI-BNST, pSI-VMH, and pSI-PAG projection in male mice. Right, distribution of attack episodes during photostimulation of the pSI-PAG projection (magenta, ChR2 group; gray, EYFP group; n = 5 mice). (F) Percentage of trials, latency to attack, attack duration, and attack events in the photostimulation (ChR2) and control (EYFP) groups during the pre-laser, laser, and post-laser phases. Pre *vs* Laser *vs* Post in the ChR2 group: *P* = 0.123, \*\*\**P* < 0.001, \**P* = 0.0271, *P* = 0.123, one-way ANOVA or Friedman test, n = 5 mice; Laser in EYFP and the ChR2 group: \*\**P* = 0.0022, \*\*\**P* < 0.001, \*\**P* = 0.0044, \*\**P* = 0.0022, two-tailed unpaired t-test and Mann-Whitney test, n = 5/6 mice. (G) Schematic of the behavioral design to test various forms of aggression induced by photostimulation of pSI^-PAG^ projection terminals in the PAG under various conditions. (H-K) Aggression probability (H), latency to attack onset (I), duration (J), and events (K) of various forms of aggression induced by photostimulation of pSI^-PAG^ projection terminals in the PAG. ‘(E)’ and ‘(N)’ under the x-axis indicate that the aggression during photostimulation was significantly increased (E) or not changed (N), determined by one-way ANOVA or the Friedman test. ‘NT’ under the x-axis indicates that this index of aggression was not tested under this condition. (L) Left, schematic of Cre-dependent expression of ChR2-mCherry or mCherry in pSI^CaMKIIα^ neurons and optogenetic activation of the terminals of pSI^CaMKIIα^ neurons projecting to the PAG or BNST. Middle, schematic and example images of c-Fos expression (green) in the PAG induced by local light stimulation of pSI^CaMKIIα^ neuronal terminals in the PAG. Right, example image of pSI^CaMKIIα^ neuronal terminals in the BNST. (M) Schematic of the timing and behavioral paradigm with optical activation of pSI^CaMKIIα^ neuron terminals in a socially-housed mouse in its home cage encountering a male intruder. (N) Behavioral paradigm of a socially-housed mouse in its home cage attacking a singly-housed intruder induced by optical activation of pSI^CaMKIIα^ neuronal terminals. (O) Summary data showing attacks were induced by optogenetic activation of pSI^CaMKIIα^ terminals in the PAG but not in the BNST in socially-housed male mice. (P) Summary data showing socially-housed male mice in their home cages initiated attacks on singly-housed male intruders during optical activation of the pSI^CaMKIIα^ -PAG projection. Percentage of trials with attacks, latency to attack, attack duration, and attack events induced by optogenetics during the pre-laser, laser, and post-laser phases in the control mCherry group (n = 6 mice) and in the ChR2 group (n = 8 mice). Pre *vs* Laser *vs* Post in the ChR2 group: \*\*\**P* < 0.001, \*\*\**P* < 0.001, \*\*\**P* < 0.001, \*\*\**P* < 0.001, Friedman test, n = 8 mice; Laser in the mCherry and ChR2 groups: \*\*\**P* < 0.001, \*\*\**P* < 0.001, \*\*\**P* < 0.001, \*\*\**P* < 0.001, two-tailed unpaired t-test, n = 6/8 mice. Significance was determined by the Friedman test and unpaired t-test. (Q) Summary data showing socially-housed female mice in their home cages initiated attacks on singly-housed male intruders during optical activation of the pSI^CaMKIIα^-PAG projection (\*\**P* = 0.007, \*\**P* = 0.0056, \**P* = 0.0451, \**P* = 0.0182, n = 3 mice, significance determined by one-way ANOVA). (R) Overlays of CaMKIIα-ChR2-mCherry (left) and CaMKIIα-mCherry (middle) expression in pSI in male mice, Right, overlay of CaMKIIα-ChR2-mCherry expression in pSI of female mice. ns, not significant. Data are presented as the mean ± SEM.

**Figure S11.**
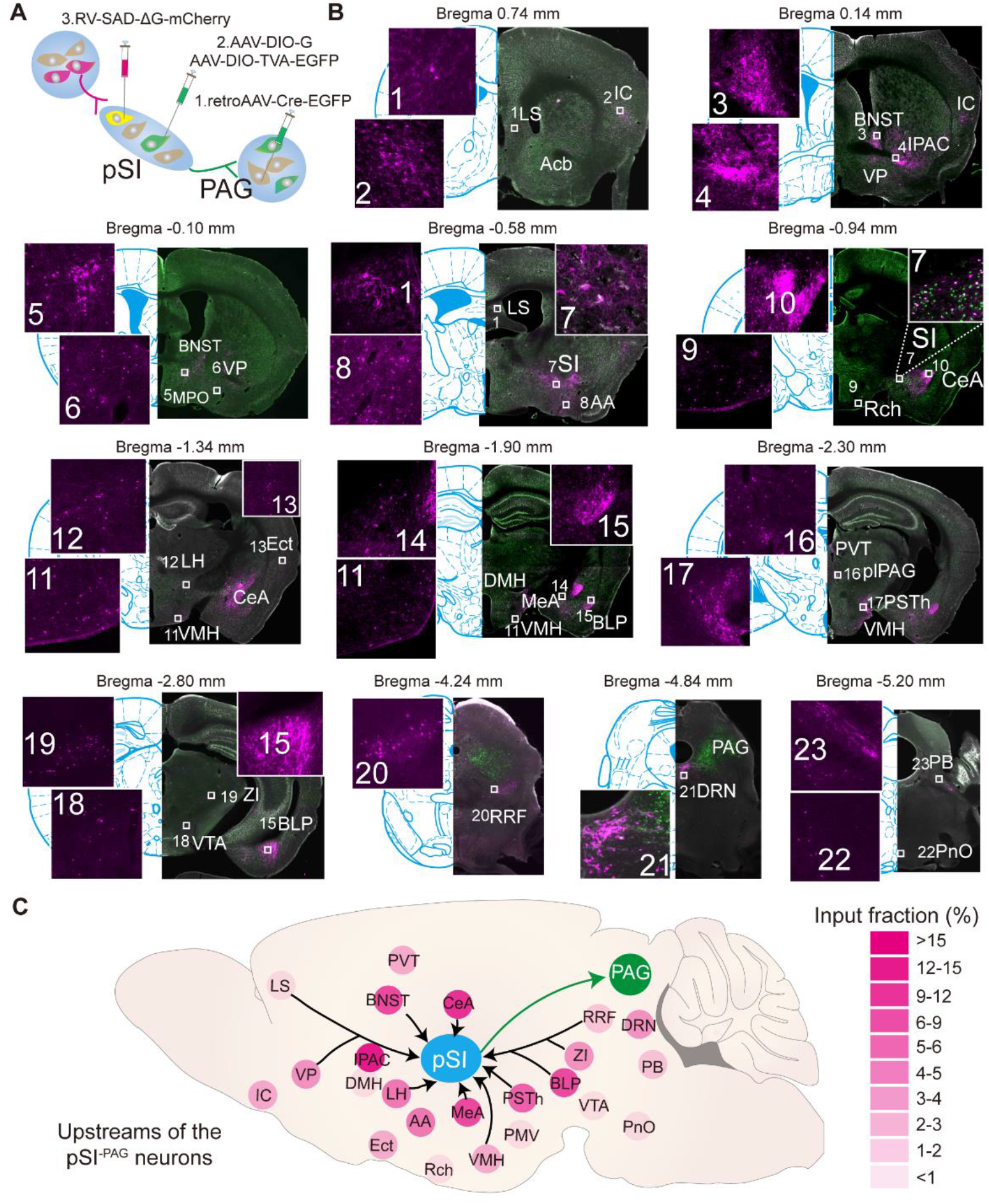
Inputs of pSI^-PAG^ Neurons, Related to Figure 6 (A) Strategy of retrograde transsynaptic tracing of pSI^-PAG^ neurons. (B) Horizontal section series showing the brain-wide distribution of inputs to pSI^-PAG^ neurons revealed by the retrograde transsynaptic tracing experiments. Green, EGFP; magenta, mCherry from RVdG. (C) Overview of the inputs into the pSI^-PAG^ neurons as shown in (B). Right is the quantification of inputs to pSI^-PAG^ neurons, Data are shown as the percentage of inputs from a given region relative to total inputs throughout a given brain (n = 3 mice). Heat map coloration corresponds to the percentage of total labeling

**Figure S12.**
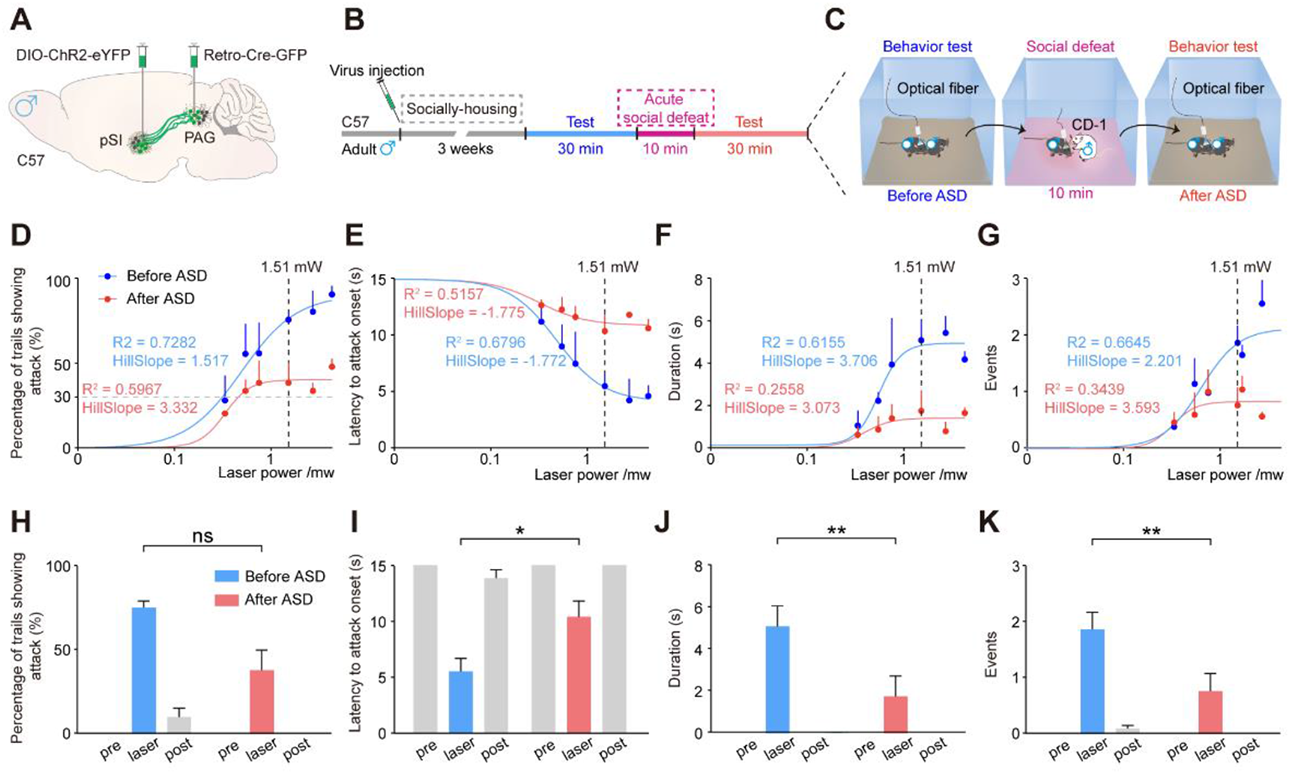
Effect of ASD on the Aggressive Activity Triggered by Photostimulation of pSI^-PAG^ Neurons, Related to Figure 6 (A) Strategy for virus injection and optical activation of pSI^-PAG^ neurons in male mice. (B) Schematic for testing inter-male aggression induced by different stimulation intensities before and after ASD. (C) Behavioral paradigm of inter-male aggression during photoactivation of pSI^-PAG^ neurons at different stimulation intensities before and after ASD. (D-G) Nonlinear regression of the probability of trials showing attack (D), latency to attack onset (E), duration (F), and events (G) of the inter-male aggression triggered by photoactivation of pSI^- PAG^ neurons at different intensities. Different colored lines are non-linear fits before and after ASD (pooled across all mice and binned light intensities). (H-K) Probability of trials showing attack (H), latency to attack onset (I), duration (J) and events (K) of the inter-male aggression induced during laser stimulation with a specific laser intensity (1.51 mW) before and after ASD (n = 3 mice). Before ASD *vs* After ASD: from left to right, \**P =* 0.0306, \*\**P =* 0.0045, \*\**P =* 0.0022, *P =* 0.2500, two-tailed paired t-test and Wilcoxon matched-pairs signed-rank test. ns, not significant. Data are presented as the mean ± SEM.

**Figure S13.**
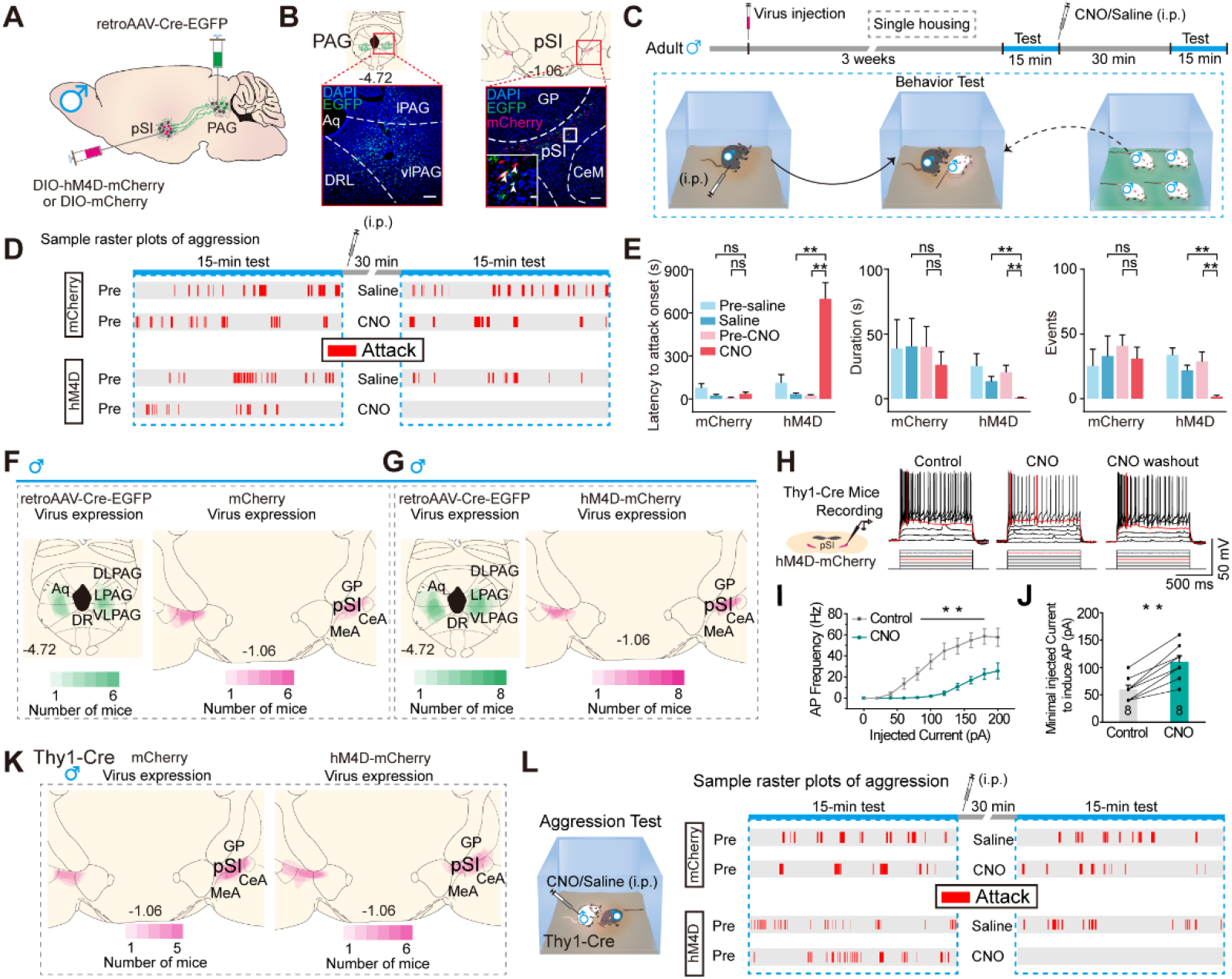
pSI^-PAG^ and pSI^Thy1^ Neurons Are Necessary for Inter-male Aggression, Related to Figure 7 (A) Schematic of the expression of AAV-Retro-Cre-EGFP in the PAG to drive Cre-dependent expression of AAV-DIO-hM4Di-mCherry in pSI^-PAG^ neurons. (B) Left, schematic and representative image showing the expression of retrograde AAV-Retro-Cre-EGFP (green) in the PAG (scale bar, 100 μm). Right, schematic and representative image showing Cre-dependent expression of AAV-DIO-hM4Di-mCherry (red) in pSI^-PAG^ neurons infected with AAV-Retro-Cre-EGFP (green, arrowheads) (scale bars, 100 μm; inset, 10 μm). (C) Schematic of the timing (upper) and behavioral paradigm (lower) for pharmacogenetic inhibition of pSI^-PAG^ neurons of a singly-housed male mouse in its home cage encountering a socially-housed male intruder (lower). (D) Example raster plots of behavioral recordings from mice during 15-min encounters in the control group (mCherry, n = 6 mice) and the hM4Di group (n = 8 mice) (red marks, episodes of aggression). (E) Latency to attack (left), attack duration (middle), and attack events (right) during the pre-and post-injection phases in control and hM4Di groups after saline or CNO injection. Latency, W = 13.00, \*\**P* < 0.01; duration, W = –36.00, \*\**P* < 0.01; and events, W = –36.00, \*\**P* < 0.01 in the hM4Di group (Wilcoxon matched-pairs signed-rank test and two-tailed paired t-test, n = 8 mice). (F) Overlays of EGFP and mCherry expression from mice bilaterally injected with retrograde AAV-Retro-Cre-EGFP virus into the PAG and anterograde AAV-DIO-mCherry virus into the pSI. (G) Overlays of EGFP and hM4D-mCherry expression from mice bilaterally injected with retrograde AAV-Retro-Cre-EGFP virus into the PAG and anterograde AAV-DIO-hM4D-mCherry virus into the pSI. (H) Left, representative whole-cell current clamp recordings from a pSI neuron (upper) in response to intracellular current injection of a pulse train from 0 pA to 120 pA in a step of 20 pA (lower) before, during, and after 5 μM CNO perfusion. Right, red traces indicate the minimal current to induce action potentials. (I) Summarized response curves showing the number of induced action potentials at different injected current steps in the CNO and control group. Paired t-test, \*\**P* = 0.003, n = 8 neurons. (J) Minimal injected current to induce action potential (APs) in CNO and control group. Paired t-test, \*\**P* = 0.002, n = 8 neurons. (K) Overlays of mCherry (left) and hM4d-mCherry expression (right) in male Thy1-Cre mice with AAV-DIO-mCherry and AAV-DIO-hM4D-mCherry virus bilaterally injected into the pSI. (L) Left, aggression behavioral paradigm. Right, example raster plots of attack behavior recorded during 15-min encounters. ns, not significant. Data are presented as the mean ± SEM.

**Figure S14.**
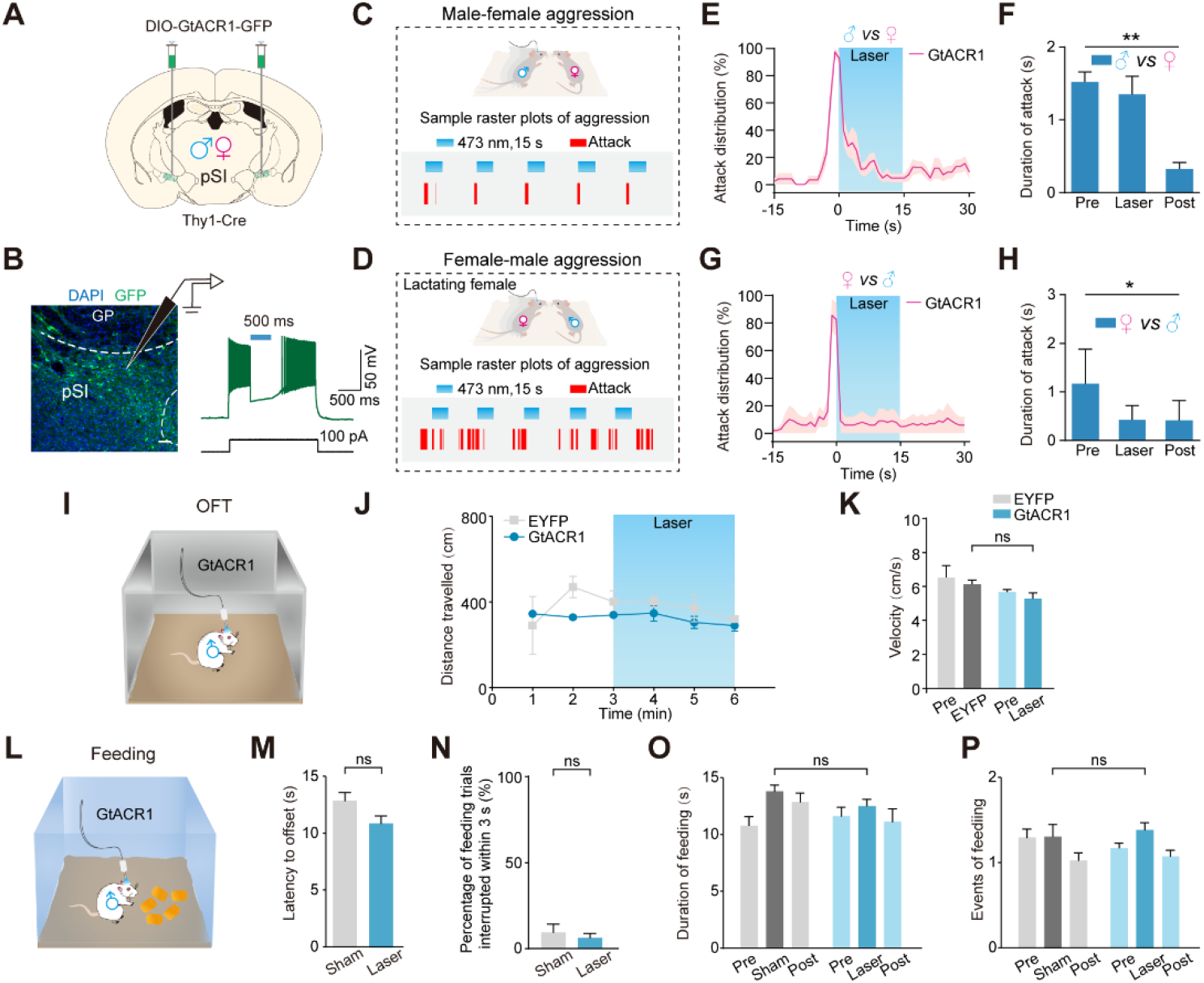
Inactivation of pSI^Thy1^ Neurons Interrupts Female-male and Male-female Aggression, Related to Figure 7 (A) Schematic of bilateral injection of DIO-GtACR1-GFP virus into the pSI of male and female Thy1-Cre mice. (B) Left, example image showing the expression of DIO-GtACR1-GFP in pSI^Thy1^ neurons (scale bar, 50 μm). Right, light-induced inhibition of action potentials (upper green trace) evoked by intracellular current injection (lower black trace) in a GtACR1-expressing pSI^Thy1^ neuron (blue bar, application of a LED-generated blue light pulse; n = 8 neurons). (C, D) Upper, behavioral paradigms to assess the effects of optogenetic inhibition of pSI^Thy1^ neurons on male-female aggression (C) and female-male aggression (D). Lower, example raster plots showing the time-locked interruption of these two types of aggressive behaviors by optogenetic inhibition of pSI^Thy1^ neurons. (E) Distribution of attack episodes interrupted by optogenetic inhibition of pSI^Thy1^ neurons in male-female aggression (n = 6 mice). (F) Attack duration in the pre-laser, laser, and post-laser phases by optogenetic inhibition of aggressive behavior; \*\**P* = 0.0038. (G) Distribution of attack episodes interrupted by optogenetic inhibition of pSI^Thy1^ neurons in female-male aggression (n = 4 lactating adult female mice). (H) Attack duration in the pre-laser, laser, and post-laser phases by optogenetic inhibition of aggressive behavior; \**P* = 0.0417. (I) Behavioral paradigm to test the effect of optogenetic inhibition of pSI^Thy1^ neurons on general locomotion in the open field test (OFT). (J, K) Inhibition of pSI^Thy1^ neurons has no effect on the distance traveled (J) and velocity (K) of locomotor activity in the open field (n = 3/8 mice). EYFP *vs* Laser: *P =* 0.1167. (L) Behavioral paradigm for assessing feeding behavior in response to optogenetic inhibition of pSI^Thy1^ neurons. (M-P) Latency to feeding offset (M), probability of feeding trials interrupted within 3 s (N), duration of feeding (O), and feeding events (P) of the feeding behavior before, during, and after optical silencing of pSI^Thy1^ neurons with a laser intensity the same as that inhibiting attacks (n = 8/8 mice). Sham *vs* Laser: from left to right, *P =* 0.0621, *P =* 0.0621, *P =* 0.0719, *P =* 0.6094. ns, not significant. Data are presented as the mean ± SEM.

**Supplemental Movies S1-S5**

**Movie S1. Optogenetic activation of pSI^-PAG^ neurons in a C57 male mouse in its home cage evokes attacks on a singly-housed male intruder, Related to Figure 4**.

The male mouse (black) transfected with ChR2 in pSI^-PAG^ neurons and implanted with a fiber-optic cable in the pSI was photostimulated during the period indicated by “Light on”.

**Movie S2. Optogenetic activation of pSI^-PAG^ neurons evokes arousal responses, including an enlarged pupil in a head-fixed C57 male mouse, Related to Figure 4.**

The male mouse (black) transfected with ChR2 in pSI^-PAG^ neurons and implanted with a fiber-optic cable in the pSI was photostimulated during the period indicated by “Light on”.

**Movie S3. Optogenetic activation of pSI^-PAG^ neurons in a C57 female mouse in its home cage evokes attack on a socially-housed male intruder, Related to Figure 5.**

The female mouse (black) transfected with ChR2 in pSI^-PAG^ neurons and implanted with a fiber-optic cable in pSI was photostimulated during the period indicated by “Light on”.

**Movie S4. Optogenetic activation of pSI^-PAG^ neurons in a C57 male mouse evokes defensive attack on a male CD-1 mouse in the latter’s cage, Related to Figure 5.**

The male mouse (black) transfected with ChR2 in pSI^-PAG^ neurons and implanted with a fiber-optic cable in pSI was photostimulated during the period indicated by “Light on”.

**Movie S5. Optogenetic silencing of pSI^Thy1^ neurons in a male Thy1-Cre mouse interrupts a naturally-occurring attack on a socially-housed mouse in a time-locked manner, Related to Figure 7.**

The Thy1-Cre male mouse (white) transfected with GtACR1 in pSI^Thy1^ neurons and implanted with a fiber-optic cable in pSI was photostimulated during the period indicated by “Light on”.

**Table S1.**
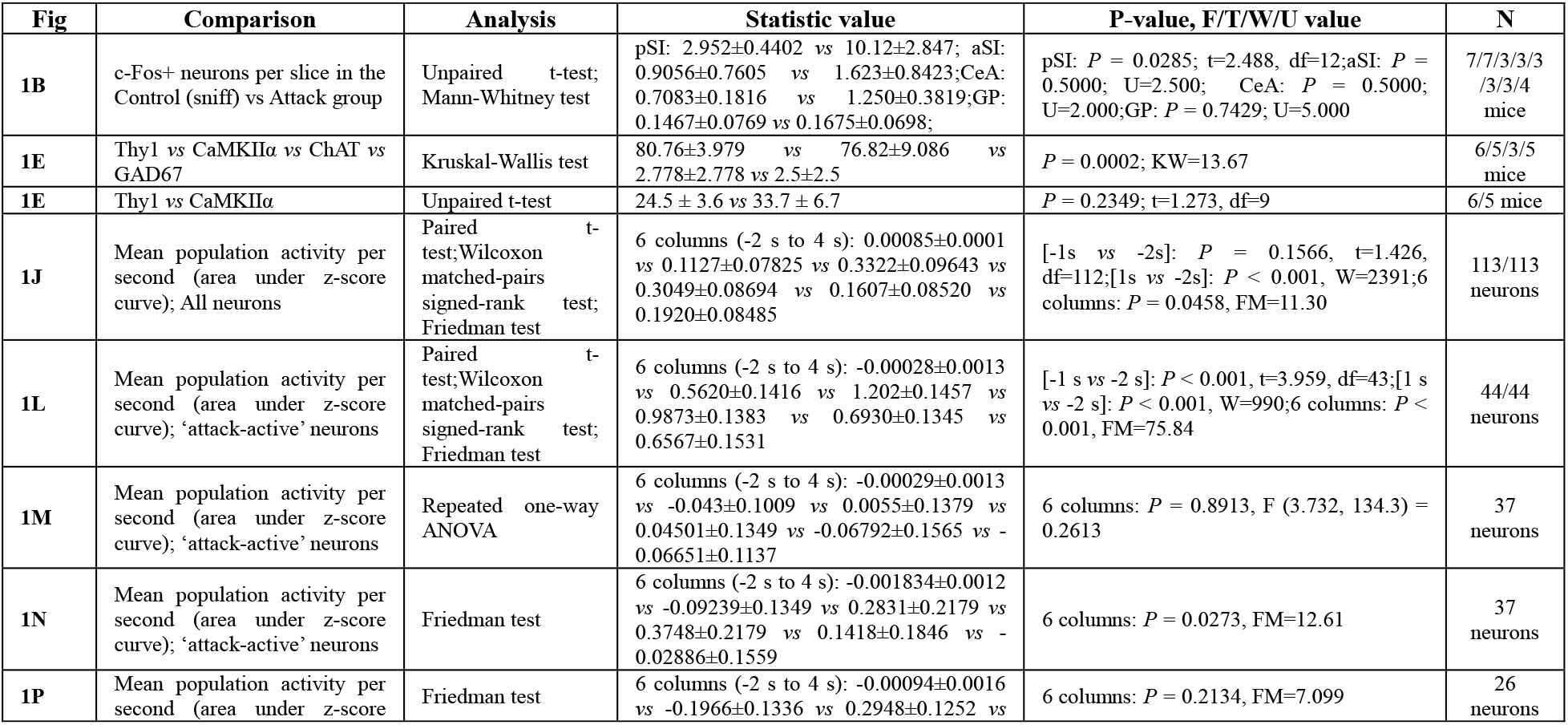
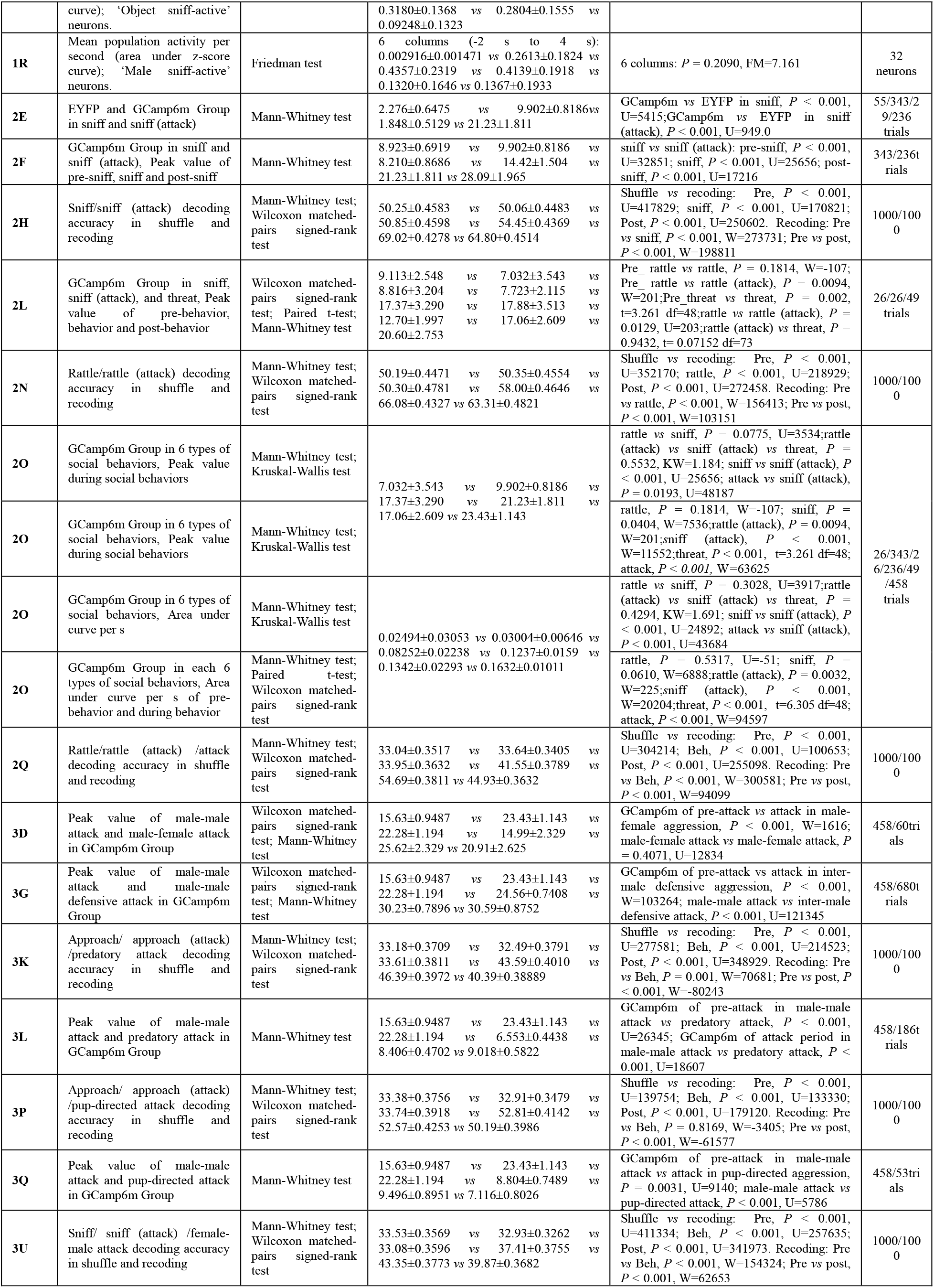
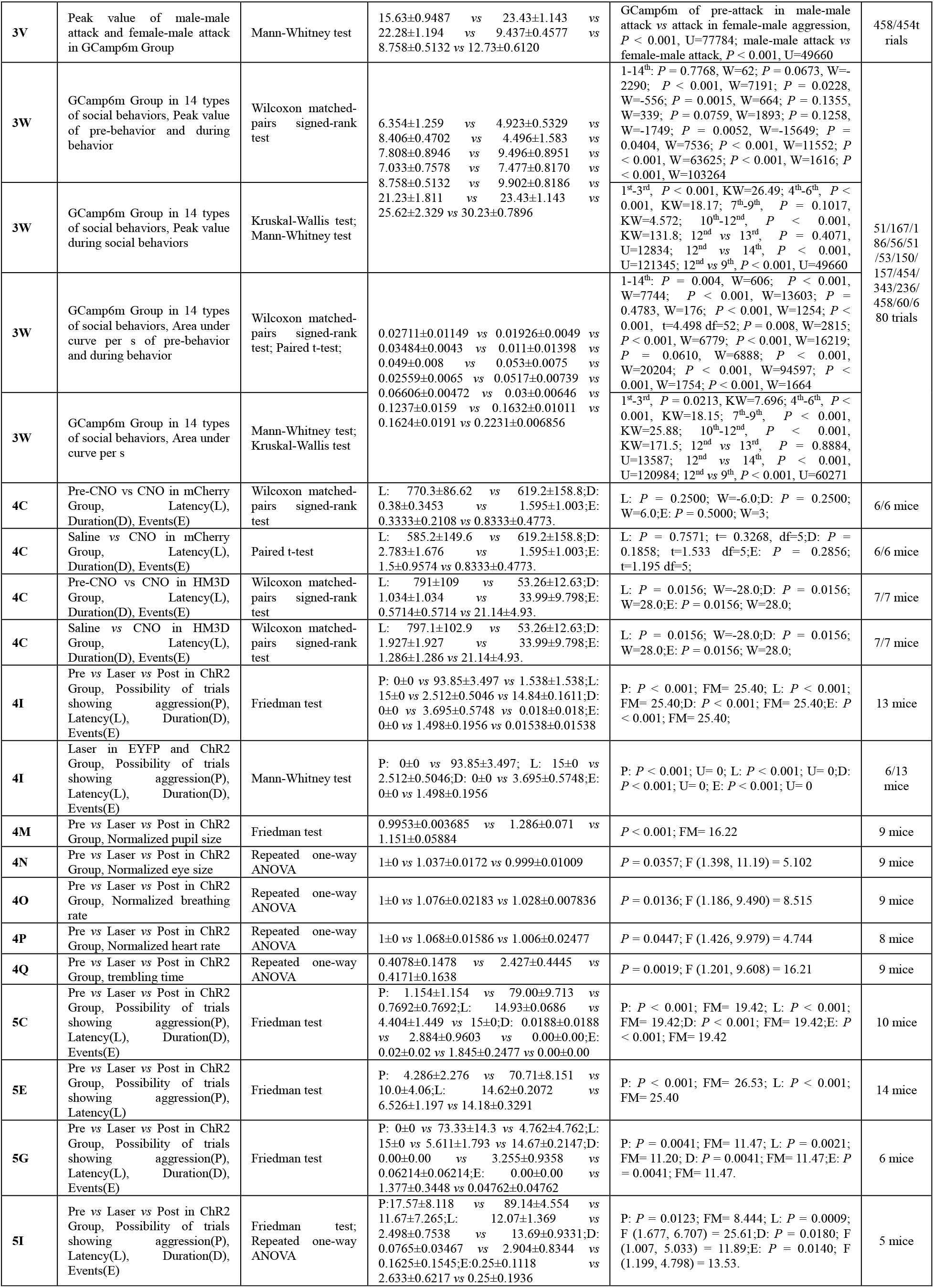
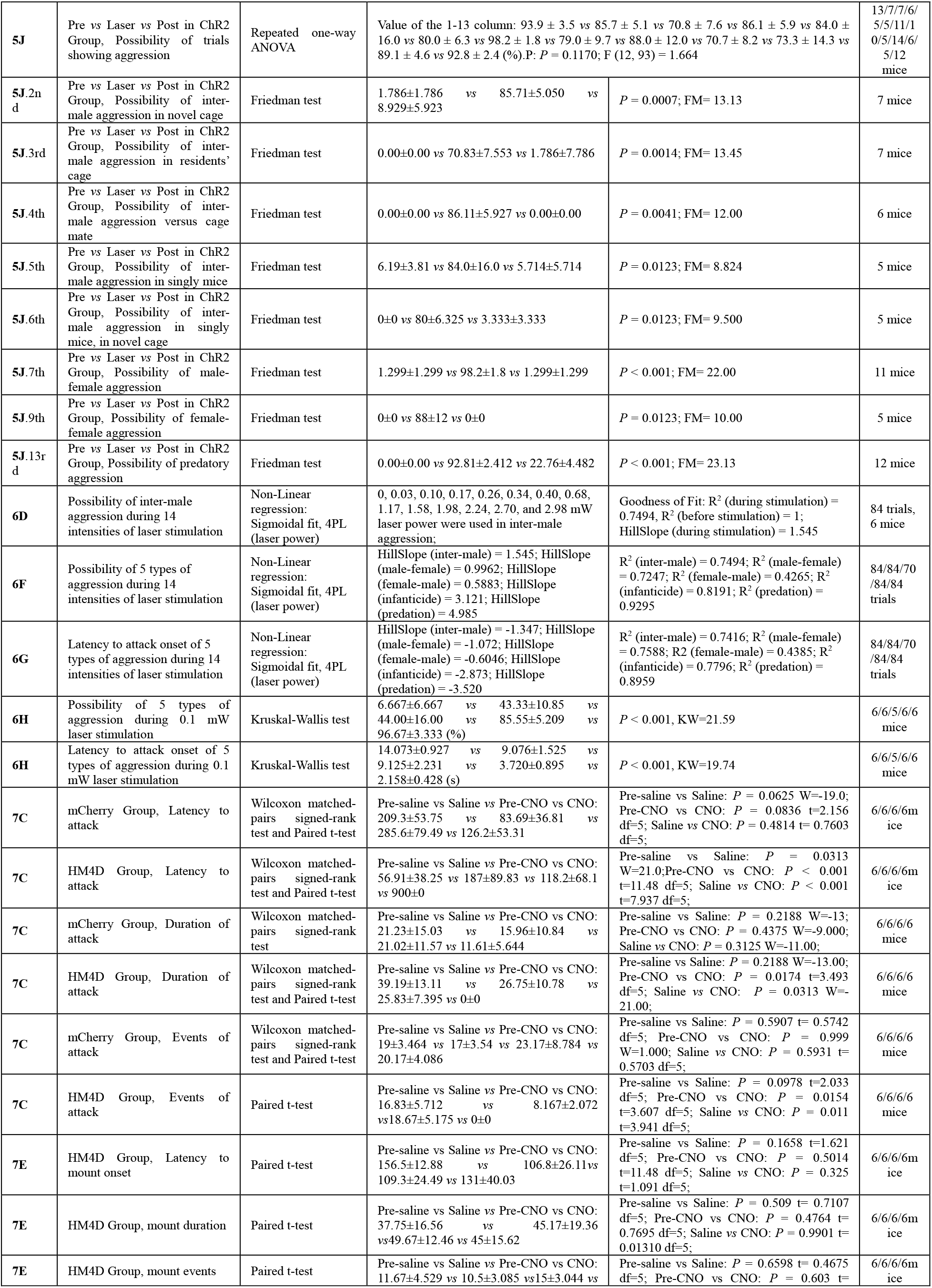
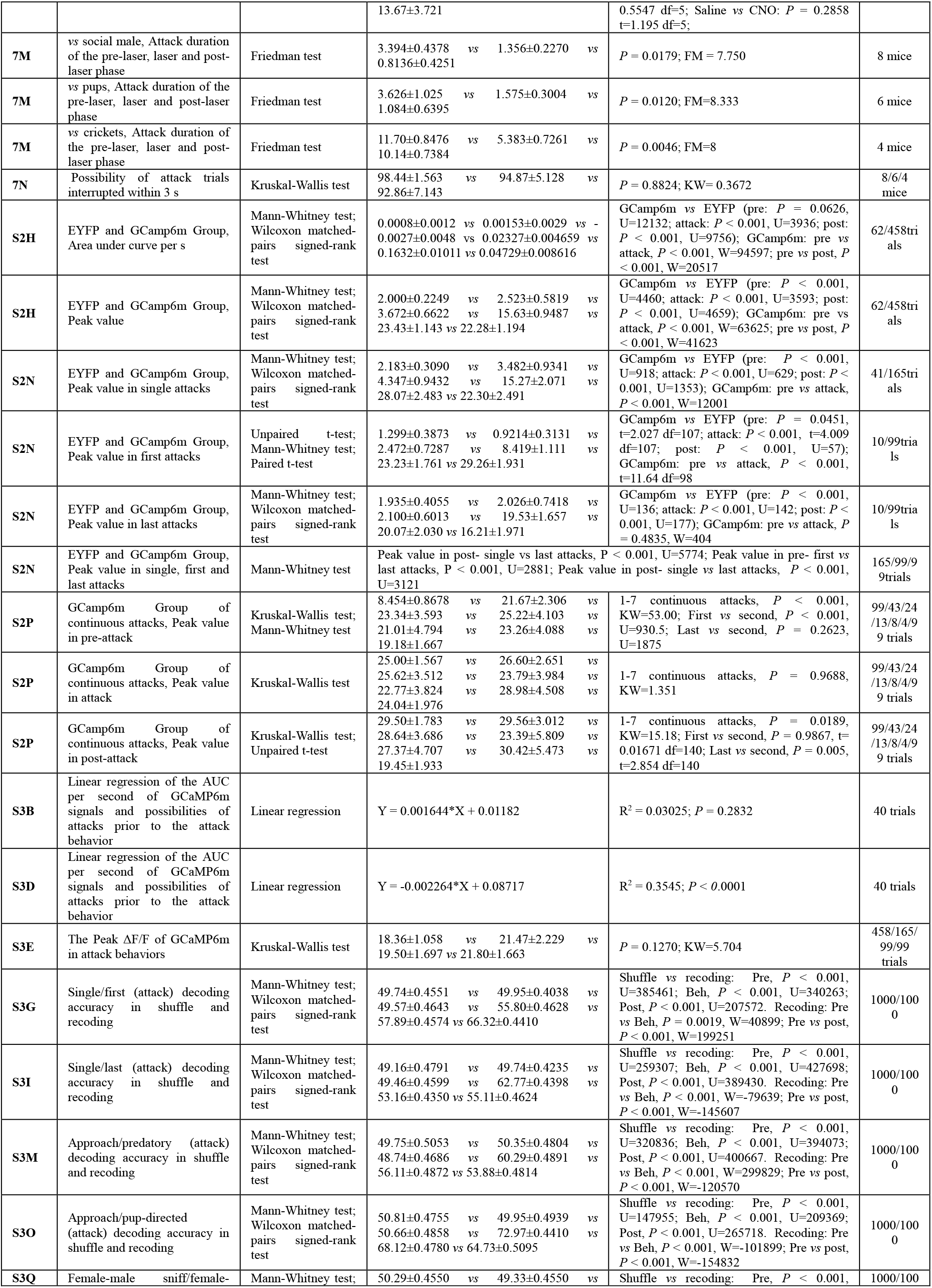
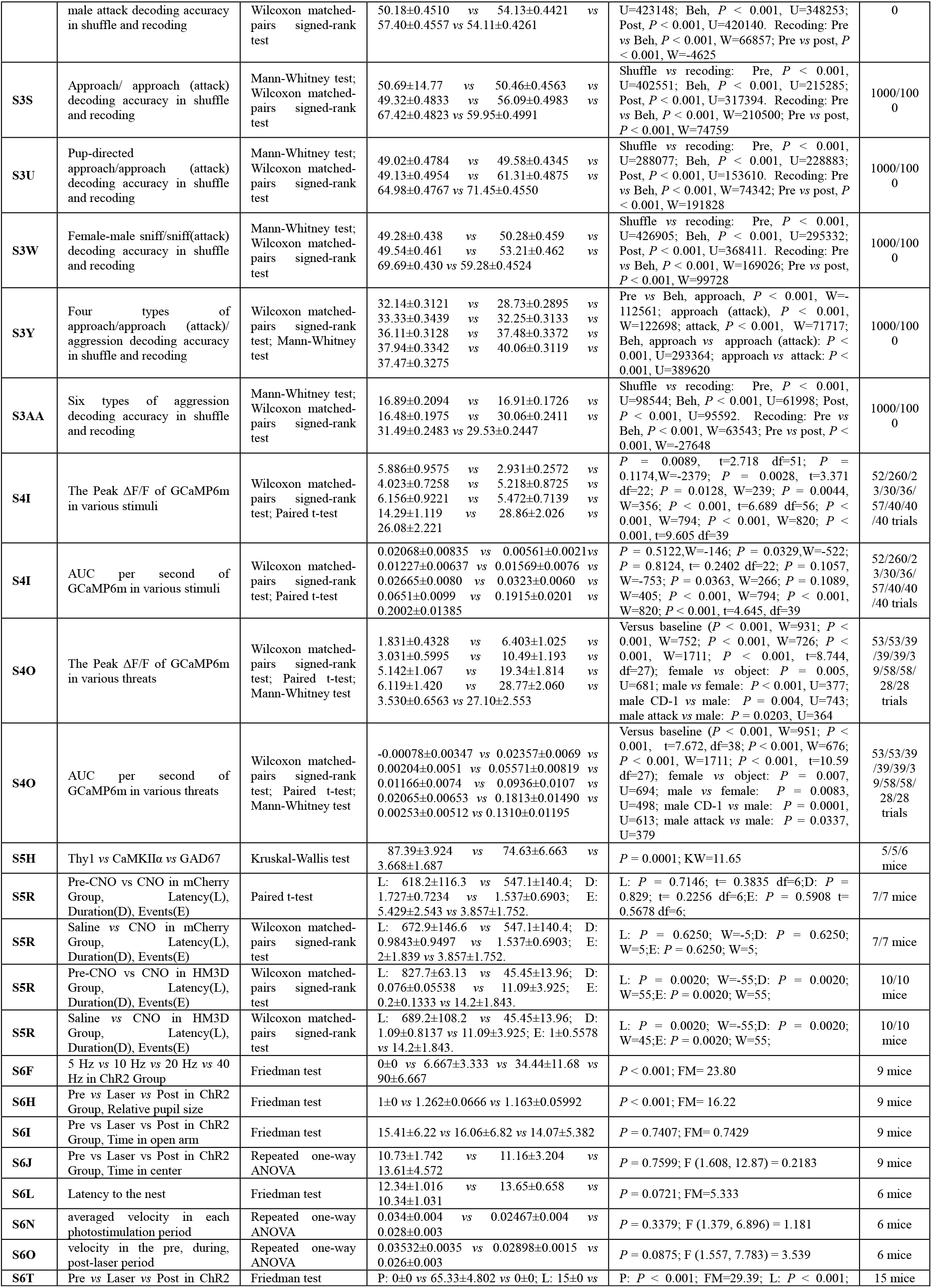
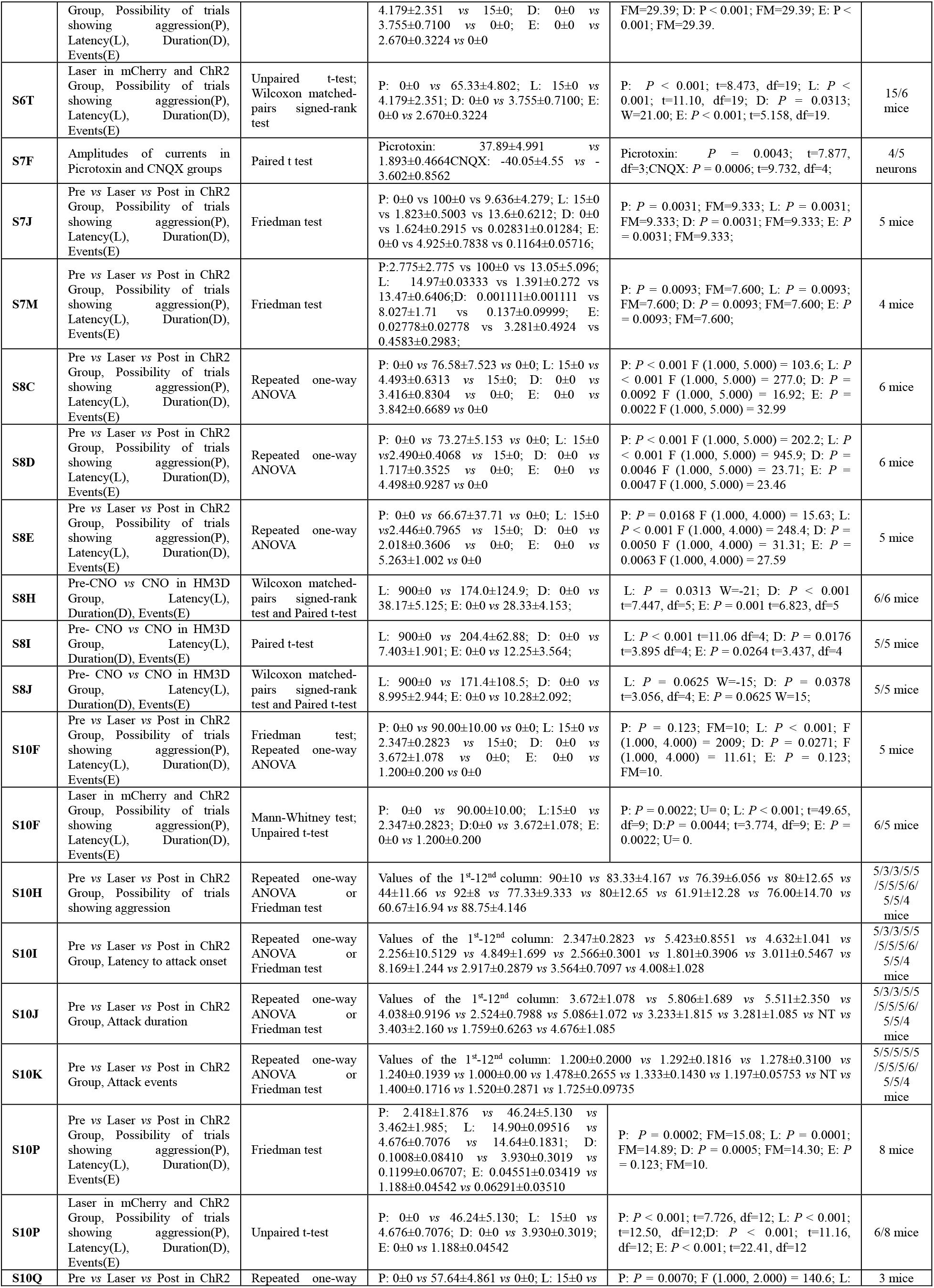
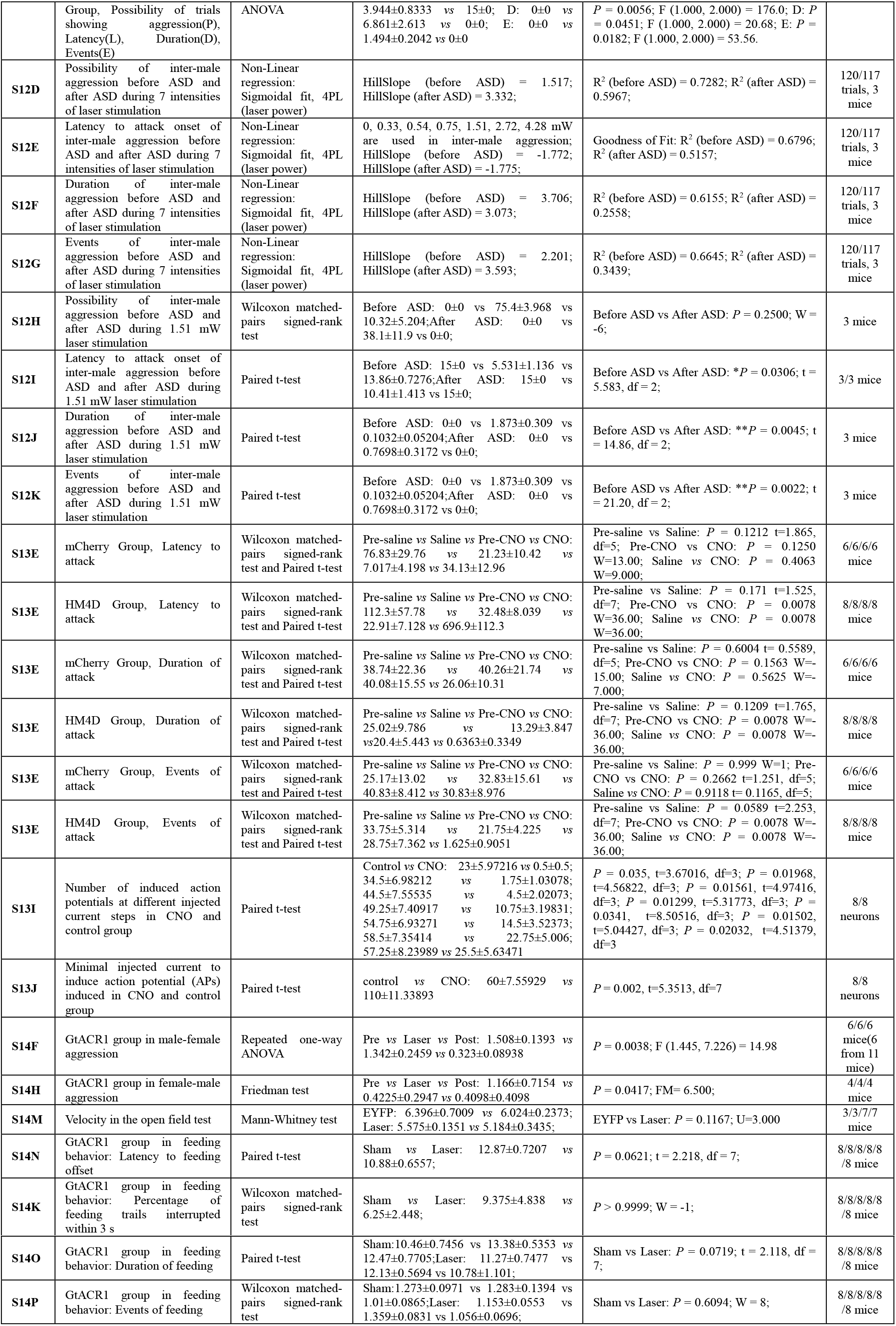
Statistical data for all Figures

**Table S2.** Functional manipulation studies of social behaviors from this paper

| Figures | Tested Mice(Strain, Sex,Social experience) | Conditions(Opponents, Environment) | Manipulation | Region | Cell-type | Attack | Mating |
| --- | --- | --- | --- | --- | --- | --- | --- |
| Figure4A-C | White socially adult virgin Thy1-Cre male mice (FVB/N) | Single adult males (C57); novel cage | Chemogenetic activation | pSI | pSI<sup>Thy1</sup> | Enhanced attack | No effect |
| Figure4F-I | Socially adult virgin males (C57) | Single adult males; home cage | Optogenetic activation | pSI | pSI<sup>PAG</sup> | Enhanced attack | No effect |
| Figure5B-C | Socially adult virgin females (C57) | Single adult males; home cage | Optogenetic activation | pSI | pSI<sup>PAG</sup> | Enhanced attack | No effect |
| Figure5D-E | Single adult sexually-experienced males (C57) | Pups; home cage | Optogenetic activation | pSI | pSI<sup>PAG</sup> | Enhanced attack | Not tested |
| Figure5F-G | Single adult virgin males (C57) | Moving gloves; home cage | Optogenetic activation | pSI | pSI<sup>PAG</sup> | Enhanced attack | Not tested |
| Figure5H-I | Socially adult virgin males (C57) | Single adult CD-1 males; CD-1's cage | Optogenetic activation | pSI | pSI<sup>PAG</sup> | Enhanced attack | No effect |
| Figure5J. 2nd | Socially adult virgin males (C57) | Single adult males; novel cage | Optogenetic activation | pSI | pSI<sup>PAG</sup> | Enhanced attack | No effect |
| Figure5J. 3rd | Socially adult virgin males (C57) | Single adult males; residents' cage | Optogenetic activation | pSI | pSI<sup>PAG</sup> | Enhanced attack | No effect |
| Figure5J. 4th | Socially adult virgin males (C57) | Single adult males, cage-mate; home cage | Chemogenetic activation | pSI | pSI<sup>PAG</sup> | Enhanced attack | No effect |
| Figure5J. 5th | Single adult virgin males (C57) | Single adult males; home cage | Optogenetic activation | pSI | pSI<sup>PAG</sup> | Enhanced attack | No effect |
| Figure5J. 6th | Single adult virgin males (C57) | Single adult males; novel cage | Optogenetic activation | pSI | pSI<sup>PAG</sup> | Enhanced attack | No effect |
| Figure5J. 7th | Socially adult virgin males (C57) | Single adult females; home cage | Optogenetic activation | pSI | pSI<sup>PAG</sup> | Enhanced attack | No effect |
| Figure5J. 9th | Socially adult virgin females (C57) | Single adult females; home cage | Optogenetic activation | pSI | pSI<sup>PAG</sup> | Enhanced attack | No effect |
| Figure5J.12<sup>nd</sup> | Socially housed males (C57) | Crickets; home cage | Optogenetic activation | pSI | pSI<sup>PAG</sup> | Enhanced attack | Not tested |
| Figure6 | Socially adult virgin males (C57) | Single adult males; home cage | Chemogenetic activation | pSI | pSI<sup>PAG</sup> | Enhanced attack | No effect |
| Figure6 | Socially adult virgin males (C57) | Single adult females; home cage | Optogenetic activation | pSI | pSI<sup>PAG</sup> | Enhanced attack | No effect |
| Figure6 | Socially adult virgin females (C57) | Single adult males; home cage | Optogenetic activation | pSI | pSI<sup>PAG</sup> | Enhanced attack | No effect |
| Figure6 | Single adult sexually-experienced males (C57) | Pups; home cage | Optogenetic activation | pSI | pSI<sup>PAG</sup> | Enhanced attack | Not tested |
| Figure6 | Socially housed males (C57) | Crickets; home cage | Optogenetic activation | pSI | pSI<sup>PAG</sup> | Enhanced attack | Not tested |
| Figure7A-C | Single adult Thy1-Cre males (FVB/N) | Socially adult males (C57); home cage | Chemogenetic inhibition | pSI | pSI<sup>Thy1</sup> | Reduced attack | Not tested |
| Figure7D-E | Socially adult sexually-experienced Thy1-Cre males (FVB/N) | Socially adult females (C57); home cage | Chemogenetic inhibition | pSI | pSI<sup>Thy1</sup> | Not tested | No effect |
| Figure7F-N | Single adult Thy1-Cre males (FVB/N) | Socially adult males (C57); home cage | Optogenetic inhibition | pSI | pSI<sup>Thy1</sup> | Reduced attack | Not tested |
| Figure7F-N | Single virgin adult Thy1-Cre males (FVB/N) | Pups; home cage | Optogenetic inhibition | pSI | pSI<sup>Thy1</sup> | Reduced attack | Not tested |
| Figure7F-N | Single adult Thy1-Cre males (FVB/N) | Crickets (5-10); home cage | Optogenetic inhibition | pSI | pSI<sup>Thy1</sup> | Reduced attack | Not tested |
| FigureS5 | Socially adult virgin males (C57) | Single adult males; novel cage | Chemogenetic activation | pSI | pSI<sup>PAG</sup> | Enhanced attack | No effect |
| FigureS6 | Socially adult virgin males (C57) | Single adult males; home cage | Optogenetic activation | pSI | pSI<sup>CaMKIIα</sup> | Enhanced attack | No effect |
| FigureS7 | Socially adult virgin Thy1-Cre males (FVB/N) | Socially adult males; home cage | Optogenetic activation | pSI | pSI<sup>CaMKIIα</sup> | Enhanced attack | No effect |
| FigureS7 | Socially adult virgin Thy1-Cre males (FVB/N) | Crickets; home cage | Optogenetic activation | pSI | PAG projecting pSI<sup>Thy1</sup> | Enhanced attack | Not tested |
| FigureS8 | Socially adult virgin males or females (C57) | Single adult males or females; home cage | Optogenetic activation | pSI | pSI<sup>CaMKIIα</sup> | Enhanced attack | No effect |
| FigureS8 | Socially adult virgin Thy1-Cre males or females (FVB/N) | Single adult males or females (C57); novel cage | Chemogenetic activation | pSI | pSI<sup>Thy1</sup> | Enhanced attack | No effect |
| FigureS10A-F | Socially adult virgin males (C57) | Single adult males; home cage | Optogenetic activation | PAG | pSI terminals in PAG | Enhanced attack | No effect |
| **FigureS10A-E** | Socially adult virgin males (C57) | Single adult males; home cage | Optogenetic activation | BNST | pSI terminals in BNST | No effect | No effect |
| **FigureS10A-E** | Socially adult virgin males (C57) | Single adult males; home cage | Optogenetic activation | VMH | pSI terminals in VMH | No effect | No effect |
| **FigureS10G-K** | Adult virgin males (C57); as for Fig. 5 | As for Fig. 5 | Optogenetic activation | PAG | pSI terminals in PAG | Enhanced attack | No effect |
| **FigureS10 L-R** | Socially adult virgin males or females (C57) | Single adult males; home cage | Optogenetic activation | PAG | pSI<sup>CaMKIIα</sup> terminals in PAG | Enhanced attack | No effect |
| **FigureS10 L-R** | Socially adult virgin males (C57) | Single adult males; home cage | Optogenetic activation | BNST | pSI<sup>CaMKIIα</sup> terminals in BNST | No effect | No effect |
| **FigureS12** | Single adult sexually-experienced males (C57) | Single adult males; home cage; later in CD-1's cage for ASD | Optogenetic activation | pSI | pSI<sup>PAG</sup> | Enhanced attack, be inhibited by ASD | Not tested |
| **FigureS13A-E** | Single adult sexually-experienced males (C57) | Socially adult males; home cage | Chemogenetic inhibition | pSI | pSI<sup>PAG</sup> | Reduced attack | Not tested |
| **FigureS14** | Single adult Thy1-Cre males (FVB/N) | Socially adult females (C57); home cage | Optogenetic inhibition | pSI | pSI<sup>Thy1</sup> | Reduced attack | Not tested |
| **FigureS14** | Single adult Thy1-Cre females (FVB/N) | Socially adult males (C57); home cage | Optogenetic inhibition | pSI | pSI<sup>Thy1</sup> | Reduced attack | Not tested |

'Enhanced attack' or 'reduced attack' indicates that the aggressive behavior during manipulation was significantly changed; 'No effect' indicates aggressive behavior during manipulation was not observed or was not significantly changed.

## Notes

### Competing Interest Statement

The authors have declared no competing interest.

